# Bacteroidetes contribute to the carbon and nutrient cycling of deep sea through breaking down diverse glycans

**DOI:** 10.1101/2020.11.07.372516

**Authors:** Rikuan Zheng, Ruining Cai, Rui Liu, Ge Liu, Chaomin Sun

**Affiliations:** CAS Key Laboratory of Experimental Marine Biology & Center of Deep Sea Research, Institute of Oceanology, Chinese Academy of Sciences, Qingdao, China; Laboratory for Marine Biology and Biotechnology, Qingdao National Laboratory for Marine Science and Technology, Qingdao, China; College of Earth Science, University of Chinese Academy of Sciences, Beijing, China; Center of Ocean Mega-Science, Chinese Academy of Sciences, Qingdao, China

## Abstract

Bacteroidetes are thought to be specialized for the degradation of algae-derived ocean polysaccharides and are a major contributor to the marine carbon and nutrient cycling. Here, we first show Bacteroidetes are the second most abundant phylum bacteria in deep-sea cold seep and possess more genes associated with polysaccharides degradation than other bacteria through metagenomics methods. We further isolate a novel Bacteroidetes species, *Maribellus comscasis* WC007^T^, which can efficiently degrade numerous different polysaccharides including: cellulose, pectin, fucoidan, mannan, xylan and starch. These results are verified by transcriptomic analyses and growth assays. Notably, we find cellulose promotes abundant bacterial growth, and using transcriptomics and metabolomics we finally report on the underlying mechanisms of cellulose degradation and utilization, as well as potential contributions to the carbon cycling. Overall, our results suggest Bacteroidetes play key roles in the deep-sea carbon and nutrient cycling, likely due to their high abundance and prominent polysaccharide degradation capabilities.

**One Sentence Summary:** Bacteroidetes contribute to ocean carbon and nutrient cycle.

## Introduction

Roughly half of the global net primary biomass production is oceanic and carried out predominantly by small marine phytoplankton. As the primary biomass production, complex carbohydrates are a ubiquitous energy source for microorganisms in both terrestrial and marine ecosystems (**Grondin et al., 2017**). Complex carbohydrates, mostly in the form of polysaccharides, are the largest repository of organic carbon in the biosphere (**Malhi, 2002**). In the ocean, polysaccharides constitute a substantial fraction of biomass and marine algae are a dominant source of polysaccharides (**Benner et al., 1992; Engel et al., 2004**). Marine algae are likely composed of greater than 50% polysaccharides, which are found as structural components of cell walls and intracellular energy storage compounds (**Kloareg Quatrano, 1988**). The polysaccharides produced at the ocean surface can sink to a lower water column and even the deep sediments. With this, large amounts of mostly algal plants and animal debris are deposited from the upper ocean, providing an important organic carbon source for polysaccharide-degrading bacteria in deep-sea sediments (**Gao et al., 2017**). Algal debris is rich in difficult-to-degrade polysaccharides, such as pectin, cellulose and hemicellulose (including fucoidan, mannan, xylan, glucan, arabinogalactan and others) (**Snajdr et al., 2011**). These polysaccharides are the main components of the cell wall and intercellular layer. Breakdown of these polysaccharides by heterotrophic microorganisms plays a significant role in organic biomass degradation and greatly contributes to the marine carbon and nutrient cycling (**Azam, 1998; Moran et al., 2016**). Likewise, polysaccharides are a major nutrient source for degradation microorganisms (**Azam Malfatti, 2007**), providing these organisms with a significant growth advantage and allowing them to efficiently occupy niches in harsh deep-sea environments.

Notably, many members of the bacterial phylum Bacteroidetes are specialized in polysaccharide degradation and are thought to be the most abundant group of ocean bacteria after Proteobacteria and Cyanobacteria (**Kirchman, 2002**). Bacteroidetes are globally distributed in coastal areas (**Pommier et al., 2007**), sediment regions, hydrothermal vents and numerous other marine environments (**Alonso et al., 2007; Teeling et al., 2012**). Marine Bacteroidetes are commonly thought to play a key role in degrading phytoplankton polysaccharides (**Fernandez-Gomez et al., 2013**), likely owing to the number and diversity of carbohydrate-active enzymes (CAZymes) in their genomes (**Hahnke et al., 2016**). Bacteroidetes CAZymes are categorized into families of glycoside hydrolases (GHs), glycoside transferases (GTs), carbohydrate-binding modules (CBMs), carbohydrate esterases (CEs), polysaccharide lyases (PLs), sulfatases and a broad range of auxiliary enzymes (**Bauer et al., 2006; Cantarel et al., 2009; Fernandez-Gomez et al., 2013**). As such, Bacteroidetes have evolved unique and sophisticated degradation systems, typically called polysaccharide utilization loci (PULs). These loci are tightly regulated and are comprised of colocalized gene clusters that encode CAZymes and the protein assemblies required for the degradation of complex carbohydrates. In addition to substrate-specific CAZymes genes, Bacteroidetes PULs also contain genes encoding TonB-dependent transporters (TBDTs), which are SusCD-like extracellular lipoproteins with integral membrane beta-barrels (**Kappelmann et al., 2019**). Polysaccharides initially bind to outer membrane proteins and are cleaved into oligosaccharides by endo-active enzymes (**Kappelmann et al., 2019**). Oligosaccharides are transported from the outer membrane into the periplasm in a TonB-dependent manner. In the periplasm, oligosaccharides are protected from competition by other bacteria and are further degraded into monosaccharides, which are then moved by dedicated transporters across the cytoplasmic membrane into the cytoplasm for utilization (**Glenwright et al., 2017**).

Various mechanisms of Bacteroides polysaccharide degradation have been investigated in the human gut, including the degradation of pectin and hemicellulose (**Larsbrink et al., 2014; Rogowski et al., 2015**). Intriguingly, a study suggests human gut bacteria could acquire CAZymes encoding genes from marine bacteria, which may be a general factor of CAZyme diversity in human gut microbes (**Hehemann et al., 2010**). However, as compared to human gut Bacteroides, few studies have investigated polysaccharide degradation by marine Bacteroides, especially the deep-sea variety.

Here we first investigate the abundance of Bacteroidetes in the deep-sea cold seep by metagenomics methods. We successfully isolate a novel Bacteroidetes strain *Maribellus comscasis* WC007^T^ by a goal-directed method from the deep-sea sediments. Strain WC007^T^ efficiently degrades various polysaccharides. Using genomics, transcriptomics and metabolomics, we deeply characterize the degradation and utilization of different polysaccharides by *M. comscasis* WC007^T^. Lastly, we discuss the contribution of Bacteroidetes to the deep ocean carbon and nutrient cycling.

## Results

### Bacteroides are high abundance bacteria in deep-sea cold seep

To gain insights into the bacterial community composition of the deep sea, we performed OTUs sequencing analysis of bacteria present in cold seep sediments. Our results indicated that, at the phylum level, Proteobacteria was the most abundant domain (68.8%), followed by Bacteroidetes (13.2%), Chloroflexi (4.5%), Planctomycetes (4.0%), Firmicutes (1.9%), Acidobacteria (1.7%), Lateseibacteria (1.4%) and Spirochaetes (1.1%). Together, these constituted roughly 96.6% of all bacteria in the sampled cold seep. The remaining 3.4% was other phyla bacteria (Figure 1A). Notably, Bacteroidetes were the second most abundant phylum, suggesting Bacteroidetes are dominant in these deep-sea regions. To understand the ability of deep-sea Bacteroidetes to degrade polysaccharides, we used metagenome sequencing to predict and enumerate genes encoding CAZymes. Fifty-one high quality metagenome-assembled genomes (MAGs) for the three most abundant phyla (Proteobacteria, Bacteroidetes and Chloroflexi) were obtained by using a hybrid binning strategy combined with manual inspection and data curation (Supplementary file 2). Through the analysis of 51 MAGs, we found that Bacteroidetes had the highest abundances of CAZymes, followed by Chloroflexi and Proteobacteria (Figure 1B). We also analyzed the number of genes encoding CAZymes, SusC, SusD and sulfatase proteins in our Bacteroidetes MAGs. As expected, all 8 MAGs were rich in CAZymes, SusC, SusD and sulfatase genes (Figure 1C, Supplementary file 3), as these genes were necessary components for polysaccharide degradation.

**Figure 1.**
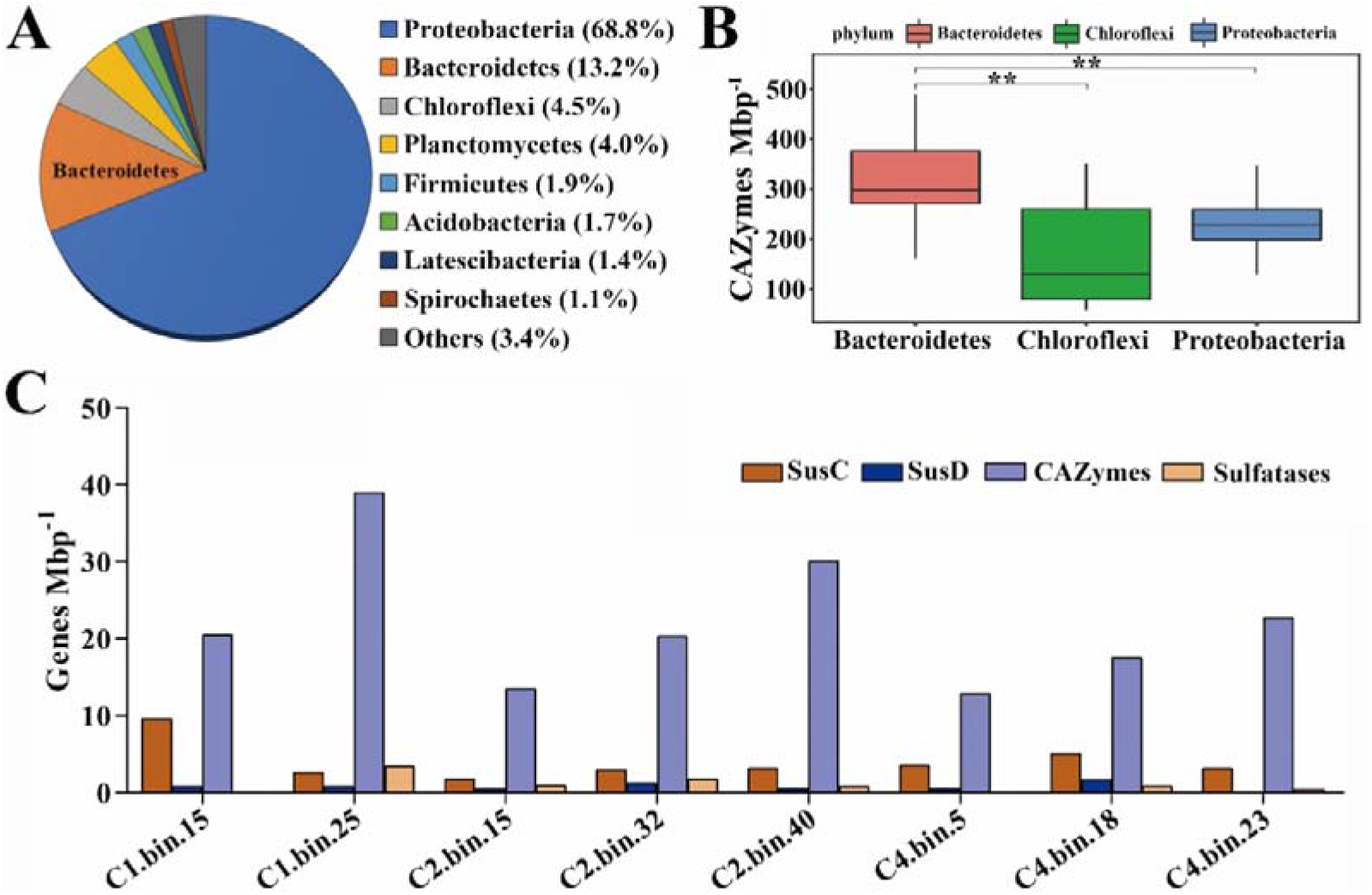
Quantification and polysaccharide degradation predictions of Bacteroides derived from deep-sea cold seep sediments. (**A**) 16S rRNA amplicon-based assessment of bacterial abundance in the deep-sea cold seep sediments. (**B**) Calculated number of genes associated with polysaccharide degradation from metagenome-assembled genomes of Bacteroides, Chloroflexi and Proteobacteria derived from deep-sea cold seep. (**C**) Calculated number of genes encoding SusC, SusD, CAZymes and sulfatases identified from 8 metagenome-assembled genomes of Bacteroides derived from deep-sea cold seep. X-axis indicates the names of Bacteroides metagenome-assembled genomes.

To further understand the types of polysaccharides degraded by deep-sea Bacteroidetes, we conducted in-depth analysis of GHs obtained from above 8 MAGs. The results showed that the major GH families of Bacteroidetes were GH2 (a family of diverse functions), GH3 (also a family of diverse functions), GH29 (a family of alpha-L-fucosidases) and GH92 (a family of mostly alpha-mannosidases) (Figure 1-figure supplement 1). This suggests Bacteroidetes may be important participants of diverse polysaccharides degradation in the deep-sea cold seep environment.

### Isolation and characterization of a new deep-sea Bacteroides species,*Maribellus comscasis* WC007^T^, that has prominent polysaccharide degradation functions

Given their high abundance and potential polysaccharide degradation functions, we isolated deep-sea derived Bacteroides to investigate novel metabolic pathways of polysaccharide degradation. Considering the superior polysaccharide degradation capability of Bacteroides, the deep-sea sediment sample was anaerobically enriched at 28 °C for one month in an inorganic medium supplemented with various polysaccharides (Figure 2A). Enriched samples were then plated on solid medium in Hungate tubes and individual colonies with distinct morphology were picked and cultured (Figure 2A). As expected, most cultured colonies were identified as Bacteroides. Among them, strain WC007^T^ grew significantly faster than other Bacteroides strains and was chosen for further study.

**Figure 2.**
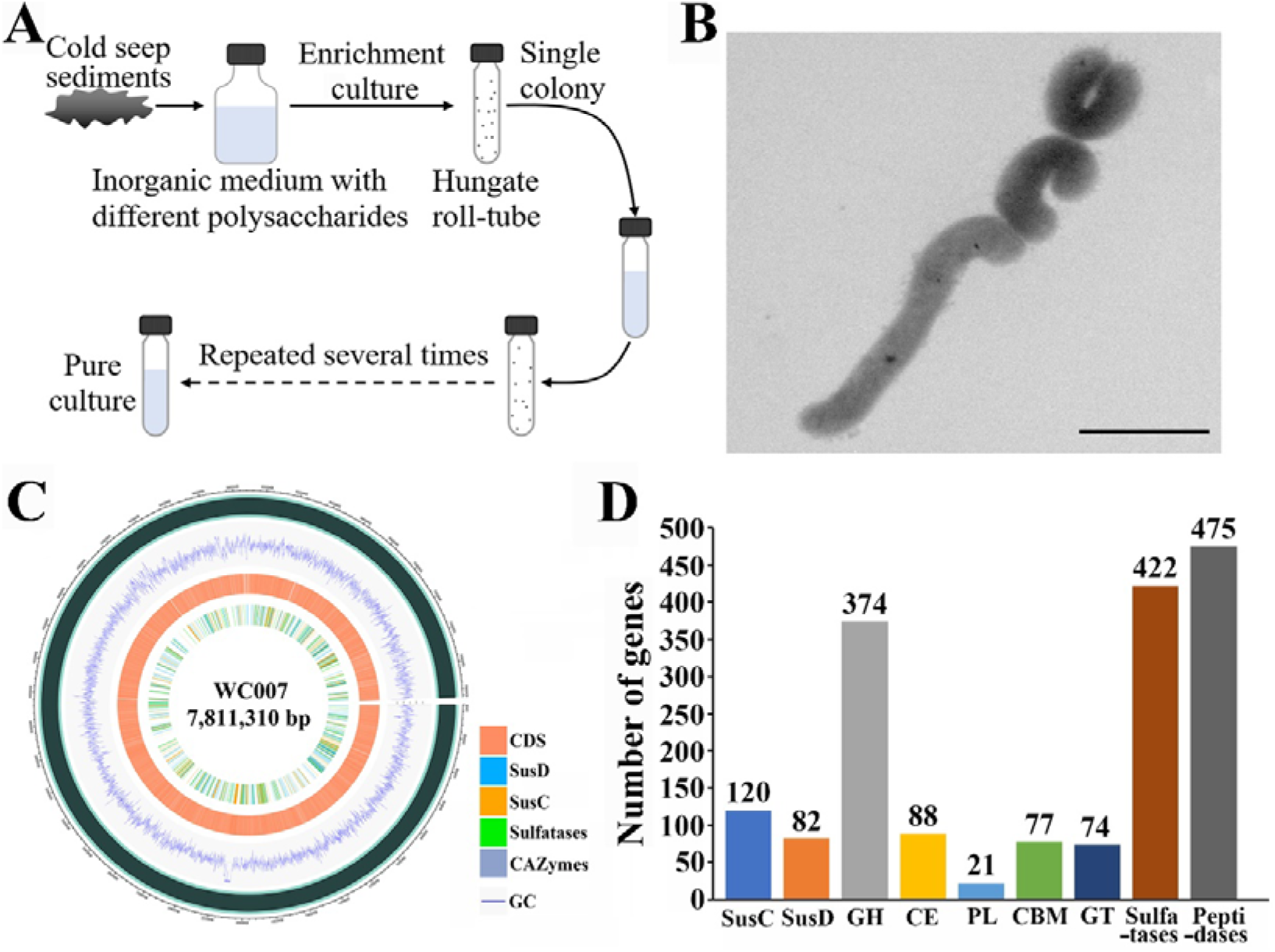
Goal-directed isolation of a novel deep-sea Bacteroides species possessing strong polysaccharide degradation capabilities. (**A**) Diagrammatic scheme for enrichment and isolation of deep-sea Bacteroides. (**B**) TEM observation of *Maribellus comscasis* WC007^T^, bar = 2 μm. (**C**) Genomic distribution of SusC, SusD, CAZymes, sulfatases genes in *M. comscasis* WC007^T^. (**D**) Quantification of SusC, SusD, CAZymes, sulfatases and peptidases genes in the *M. comscasis* WC007^T^ genome.

Cells of strain WC007^T^ were curved, approximately 2.0-4.0 × 0.5-0.8 μm in size, and had no flagellum, as indicated by TEM (Figure 2B). The whole genome size of strain WC007^T^ was 7,811,310 bp with a DNA G+C content of 38.38%. A sequence similarity calculation using the NCBI server indicated the closest relatives of strain WC007^T^ were *Maribellus luteus* XSD2^T^ (95.70%), *Mariniphaga sediminis* SY21^T^ (93.44%) and *Draconibacterium sediminis* MCCC 1A00734^T^ (92.99%). Phylogenetic analyses based on the 16S rRNA and genome sequences showed strain WC007^T^ belonged to the genus *Maribellus* and formed an independent phyletic line with the type strain *Maribellus luteus* XSD2^T^ (Figure 2-figure supplement 2 and 3). Therefore, we propose strain WC007^T^ to be classified as the type strain of a novel species in the genus *Maribellus*, for which the name *Maribellus comscasis* sp. nov. is proposed.

Details of the phenotypic, chemotaxonomic, and genotypic characterizations of WC007^T^ were shown in Supplementary files 4-6. When compared to its closest relative *Maribellus luteus* XSD2^T^, strain WC007^T^ showed a distinct capability for degrading cellulose, pectin and xylan (Supplementary file 5), indicating its potential for polysaccharide degradation. Consistently, genes encoding SusC, SusD, CAZymes and sulfatase were ubiquitously distributed in the WC007^T^ genome (Figure 2C). Moreover, detailed analysis with the Pfam, TIGRFAM, dbCAN, MEROPS and CAZyme databases showed that strain WC007^T^ had 634 CAZymes (including 374 GHs, 88 CEs, 21 PLs, 77 CBMs and 74 GTs), 120 SusC-like proteins, 82 SusD-like proteins, 422 sulfatases and 475 peptidases (Figure 2D). The numbers of genes encoding CAZymes, SusC/SusD pairs and sulfatases in WC007^T^ were significantly higher than those of other Bacteroides from different habitats including human gut, lake, shallow ocean and deep sea (Figure 2-figure supplement 4 and Supplementary file 7). Overall, these data suggest WC007^T^ is a bacterium with a notable polysaccharide degradation capability. Studies focusing on this strain may provide a more comprehensive understanding of carbon cycling in the deep sea.

### Prediction and validation of polysaccharides degradation by*M. comscasis* WC007T

To gain further insights into the potential mechanisms of polysaccharide degradation by *M. comscasis* WC007^T^, we conducted an in-depth analysis of the CAZymes, Sus transport genes and sulfatases present in the WC007^T^ genome. In previous studies (**Kabisch et al., 2014; Kappelmann et al., 2019; Lapebie et al., 2019**), polysaccharide utilization loci (PULs) are defined as the presence of CAZymes with predicted polysaccharide degradation function and a gene pair encoding SusC/D-like proteins (**Sonnenburg et al., 2010**). We identified 82 PUL-containing gene clusters with a total of 82 unique *susC/D* gene pairs (Figure 3A, Figure 3-figure supplement 1–6, Supplementary file 8). Every PUL carried one *susC/D* pair and one or more CAZyme encoding gene(s). Of these 82 PULs, 44 were linked to either dedicated polysaccharide or polysaccharide classes according to in-depth annotations. The remaining 38 PULs could not be definitively inferred. Surprisingly, 58 of these PULs comprised sigma-70/anti-sigma factors, a rare observation in Bacteroidetes PULs, suggesting potentially novel regulation pathways exist in deep-sea Bacteroidetes. Substrate predictions suggested these PULs likely degraded six types of polysaccharides including: cellulose, pectin, fucoidan, mannan, xylan and starch (Figure 3A and Supplementary file 9). These were further described in the Supplementary results.

**Figure 3.**
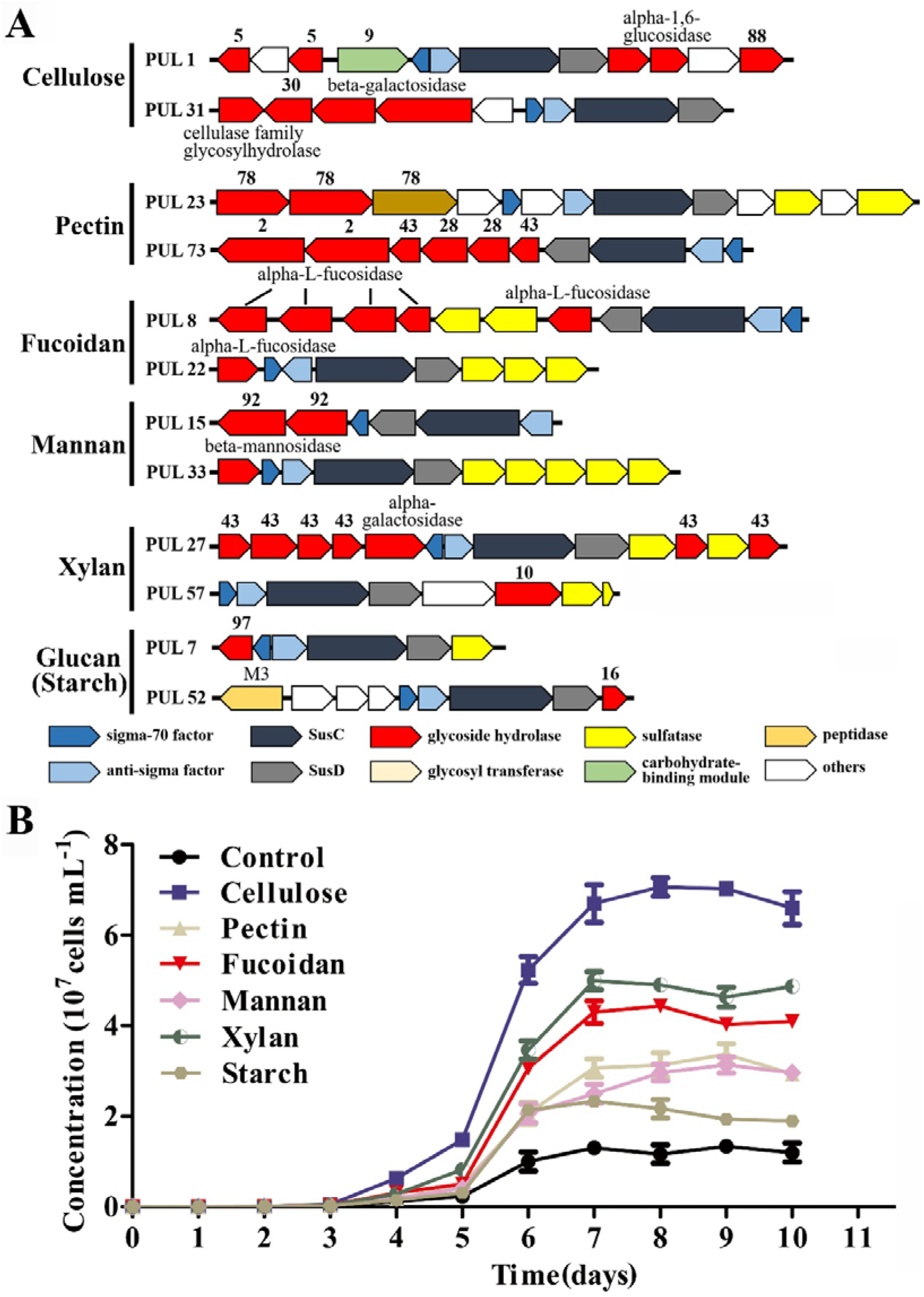
Prediction and verification of polysaccharide degradation and utilization by *M. comscasis* WC007^T^. (**A**) Selected putative PULs responsible for degradation of cellulose, pectin, fucoidan, mannan, xylan and starch polysaccharides. Genes encoding CAZymes, SusC/SusD, sigma factor, anti-sigma factor, sulfatase and peptidase are depicted in different colors. Hypothetical proteins and genes involved in other metabolic functions are depicted in white. (**B**) Growth assays of *M. comscasis* WC007^T^ in inorganic medium supplemented with cellulose, pectin, fucoidan, mannan, xylan or starch polysaccharides (all at a final concentration of 1 g/L).

To verify the predicted polysaccharide substrate spectra of *M. comscasis* WC007^T^, we grew WC007^T^ in medium containing various polysaccharides as the sole organic carbon source. Unsurprisingly, all six polysaccharides (cellulose, pectin, fucoidan, mannan, xylan and starch) promoted growth of WC007^T^ (Figure 3B). However, cellulose had an almost 7-fold increased growth compared to the control group (Figure 3B). From these findings we conclude that *M. comscasis* WC007^T^ possesses versatile polysaccharide degradation capabilities.

### Transcriptomic profiling of polysaccharide degradation and utilization by *M. comscasis* WC007^T^

To further validate the roles of predicted PULs and polysaccharides degradation, we performed transcriptomic analysis of *M. comscasis* WC007^T^ during growth in medium containing different polysaccharides. For consistency, we focused on significantly up-regulated PULs and expression changes of genes encoding CAZymes, SusC/SusD, sigma-70 factor and anti-sigma factor within the corresponding PULs. We found, respectively, there were 2, 4, 5, 1, 4, 3 up-regulated PULs in cellulose (Figure 4-figure supplement 1), pectin (Figure 4-figure supplement 2), fucoidan (Figure 4-figure supplement 3), mannan (Figure 4-figure supplement 4), xylan (Figure 4-figure supplement 5) and starch (Figure 4-figure supplement 6) treatment groups. PUL 29 was expressed in three polysaccharides treatment groups (pectin, fucoidan and starch), PUL 54 was expressed in both pectin and starch treatment groups, PUL 65 was expressed in both pectin and xylan treatment groups, and the remaining PULs were only presented in a single polysaccharide treatment group. It is worth noting that only four PULs were consistent with predicted and detected activities. These included PUL 73 (pectin degradation), PUL 26 (mannan degradation), PUL 27 and PUL 76 (xylan degradation). These results suggest a single PUL might function in degrading different substrates. In future studies, it will be important to clarify the specific activity of each PUL in the genome of WC007^T^.

To further characterize the dynamic changes of individual genes within WC007^T^ PULs under different treatment conditions, we analyzed the glycoside hydrolases, which were up-regulated simultaneously in three or more polysaccharides treatment groups. Genes encoding GH2, GH3, GH20, GH27, GH28, GH43, GH92 and GH95 family enzymes were selected (Figure 4A), as they are proposed to function in different polysaccharides degradation functions. We also analyzed some annotated glycoside hydrolases outside predicted PULs. We found that expression of the same GH could be induced by different polysaccharides, such as alpha-L-fucosidase (GM_418 30400), GH43 (GM_418 28935) and glycoside hydrolase (GM_418 21515). Additionally, glycoside hydrolase (GM_418 21515), whose expression was significantly up-regulated by all six polysaccharides, may play a key role in mediating polysaccharides degradation of WC007^T^. Future studies should help determine the exact function of this hydrolase. We also observed upregulation of genes encoding glycoside hydrolases outside predicted PULs, suggesting cryptic PULs may exist in WC007^T^ and may aid in the breakdown of glycans. Upon analysis of up-regulated genes encoding SusC/SusD and sigma-70 factor/anti-sigma factor located in the same PUL, we found 8 PULs were simultaneously induced in 3 or more polysaccharides treated groups (Figure 4B). The genes encoding sigma-70 factor (GM418_00730) and SusC (GM418_09255) were consistently up-regulated by 6 different polysaccharides, indicating their universal functions in polysaccharide degradation.

**Figure 4.**
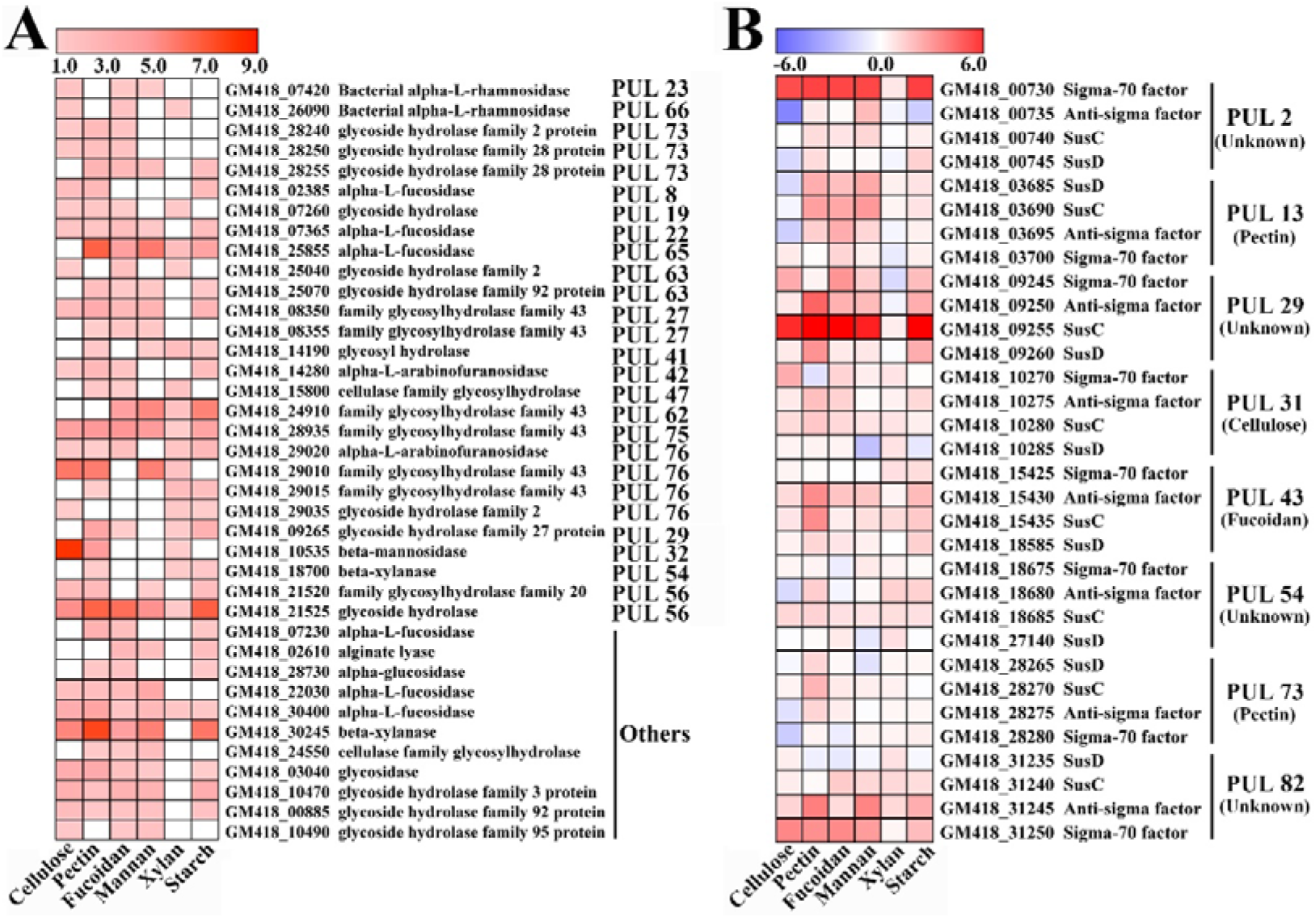
Transcriptomic analysis of *M. comscasis* WC007^T^ cultured in inorganic medium supplemented with different polysaccharides. Heat map showing differentially expressed genes encoding glycoside hydrolase genes (**A**) and SusC, SusD, sigma-70 factor and anti-sigma factor (**B**) in *M. comscasis* WC007^T^ supplemented with 1 g/L of different polysaccharides. Genes whose expression was up-regulated in three or more polysaccharides treatment groups are shown. The heat map is generated by the Heml 1.0.3.3 software.

### Metabolomics analysis of cellulose degradation by *M. comscasis* WC007^T^

Of the 6 degradation polysaccharides, cellulose promoted the most growth of *M. comscasis* WC007^T^ (Figure 3B). From this, we further investigated the degradation and utilization capabilities of cellulose by *M. comscasis* WC007^T^. We first observed degradation of cellulose by *M. comscasis* WC007^T^ by SEM. In the control group the morphological surface of cellulose was uniform and intact (Figure 5A); however, in the *M. comscasis* WC007^T^ group, cellulose was clearly degraded into small pieces, and a large amount of bacterial cells were attached to and even embedded in cellulose particles (Figure 5B-D). Transcriptomic analysis also showed PUL 32 was significantly up-regulated (up to 256-fold) when the bacterium was incubated in the medium supplemented with cellulose (Figure 5E). Additionally, all of genes within PUL 32 were significantly up-regulated, as detected by qRT-PCR (Figure 5F). Overall, our above results suggest that *M. comscasis* WC007^T^ can efficiently degrade cellulose for cellular energy and growth.

**Figure 5.**
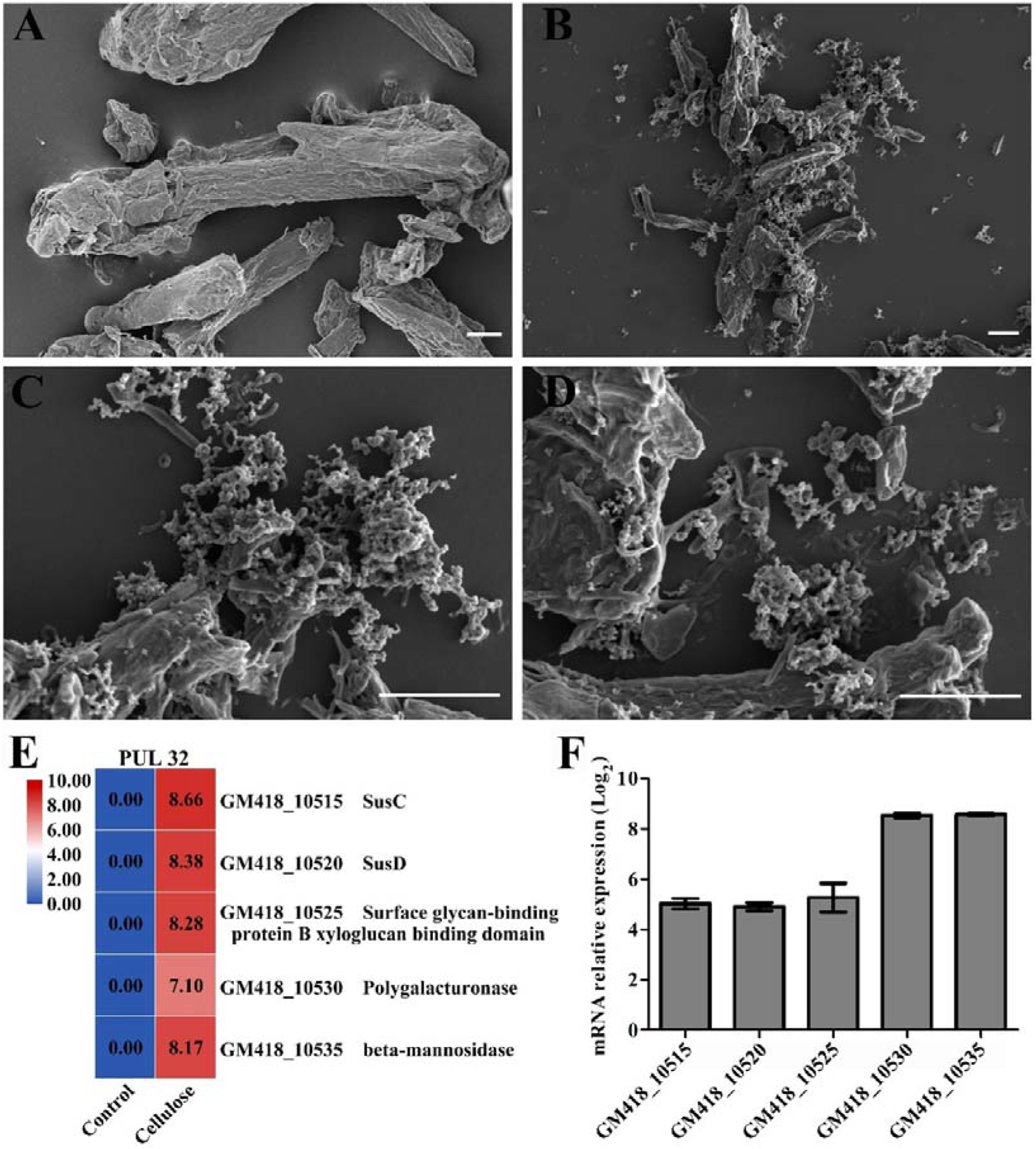
Characterization of cellulose degradation by *M. comscasis* WC007^T^. (**A**) SEM observation of cellulose. (**B-D**) SEM observation of cellulose (1 g/L) degradation by *M. comscasis* WC007^T^. Scale bar is 10 μm. (**E**) PUL 32 genes are significantly up-regulated when culturing *M. comscasis* WC007^T^ in the presence of 1 g/L cellulose. (**F**) qRT-PCR validation of gene expression changes observed in panel e.

To further understand the mechanisms of cellulose degradation by *M. comscasis* WC007^T^ and the effect of cellulose degradation on bacterial growth, we performed metabolomic analysis of *M. comscasis* WC007^T^ cultured with or without cellulose. Using the criteria of an OPLS-DA model VIP > 1 and a *P*-value < 0.05, 65 different metabolites were identified when comparing control and cellulose treatment groups (Supplementary file 10). To better understand the candidate metabolites and the relationship between metabolites and differential expression patterns, we conducted a hierarchical cluster analysis based on different metabolites. A heat map of the 65 metabolites shows the production of most detected metabolites (55 vs 65) were up-regulated and a few metabolites (10 vs 65) were down-regulated (Figure 6). The up-regulated metabolites were closely associated with amino acid metabolism, carotenoid biosynthesis, fatty acid biosynthesis, and other cellular processes. The top 20 enriched pathways were further investigated by KEGG analysis (Figure 6-figure supplement 1), which showed aminoacyl-tRNA biosynthesis, biosynthesis of amino acids and 2-Oxocarboxylic acid metabolism pathways were enriched with statistical significance (*P*-value < 0.05). This metabolomic and growth observation data (Figure 3B) as well as transcriptomic analysis (Figure 5) suggest degradation of cellulose promotes the metabolism of different amino acids and other key factors related to energy production.

**Figure 6.**
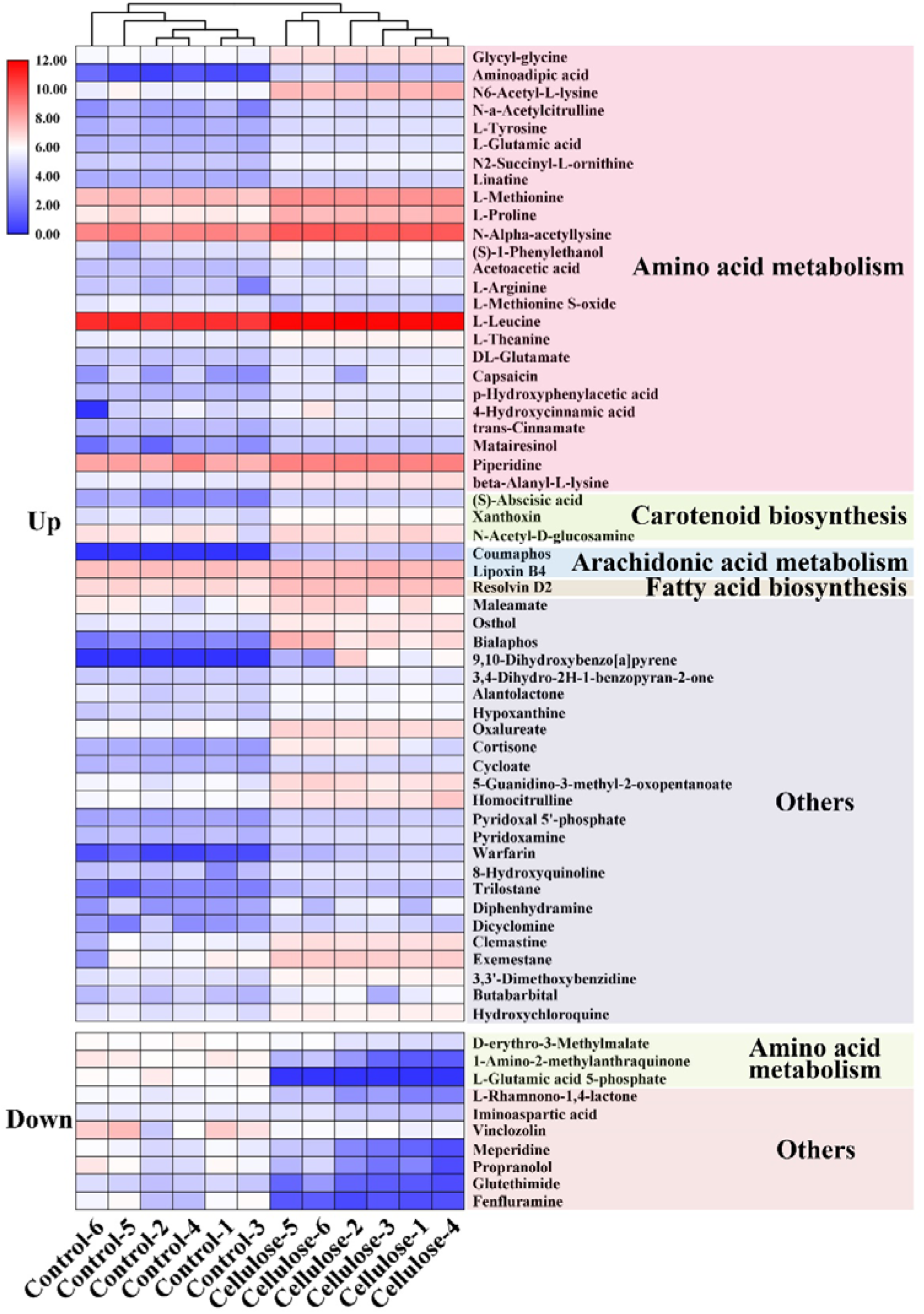
Metabonomic analysis of *M. comscasis* WC007^T^ cultured in inorganic medium supplemented with cellulose. Differential metabolites heatmap showing accumulation of different metabolites in *M. comscasis* WC007^T^ cultured with 1 g/L cellulose. “Up” indicates metabolites are up-regulated in *M. comscasis* WC007^T^ cultured with cellulose. “Down” indicates metabolites that are down-regulated in *M. comscasis* WC007^T^ cultured with cellulose. “Control 1-6” and “Cellulose 1-6” represent six different biological replicates. The color key scale indicates the abundance of metabolites.

### Combined analysis of transcriptomic and metabolomic data during cellulose degradation by *M. comscasis* WC007^T^

To better understand the correlation between our transcriptomic and metabolomic data of *M. comscasis* WC007^T^ cultured with cellulose, we selected common metabolic pathways for further analysis, based on gene expression differences and metabolite production (Supplementary file 11). A total of 18 pathways were enriched, of which 4 pathways were enriched significantly. These 4 pathways were biosynthesis of amino acids, carbon metabolism, arginine biosynthesis and purine metabolism (Figure 7-figure supplement 1). These metabolic pathways are mainly involved in energy production and likely contribute to the growth of *M. comscasis* WC007^T^. Major metabolites involved in saccharide metabolism, amino acids metabolism and tricarboxylic acid (TCA) cycle were increased, and most of the key related enzymes were also up-regulated (Figure 7).

**Figure 7.**
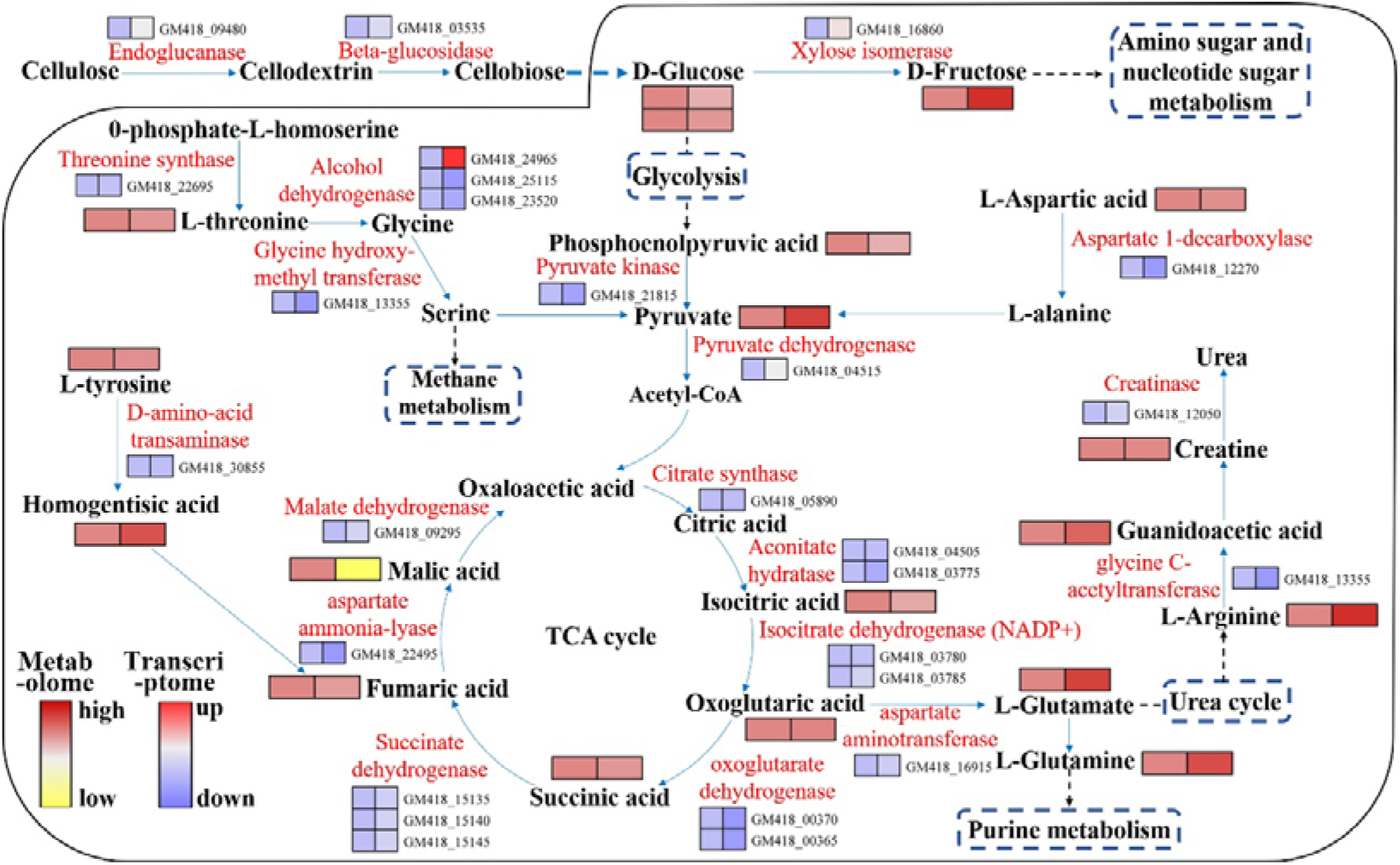
Combined analysis of the *M. comscasis* WC007^T^ transcriptome and metabolome while cultured with cellulose. Metabolite and gene expression changes in three biosynthesis pathways (metabolism of saccharides and amino acids, and TCA cycle) from the combined analysis of transcriptomic and metabolomics data. Enzymes involved in the above pathways are marked in red. The relative content of each metabolite identified in *M. comscasis* WC007^T^ is shown as a heat map from low (yellow) to high (red) as represented by the color scale. Similarly, gene expression levels for each enzyme associated with the above pathways are shown as a heat map from low (blue) to high (red) as represented by the color scale. Two columns for each metabolite or gene represent the control (left) and the cellulose treatment group (right), respectively.

Investigating saccharide metabolism, we found endoglucanase, beta-glucosidase and xylose isomerase gene expression was increased. These enzymes were associated with degradation of cellulose to D-glucose, which is then converted to D-fructose. It is worth noting that crucial amino acids are also increased, including: L-arginine, L-glutamate, L-threonine, homogentisic acid and guanidoacetic acid. These amino acids could produce intermediates required for the TCA cycle (Figure 7). Correspondingly, aconitate hydratase, isocitrate dehydrogenase (NADP^+^), oxoglutarate dehydrogenase and succinate dehydrogenase gene expression was up-regulated. This may be coordinated to enhance the biosynthesis and degradation of isocitric acid, oxoglutaric acid, succinic acid and fumaric acid, and eventually facilitate the TCA cycle and energy production. Overall, the observed changes in metabolite concentrations were consistent with enzyme gene expression changes, further supporting our transcriptomic and metabolomic results and the contribution of cellulose to bacterial growth.

## Discussion

Carbohydrates are a valuable natural resource and account for roughly 75% of the biomass on Earth. Different microorganisms have developed unique strategies to utilize this chemical energy for survival (**Thomas et al., 2011**). Bacteroidetes, globally distributed in human gut, coastal ocean, marine sediments and other environments, play key roles in the degradation of high molecular weight carbohydrates (**Cottrell Kirchman, 2000; Fernandez-Gomez et al., 2013; Thomas et al., 2011**). As such, Bacteroidetes are recognized as one of the dominant phyla of ocean bacterioplankton (**Diez-Vives et al., 2019**) and are the most important decomposer for algae derived carbohydrates, actively driving the ocean carbon and nutrient cycling (**Kruger et al., 2019; Unfried et al., 2018**). As compared to studies of Bacteroidetes derived from the shallow ocean, very little research has been done on the deep-sea counterparts, likely due to the difficulties of sampling and cultivation.

In our current study, we first investigate the abundance of Bacteroides in the deep-sea cold seep through metagenomic analysis. We find that similar to other environments, Bacteroides are the second most abundant bacteria in the deep-sea biotope (Figure 1A). We also find the number of genes encoding CAZymes in Bacteroides is significantly higher than that in other deep-sea bacteria (Figure 1B and 1C), suggesting this phylum plays an essential role in carbohydrate degradation and even the carbon cycle of this habitat. Many Bacteroides strains have been isolated from common environments, however, there are few pure cultures obtained from deep-sea habitats. To enrich for deep-sea Bacteroides, we perform an anaerobic enrichment step, supplementing different polysaccharides in the inorganic medium. As expected, we enrich and isolate a novel species of Bacteroides (*M. comscasis* WC007^T^) capable of degrading various polysaccharides. It is worth noting that this novel species prefers the anaerobic growth conditions of its native deep-sea habitat and duplicates very slowly in aerobic conditions. Most importantly, we develop an efficient goal-directed method to enrich and isolate Bacteroides from deep-sea environments (Figure 2A), a method which will likely be useful for isolating Bacteroides from other environments in the future.

*M. comscasis* WC007^T^ contains numerous glycoside hydrolases (374 GHs) and 82 pairs SusC/D-like protein genes (Figure 2C and 2D). This gene abundance is far more than the number identified in Bacteroides isolated from other environments (Figure 2 – figure supplement 3). We predict *M. comscasis* WC007^T^ has a total of 82 PULs and propose these PULs function to degrade six different polysaccharides (cellulose, pectin, fucoidan, mannan, xylan and starch) (Figure 3A). We validate these findings using growth assays (Figure 3B). The biggest difference between PULs identified in other Bacteroides and *M. comscasis* WC007^T^ is the large number of sigma/anti-sigma factors in the PULs of *M. comscasis* WC007^T^ (Figure 3 – figure supplement 1–6). We find that the expression of these sigma/anti-sigma factors changes with different polysaccharide supplements (Figure 4B), suggesting they function for polysaccharides degradation by *M. comscasis* WC007^T^. Transport and degradation of polysaccharides by *M. comscasis* WC007^T^ depend on typical TonB-dependent regulatory systems (**Koebnik, 2005**). These systems consist of a specialized outer membrane-localized TonB-dependent receptor (such as SusC) that interacts with its energizing TonB-ExbBD protein complex, a cytoplasmic membrane-localized anti-sigma factor and an extracytoplasmic function (ECF)-subfamily sigma factor. This sophisticated complex senses signals from outside the bacterial cell and transmits them via two membranes into the cytoplasm, leading to transcriptional activation of target genes (**Koebnik, 2005**). In Bacteroidetes, predictions have been made for the existence of sigma/anti-sigma factors in the PULs, but few PULs have been verified for their polysaccharide degradation activity (**Koebnik, 2005**). Our transcriptomic results clearly suggest polysaccharide degradation and utilization in *M. comscasis* WC007^T^ is strictly regulated by sigma/anti-sigma factors and is mediated by the TonB-dependent regulatory systems. This may be a universal mechanism for deep-sea Bacteroidetes and warrants further investigation.

From our metagenomic, transcriptomic and metabolomic results, we propose *M. comscasis* WC007^T^ and other Bacteroidetes are major contributors to polysaccharide degradation in the deep-sea environment. Polysaccharide degradation by heterotrophic microbes is known to be a key process in Earth’s carbon and nutrient cycling (**Unfried et al., 2018**). Notably, during cellulose degradation, *M. comscasis* WC007^T^ contributes to the carbon cycle not only by promoting the saccharide and amino acid metabolism but also by facilitating the urea cycle and methane metabolism, the most common carbon source for cold seep microorganisms (Figure 7). Therefore, we propose Bacteroidetes play an essential role in the carbon and nutrient cycling of the deep sea, owing to their polysaccharide degradation functions and biological abundance. Algal polysaccharides are an important bacterial nutrient source and a central component of marine food webs. They are also key contributors to the carbon cycling at the ocean surface and the deep ocean (**Kabisch et al., 2014; Kappelmann et al., 2019; Kruger et al., 2019**). Given the carbon cycling associated with algal polysaccharides in the whole ocean ecosystem, it is reasonable to imagine the most easily degraded polysaccharides are utilized by aerobic microorganisms at the ocean surface (Figure 8). The more difficult-to-degrade polysaccharides aggregate to form particulate detritus and these sinking particles become hotspots for organic carbon, which subsequently play a key role in the export of matter from the euphotic surface to the deep ocean sediment (**Azam Malfatti, 2007**). Once the carbohydrate-derived particles reach the deep sea, efficient degraders like *M. comscasis* WC007^T^ initially recognize the extracellular polysaccharides or oligosaccharides via TonB-dependent outer membrane transporters (SusC and SusD complex). SusC then interacts with TonB through the TonB box, providing energy through the TonB-ExbB-ExbD complex located on the cytoplasmic membrane, allowing these degraders to uptake the oligosaccharides into the periplasm. Meanwhile, TonB-dependent transducers possess a unique N-terminal extension that interacts with the anti-sigma factor and promotes ECF sigma factor binding to the RNA polymerase core enzyme. CAZymes are then activated and released to cleave oligosaccharides into monosaccharide and disaccharides, which are then transported into the cytoplasm for metabolism and energy production to promote bacterial growth. Notably, TonB-dependent uptake systems are sometimes semi-selfish (**Cuskin et al., 2015**), with some cleavage products becoming available to other bacteria, which could greatly promote the carbon and nutrient cycling in the deep biosphere.

**Figure 8.**
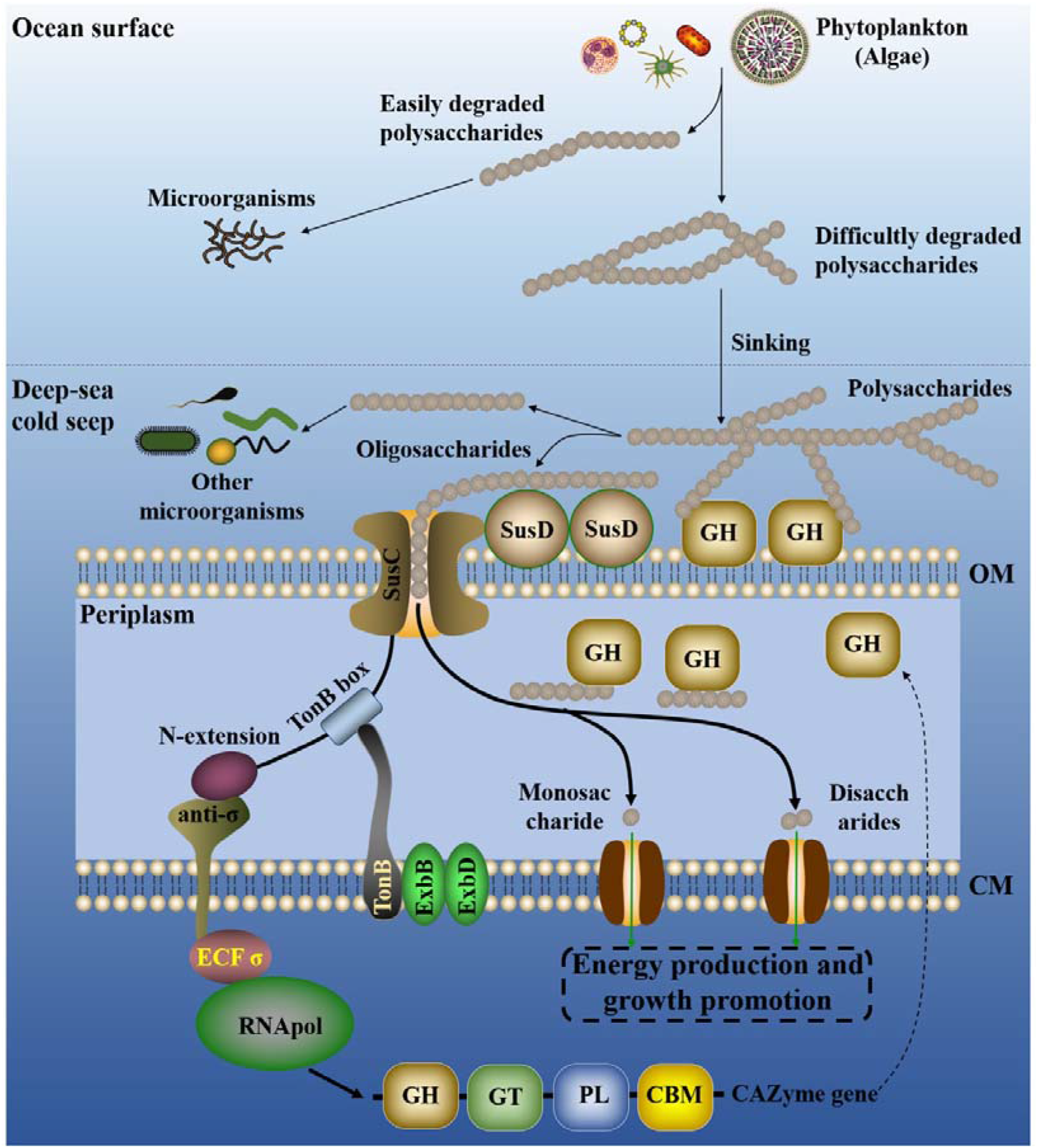
Proposed model of polysaccharide degradation by Bacteroides and contributions to the ocean carbon and nutrient cycling. SusC, starch utilization system subunit C (TonB-dependent receptor/transporter); SusD, starch utilization system subunit D (outer membrane sugar-binding protein); CAZymes, carbohydrate-active enzymes; GH, glycoside hydrolases; GT, glycosyl transferases; PL, polysaccharide lyases; CBM, carbohydrate-binding modules; anti-σ, anti-sigma factor; ECF σ, ECF-subfamily sigma factor; RNApol, RNA polymerase; TonB, a kind of typical periplasmic protein; ExbB and ExbD, plasma membrane proteins providing energy to TonB; CM, cytoplasmic membrane; OM, outer membrane.

## Materials and Methods

### Sampling and operational taxonomic units (OTUs) analysis

The deep-sea sediment sample was collected by *RV KEXUE* from a typical cold seep in the South China Sea (E 119°17’07.322’’, N 22°06’58.598’’) at a depth of approximately 1,146 m in July of 2018 as described previously (**Zhang et al., 2020**). In order to understand the abundance of Bacteroidetes in deep-sea cold seep, operational taxonomic units (OTUs) analysis was performed. Briefly, total genomic DNA extracted from the sample was diluted to 1 ng/μL using sterile water and used for PCR template. Bacterial 16S rRNA genes of distinct regions (16S V4/V5) were amplified using specific primers (515F-GTGCCAGCMGCCGCGG and 907R-CCGTCAATTCMTTTRAGTTT) with the barcode, and the PCR products were purified with Qiagen Gel Extraction Kit (Qiagen, Germany) for library construction. Sequencing library was generated using TruSeq® DNA PCR-Free Sample Preparation Kit (Illumina, USA) following manufacturer’s instructions. And sequences analyses were performed by Uparse software (**Edgar, 2013**). Sequences with ≥97% similarity were assigned to the same OTU. Representative sequence for each OTU was screened for further annotation. For each representative sequence, the Silva Database (**Quast et al., 2013**) was used based on Mothur algorithm to annotate taxonomic information.

### Metagenomic sequencing, assembly, binning and annotation

For metagenomic sequencing, total DNA from the sample (20 g) was extracted using the Qiagen DNeasy® PowerSoil® Pro Kit (Qiagen, Hilden, Germany) and the integrity of DNA was evaluated by gel electrophoresis. Then 0.5 μg DNA of each sample was used to prepare libraries. Libraries were prepared with an amplification step for each sample. DNA was cleaved into 50 ~ 800 bp fragments using Covaris E220 ultrasonicator (Covaris, Brighton, UK) and some fragments between 150 ~ 250 bp were selected using AMPure XP beads (Agencourt, Beverly, MA, USA) and repaired using T4 DNA polymerase (Enzymatics, Beverly, MA, USA). All NGS sequencing was performed on the BGISEQ-500 platform (BGI, Qingdao, China), generating 100 bp paired-end raw reads. Quality control was performed by SOAPnuke (v1.5.6) (setting:-l 20 -q 0.2 -n 0.05 -Q 2 -d -c 0 -5 0 -7 1) (**Chen et al., 2017**) and the clean data were assembled using MEGAHIT (v1.1.3) (setting:--min-count 2 --k-min 33 --k-max 83 --k-step 10) (**Li et al., 2015**). Maxbin2 (**Wu et al., 2016**), metaBAT2 (**Kang et al., 2019**) and Concoct (**Alneberg et al., 2014**) were used to automatically bin from assemblies of these samples. MetaWRAP (**Uritskiy et al., 2018**) was used to purify and organize data to generate the final bins. Finally, completeness and contamination of metagenome-assembled genomes (MAGs) were assessed using checkM (v1.0.18) (**Parks et al., 2015**). These obtained MAGs were subsequently annotated by searching the predicted genes against NR (20180814), KEGG (Release 87.0), COG (update-2018_08) and Swissprot (release-2017_07). Additionally, in order to search for CAZymes genes, CAZymes were annotated based on HMMer, DIAMOND and Hotpep searches of the dbCAN2 meta server (release-2019_07), a web server for automated carbohydrate-active enzyme annotation. CAZymes were annotated and selected only as such when at least two of the searches were positive.

### Isolation and cultivation of deep-sea bacteria

To enrich the bacteria degrading polysaccharide, the sediment sample was cultured at 28 °C for one month in an anaerobic enrichment medium containing (per litre of seawater): 1.0 g NH_4_Cl, 1.0 g NaHCO_3_, 1.0 g CH_3_COONa, 0.5 g KH_2_PO_4_, 0.2 g MgSO_4_.7H_2_O, 1.0 g polysaccharide (cellulose, pectin or xylan), 0.7 g cysteine hydrochloride, 500 μL 0.1 % (w/v) resazurin and pH 7.0. This medium was prepared anaerobically as previously described (**Fardeau et al., 1997**). A 100 μL enrichment was spread on Hungate tube with modified medium (1 g yeast extract, 1 g peptone, 1 g NH_4_Cl, 1 g NaHCO_3_, 1 g CH_3_COONa, 0.5 g KH_2_PO_4_, 0.2 g MgSO.7H_2_O, 0.7 g cysteine hydrochloride, 500 μL 0.1 % (w/v) resazurin, 1,000 mL seawater, 15 g agar, pH 7.0) after 10,000 times dilution, and this medium was named ORG in this study. The Hungate tubes were anaerobically incubated at 30 °C for 7 days. Individual colonies with distinct morphology were picked using sterilized bamboo sticks and then cultured in the ORG broth. *Maribellus comscasis* WC007^T^ was isolated and purified with ORG medium by repeated use of the Hungate roll-tube methods for several rounds until it was considered to be axenic. The purity of strain WC007^T^ was confirmed routinely by transmission electron microscopy (TEM) and repeated partial sequencing of the 16S rRNA gene. Strain WC007^T^ is preserved at −80 °C in ORG broth supplemented with 20% (v/v) glycerol.

### Electron microscopy observation

To observe the morphological characteristics of *M. comscasis* WC007^T^, cells were examined using transmission electronic microscopy (TEM) (HT7700; Hitachi, Japan) with a JEOL JEM 12000 EX (equipped with a field emission gun) at 100 kV. The cells suspension of *M. comscasis* WC007^T^ was washed with Milli-Q water and centrifuged at 5,000 *g* for 5 min. Subsequently, the sample was taken by immersing copper grids coated with a carbon film for 20 min in the bacterial suspensions and washed for 10 min in distilled water and dried for 20 min at room temperature(**Han et al., 2014**).

To detect the degradation of cellulose by *M. comscasis* WC007^T^, 10 μL sample was dripped on cover slips and soaked in gelatin and dried for 30 min to allow the sample to adhere to the surface of copper grid. These samples were fixed in 2.5% glutaraldehyde for 30 min. Samples were then washed three times with PBS and dehydrated in ethanol solutions of 30%, 50%, 70%, 90% and 100% for 10 min each time. All samples were observed with scanning electronic microscopy (SEM) (S-3400N, Hitachi, Japan) at 5 kV.

### Phylogenetic analysis

Phylogenetic trees were constructed with the full-length 16S rRNA sequences by the neighbor-joining algorithm (**Saitou Nei, 1987**), maximum Likelihood (**Felsenstein, 1981**) and minimum-evolution methods (**Rzhetsky Nei, 1992**). 16S rRNA sequences of *M. comscasis* WC007^T^ (accession number MT460676) and other related taxa used for phylogenetic analysis were obtained from NCBI GenBank. Phylogenetic analysis was performed using the software MEGA version 6.0 (**Tamura et al., 2013**). And a genome-based phylogenetic tree was reconstructed using the TYGS algorithm (https://tygs.dsmz.de/) (**Meier-KolthoffGoker, 2019**).

### Genomic characterizations of *M. comscasis* WC007^T^

Genome relatedness values were calculated by multiple approaches: Average Nucleotide Identity (ANI) based on the MUMMER ultra-rapid aligning tool (ANIm), ANI based on the BLASTN algorithm (ANIb), tetranucleotide signatures (TETRA), and *in silico* DNA-DNA similarity. ANIm, ANIb, and TETRA frequencies were calculated using JSpecies WS (http://jspecies.ribohost.com/jspeciesws/) (**Richter et al., 2016**). Recommended species criteria cut-offs were used: 95% for ANIb and ANIm and 0.99 for TETRA signature (**Richter Rossello-Mora, 2009**http://ekhidna2.biocenter.helsinki.fi/AAI/old). Amino acid identity (AAI) values were calculated by AAI-profiler (http://ekhidna2.biocenter.helsinki.fi/AAI/) (**Medlar et al., 2018**). *In silico* DNA-DNA similarity values were calculated by the Genome-to-Genome Distance Calculator (GGDC) (http://ggdc.dsmz.de/) (**Meier-Kolthoff et al., 2013**). The *is*DDH results were based on the recommended formula 2, which is independent of genome size and is robust when using whole-genome sequences.

### PULs prediction and annotation

PULs of *M. comscasis* WC007^T^ were detected based on the presence of *susC*-like and *susD*-like gene pairs, which in most cases was also accompanied by the presence of CAZyme clusters, as previously suggested (**Bjursell et al., 2006**). Additional consideration required a PUL to have at least one *susC*- and *susD*-like gene pair and encode at least one degradative CAZymes from the GH families. PUL genes were predicted using a combination of HMMer searches against the Pfam (**Finn et al., 2014**), TIGRFAM (**Selengut et al., 2007**), dbCAN (**Yin et al., 2012**), MEROPS (**Rawlings et al., 2012**) and CAZy (**Lombard et al., 2014**) databases. CAZymes were annotated based on HMMer searches against the Pfam v25 and dbCAN 3.0 databases and BLASTp searches (**Altschul et al., 1990**) against the CAZy database. SusC-like protein (TonB-linked outer membrane protein) and SusD-like protein (starch-binding associating with outer membrane) were annotated by the DOE-JGI Microbial Annotation Pipeline (MGAP) (**Huntemann et al., 2016**), which used the TIGRfam model (TIGR04056) to detect SusC-like proteins and the Pfam models (PF12741, PF12771, and PF14322) to detect SusD-like proteins (**Terrapon et al., 2015**). Sulfatases were annotated by the SulfAtlas database v1.0 (**Barbeyron et al., 2016**). Peptidases were annotated based on the BLASTp searches against the MEROPS 9.13 database using the default settings of E≤10^−4^.

### Growth assay of *M.comscasis* WC007^T^

Growth assays were performed at atmospheric pressure. Briefly, 100 μL *M.comscasis* WC007^T^ culture was inoculated in Hungate tubes containing 10 mL inorganic medium (1.0 g/L NH_4_Cl, 1.0 g/L NaHCO_3_, 1.0 g/L CH_3_COONa, 0.5 g/L KH_2_PO_4_, 0.2 g/L MgSO_4_.7H_2_O, 0.7 g/L cysteine hydrochloride, 500 μL/L of 0.1 % (w/v) resazurin and pH 7.0) with different polysaccharides (1.0 g/L cellulose, 1.0 g/L pectin, 0.1 g/L fucoidan, 0.1 g/L mannan, 1.0 g/L xylan or 1.0 g/L starch), respectively. The Hungate tubes were anaerobically incubated at 30 °C for 10 days. Since *M.comscasis* WC007^T^ grew very slowly in the inorganic culture medium, we adopted the colony forming units (CFU) to measure the growth curve of *M.comscasis* WC007^T^ in an anaerobic cabinet.

### Transcriptomics analysis

Transcriptomics analysis was performed by Novogene (Tianjin, China). Cells suspension of *M. comscasis* WC007^T^ cultured in inorganic medium for 7 days was used as a control, while cell suspensions cultured in inorganic medium supplemented with different polysaccharides (1.0 g/L cellulose, 1.0 g/L pectin, 0.1 g/L fucoidan, 0.1 g/L mannan, 1.0 g/L xylan or 1.0 g/L starch) for 7 days were the experimental groups. All cultures were incubated in 2 L anaerobic bottles. For transcriptomics analyses, total *M. comscasis* WC007^T^ RNA was extracted using TRIzol reagent (Invitrogen, USA) and DNA contamination was removed using the MEGA clear™ Kit (Life technologies, USA). Detailed protocols of the following procedures including library preparation, clustering and sequencing and data analyses were described in the Supplementary information.

### Metabolomics analysis

Metabolomics analysis was performed by Genedenovo Biotechnology (Guangzhou, China). The samples for metabolomics analysis were obtained from the culture supernatant used for transcriptomic analysis, where *M. comscasis* WC007^T^ was cultured in inorganic medium supplemented with 1.0 g/L cellulose for 7 days. Briefly, 100 mg supernatant was taken and placed in containers, extraction liquid containing an internal target was added and the mixture was homogenized in ball mill kept in ice water for 4 min at 45 Hz with ultrasound treatment. After 3 rounds of homogenization, the samples were incubated for 1 h at -20 °C to precipitate proteins. Samples were then centrifuged at 12,000 *g* for 15 min at 4 °C and the supernatant was transferred into fresh tubes. Extracts were dried in a vacuum concentrator without heating and extraction liquid was added for reconstitution. Finally, the extracts were shaken for 30 s and centrifuged at 12,000 *g* for 15 min at 4 °C. Supernatants were transferred into 2 mL LC/MS glass vials for the LC-MS/MS analysis by using an UHPLC system (Agilent Technologies 1290 with a UPLC BEH Amide column) coupled to TripleTOF 6600 (Q-TOF, AB Sciex).

Metabolite identification was performed based on public metabolite databases, such as HMDB (https://www.hmdb.ca/) (**Wishart et al., 2013**), METLIN (https://metlin.scripps.edu/index.php) (**Zhu et al., 2013**), MassBank (https://www.massbank.jp/), LipidMaps (http://www.lipidmaps.org), mzClound (https://www.mzcloud.org). Principle component analysis (PCA), partial least squares discriminant analysis (PLS-DA) and hierarchical clustering analysis (HCA) were used to integrate the obtained metabolite mass spectral peaks.

The threshold of variables determined to be important in the projection (VIP) scores ≥ 1.0 together with a *P*-value of *T*-test < 0.05 were adopted to assess significant different metabolites. All differentially expressed genes and metabolites were mapped to the Kyoto Encyclopedia of Genes and Genomes (KEGG) KEGG database for pathway and enrichment analysis (**Okuda et al., 2008**).

## Data availability

The BioProject and BioSample accession numbers of metagenome-assembled genomes (MAGs) are shown in the Supplementary file 1. The complete genome sequence of *M. comscasis* WC007^T^ has been deposited at GenBank under the accession number CP046401. The raw sequencing reads for transcriptomic analysis have been deposited to NCBI Short Read Archive (accession number: PRJNA664312).

## Description of *Maribellus comscasis* sp. nov.

*Maribellus comscasis* (com.sca’sis. L. gen. pl. n. *comscasis* from the Center for Ocean Mega-Science, Chinese Academy of Sciences).

Cells are Gram-stain-negative curve-shaped, 2.0-6.0 μm in length and 0.5-0.8 μm in width. Facultative anaerobic and oxidase-positive. The temperature range for growth is 28-37 °C with an optimum at 30 °C. Growing at pH values of 6.0-8.0 (optimum, pH 7.0). Growth occurs at NaCl concentrations between 0.0-5.0% with optimum growth at 1.0% NaCl. By analyzing the hydrolysis of polysaccharides, the growth is promoted significantly by cellulose, pectin and xylan. From the sole carbon source utilization test, growth is stimulated by acetate, maltose, fructose, lactate, sorbitol and D-mannose. Weak growth occurs with ethanol and formate. The major polar lipids are phosphatidylethanolamine, unidentified phospholipid, unidentified aminolipid, unidentified lipid. Containing significant proportions (>10%) of the cellular fatty acids iso-C_15:0_, C_16:0_, summed feature 3 (containing C_16:1_ω7*c* and/or C_16:1_ω6*c*) and summed feature 8 (containing C_18:1_ω7*c* and/or C_18:1_ω6*c*).

The type strain, WC007^T^ (=MCCC 1K04777^T^ = KCTC 25169^T^), was isolated from the sediment of deep-sea cold seep, P.R. China. The DNA G+C content of the type strain is 38.4%.

## Acknowledgements

This work was funded by the Strategic Priority Research Program of the Chinese Academy of Sciences (Grant No. XDA22050301), China Ocean Mineral Resources R&D Association Grant (Grant No. DY135-B2-14), National Key R and D Program of China (Grant No. 2018YFC0310800), the Taishan Young Scholar Program of Shandong Province (tsqn20161051), and Qingdao Innovation Leadership Program (Grant No. 18-1-2-7-zhc) for Chaomin Sun. This study is also funded by the Open Research Project of National Major Science & Technology Infrastructure (*RV KEXUE*) (Grant No. NMSTI-KEXUE2017K01).

## Author contributions

RZ and CS conceived and designed the study; RZ conducted most of the experiments; RL and GL collected the samples from the deep-sea cold seep; RC helped to analyze the metagenomes; RZ and CS lead the writing of the manuscript; all authors contributed to and reviewed the manuscript.

## Figures Supplements

**Figure 1 – figure supplement 1.**
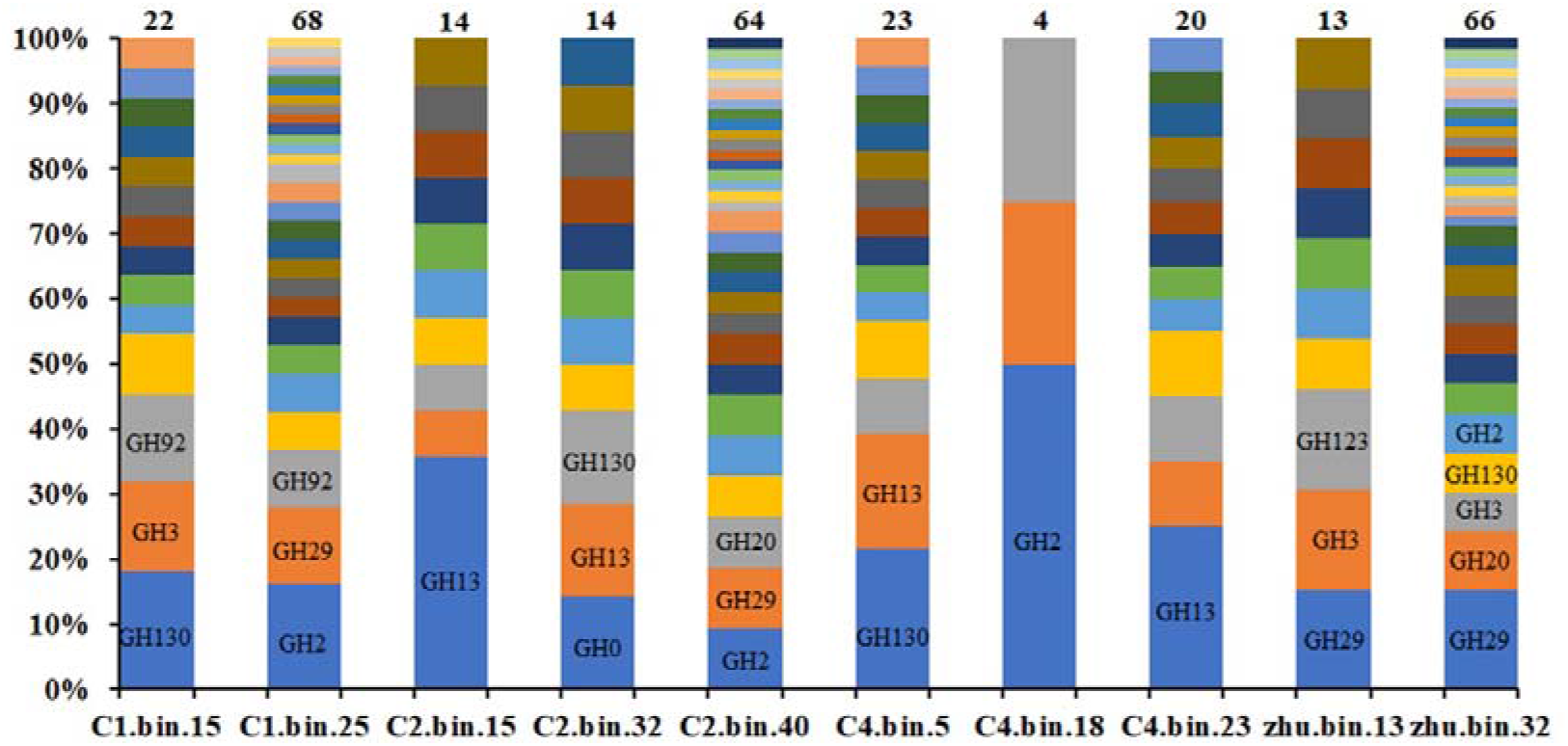
Calculation of the numbers of GH families distributed in the metagenome-assembled genomes (MAGs) of Bacteroides derived from the deep-sea cold seep. The number above the column indicates the number of GH families identified in corresponding Bacteroides MAG. The X-axis indicates the names of different Bacteroides MAGs. The Y-axis indicates the ratio of each GH family number to the total number of GH families identified in each Bacteroides MAG.

**Figure 2 – figure supplement 1.**
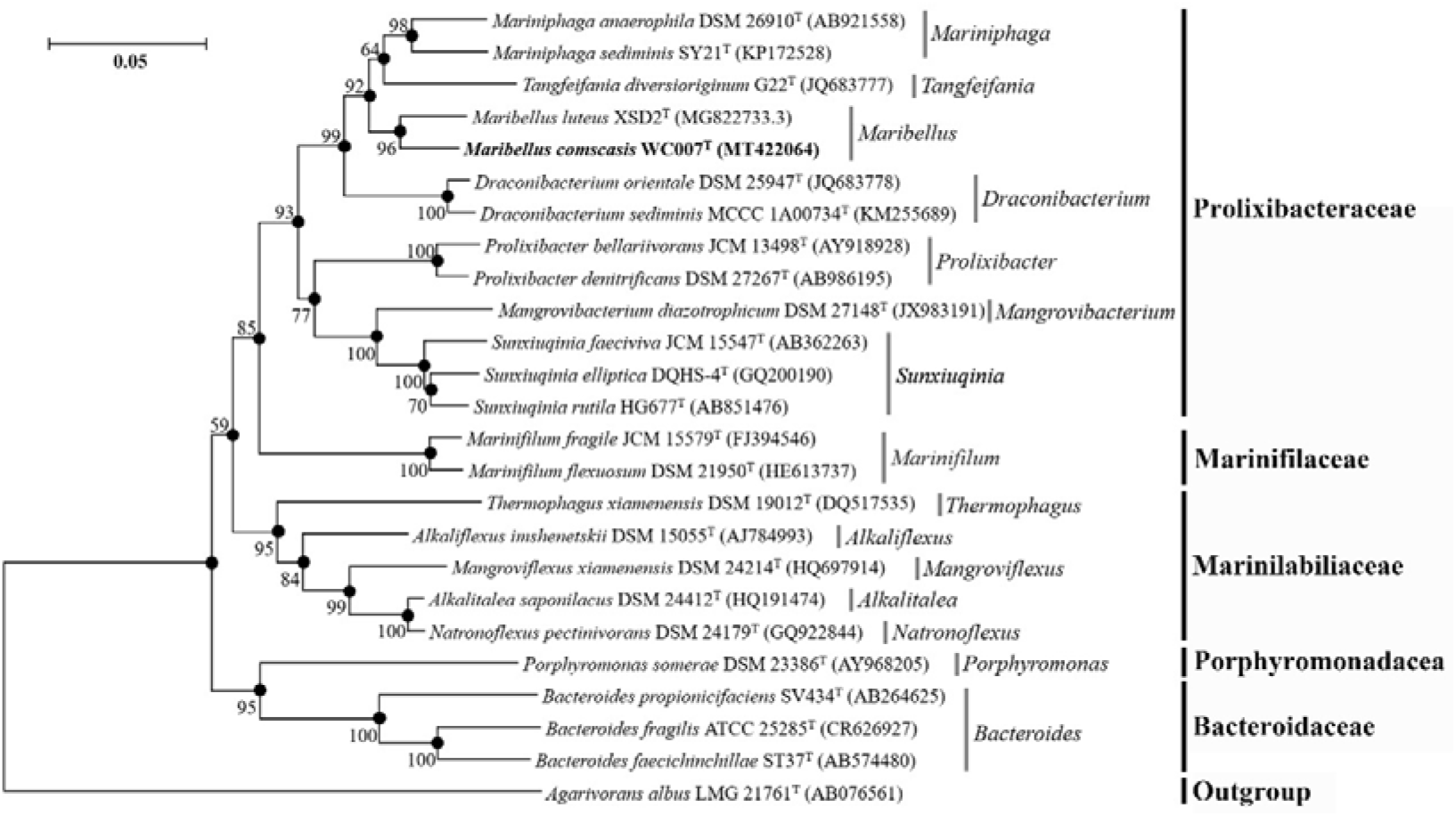
Neighbor-joining tree based on 16S rRNA sequences showing the position of the novel species *Maribellus comscasis* WC007^T^. The filled circles indicate branches of the tree that were also formed using the maximum-likelihood and minimum-evolution methods. The numbers above or below the branches are bootstrap values based on 1,000 replicates. The access number of each 16S rRNA is indicated after the strain’s name. The sequence of *Agarivorans albus* LMG 21761^T^ is used as an outgroup. Bar, 0.05 substitutions per nucleotide position.

**Figure 2 – figure supplement 2.**
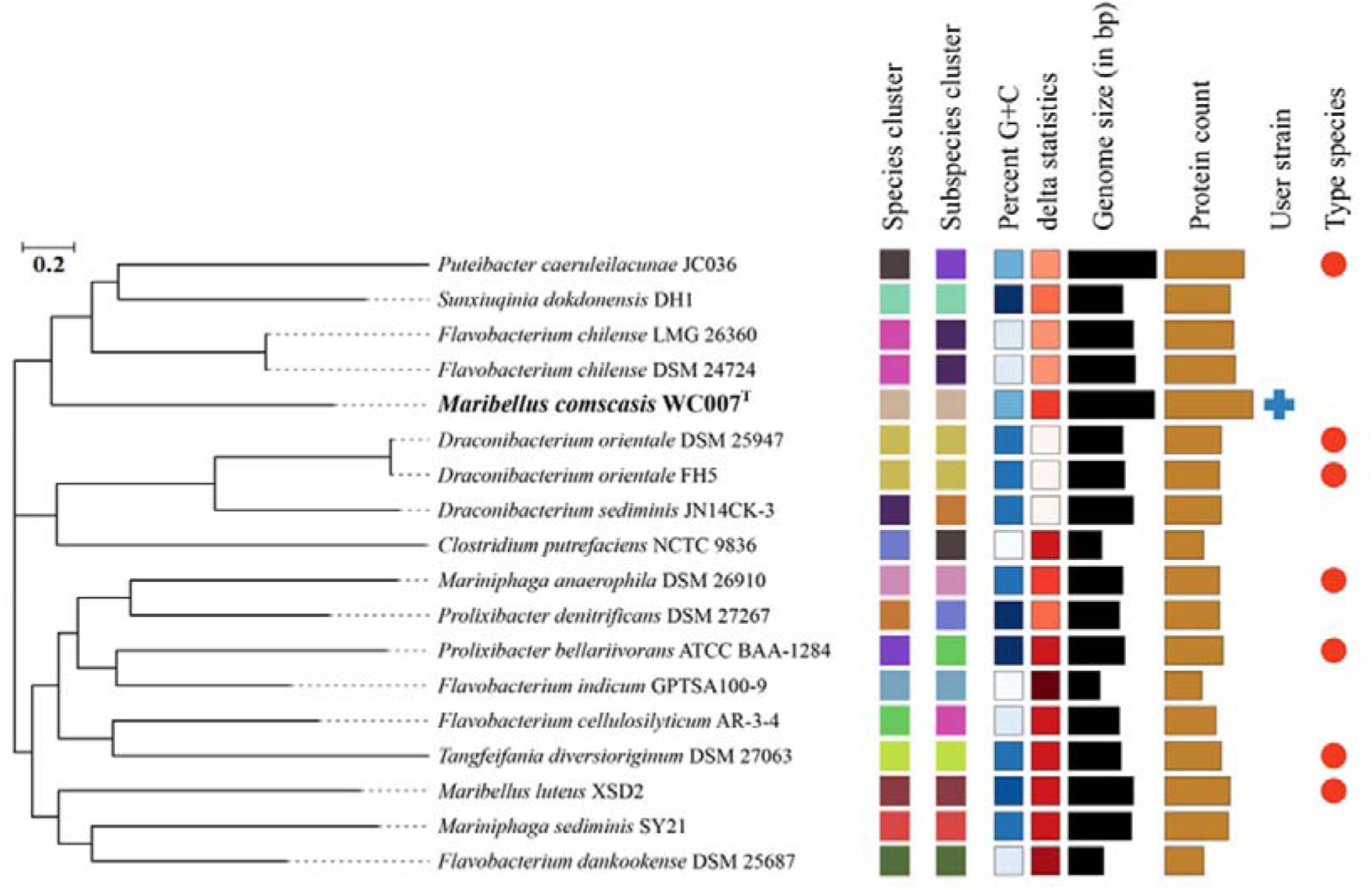
Phylogenomic tree analysis of *M. comscasis* WC007^T^ based on the TYGS algorithm (https://tygs.dsmz.de/).

**Figure 2 – figure supplement 3.**
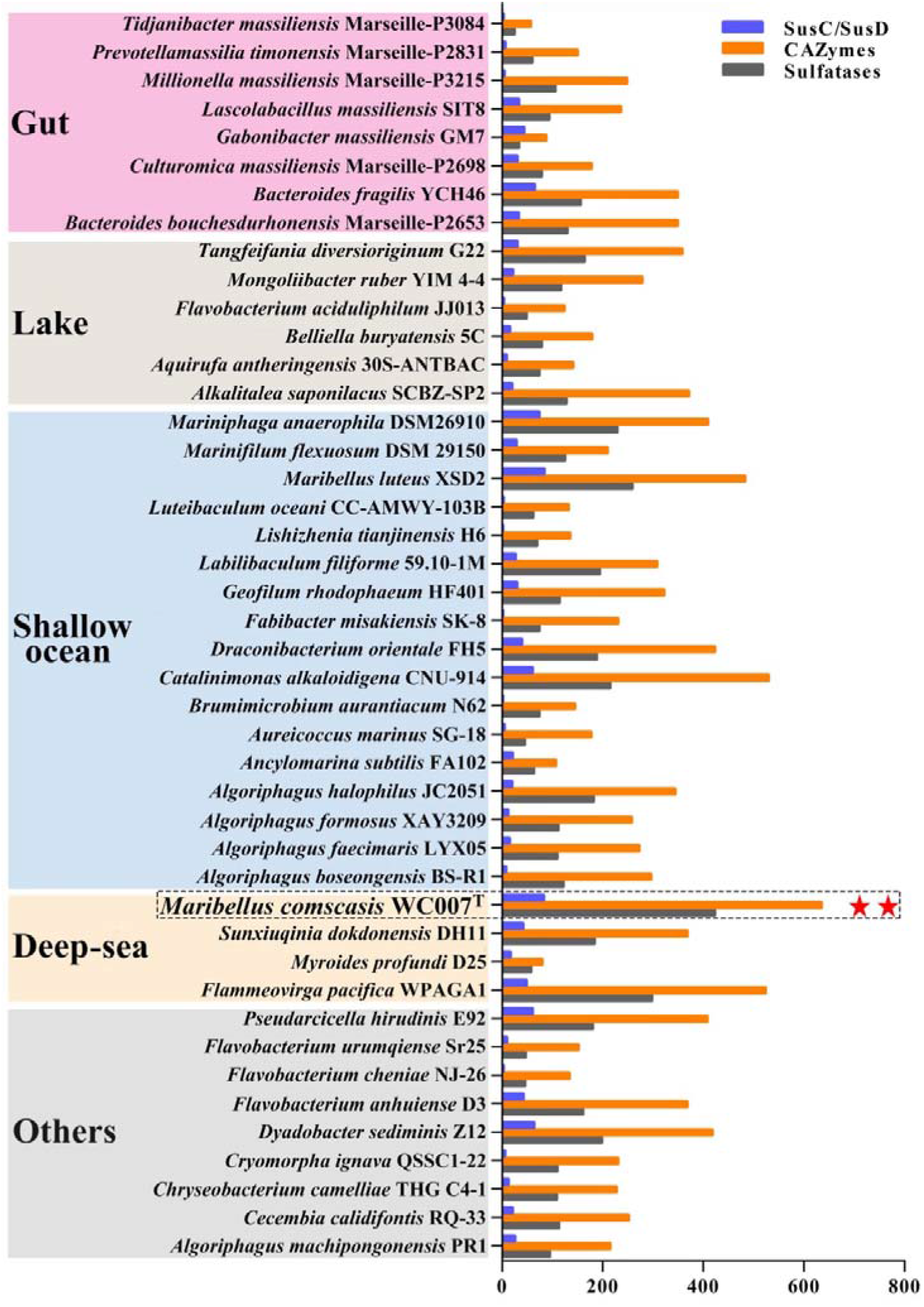
Comparison of the number of genes encoding SusC/SusD pairs, CAZymes and sulfatases identified in the genomes of Bacteroides from different environments (including gut, lake, shallow ocean, deep-sea etc.). *M. comscasis* WC007^T^ is framed by a dotted line and indicated with two pentacle stars. The detailed information related to this figure is listed in the Supplementary file 7.

**Figure 3 – figure supplement 1.**
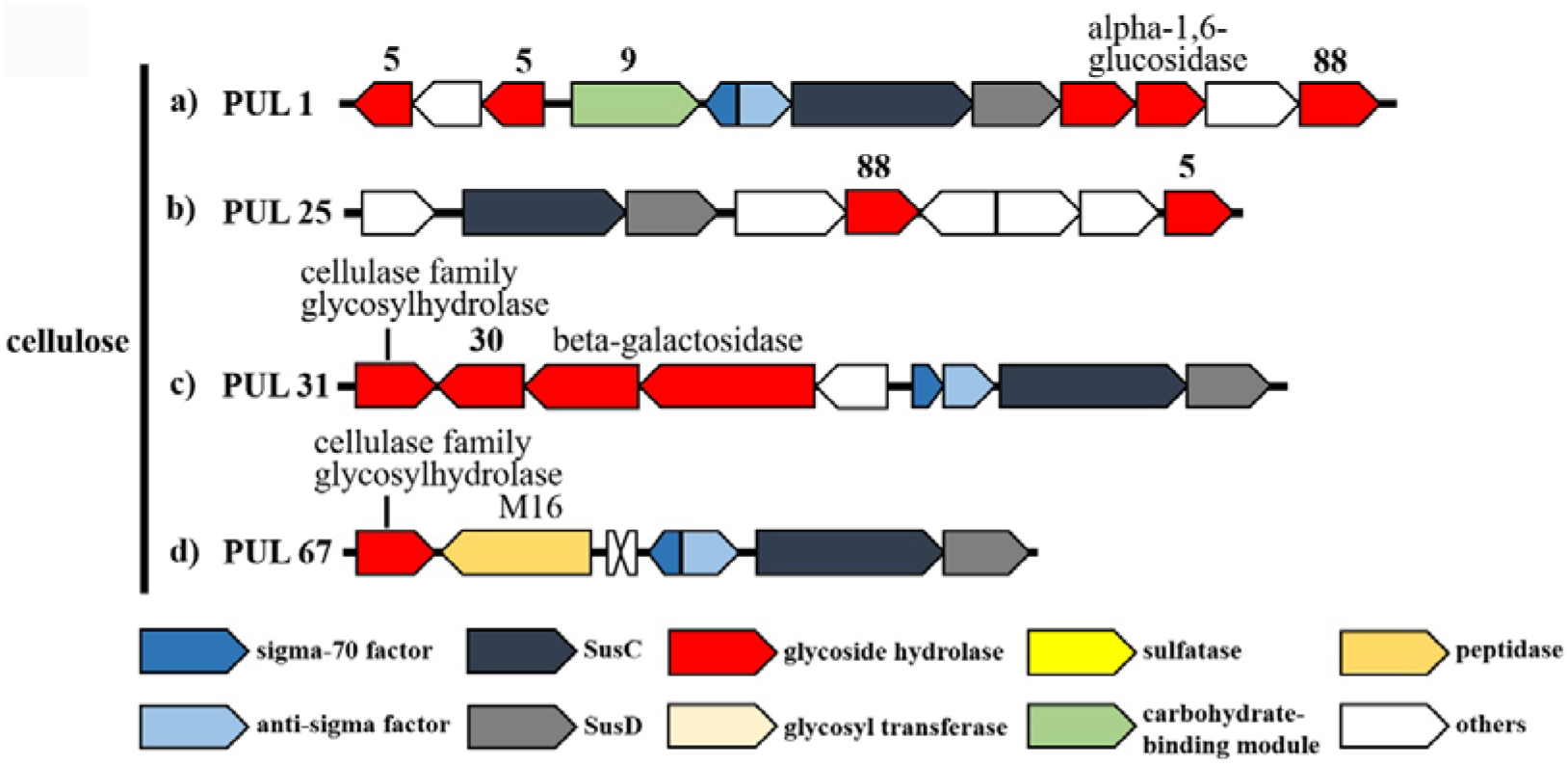
PULs predicted to target cellulose in the genome of *M. comscasis* WC007^T^. Color coding of genes indicates different gene types and corresponding numbers indicate CAZymes family associations.

**Figure 3 – figure supplement 2.**
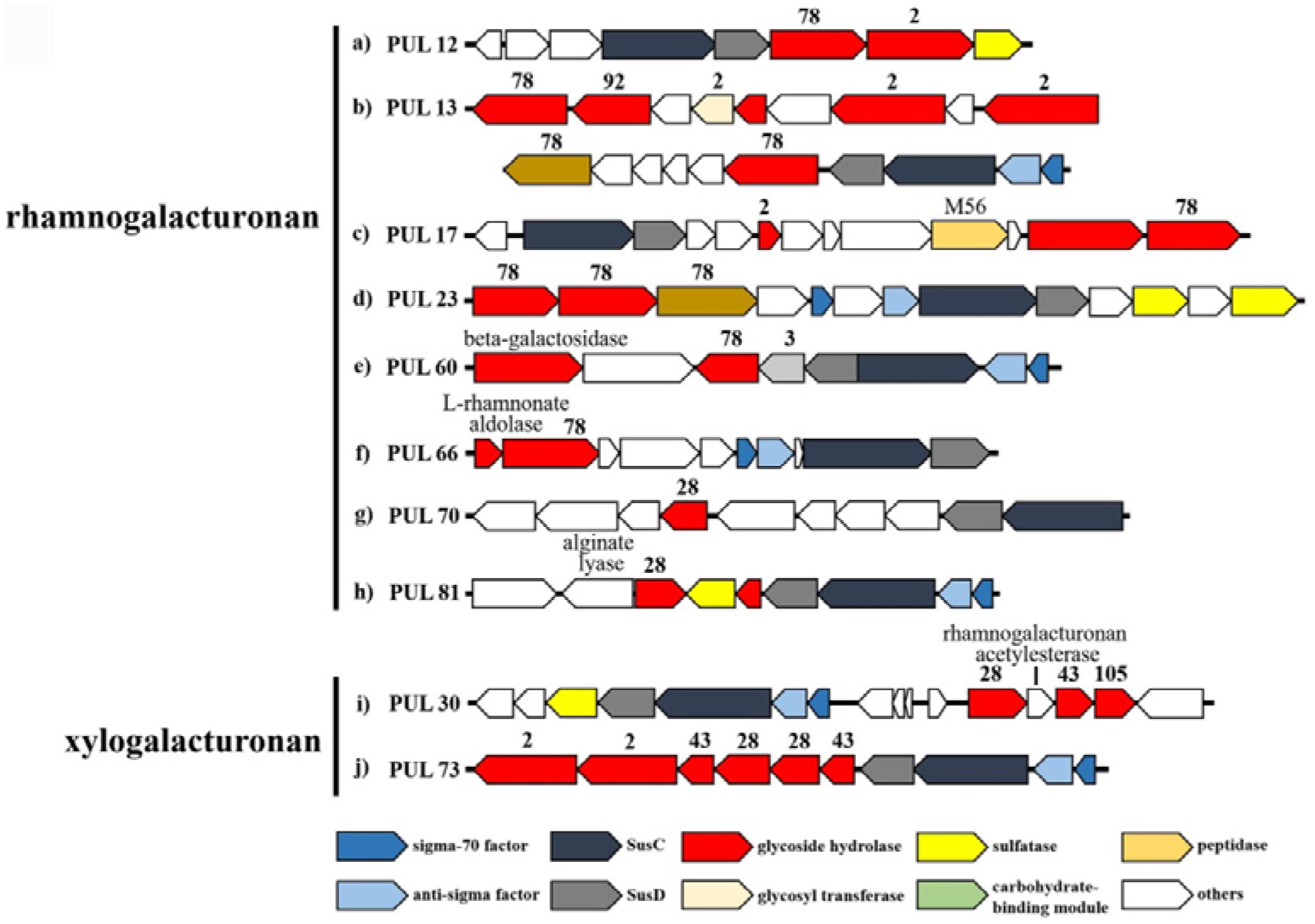
PULs predicted to target pectin (mainly rhamnogalacturonan and xylogalacturonan) in the genome of *M. comscasis* WC007^T^. Color coding of genes indicates gene types and corresponding numbers indicate CAZymes family associations.

**Figure 3 – figure supplement 3.**
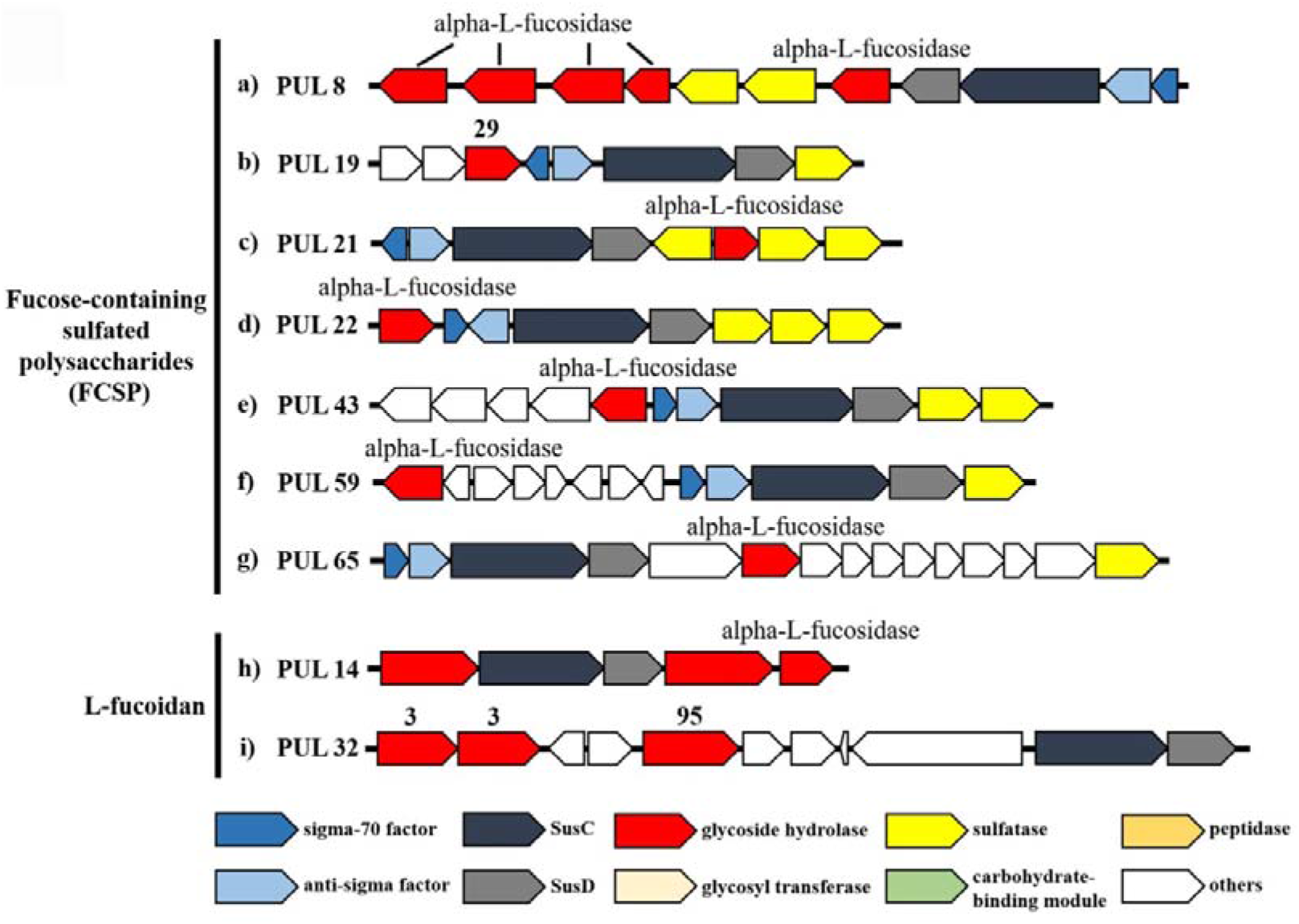
PULs predicted to target fucoidan (mainly fucose-containing sulfated polysaccharides and L-fucoidan) in the genome of *M. comscasis* WC007^T^. Color coding of genes indicates gene types and corresponding numbers indicate CAZymes family associations.

**Figure 3 – figure supplement 4.**
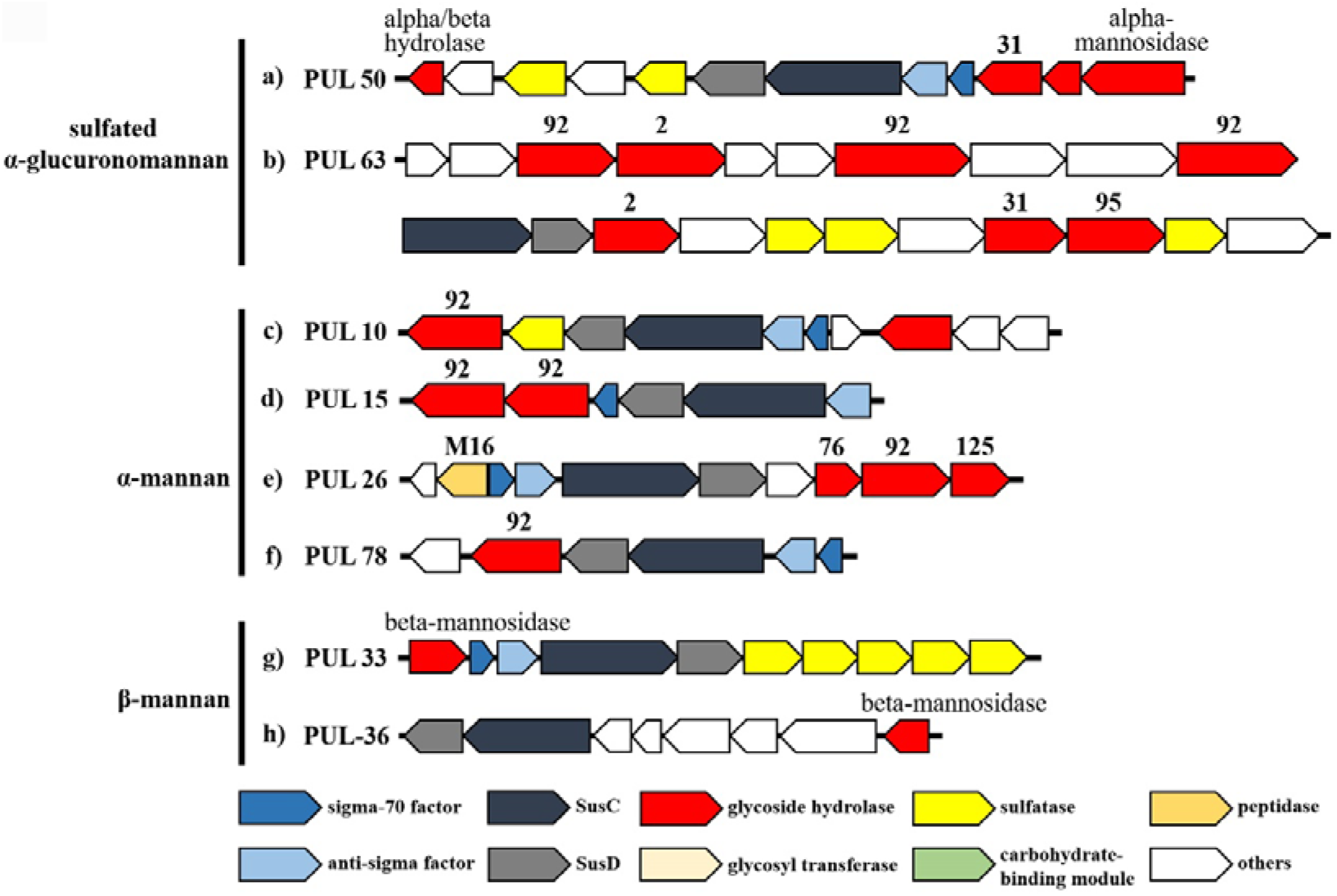
PULs predicted to target mannan (sulfated α-glucuronomannan, α-mannan and β-mannan) in the genome of *M. comscasis* WC007^T^. Color coding of genes indicates gene types and corresponding numbers indicate CAZymes family associations.

**Figure 3 – figure supplement 5.**
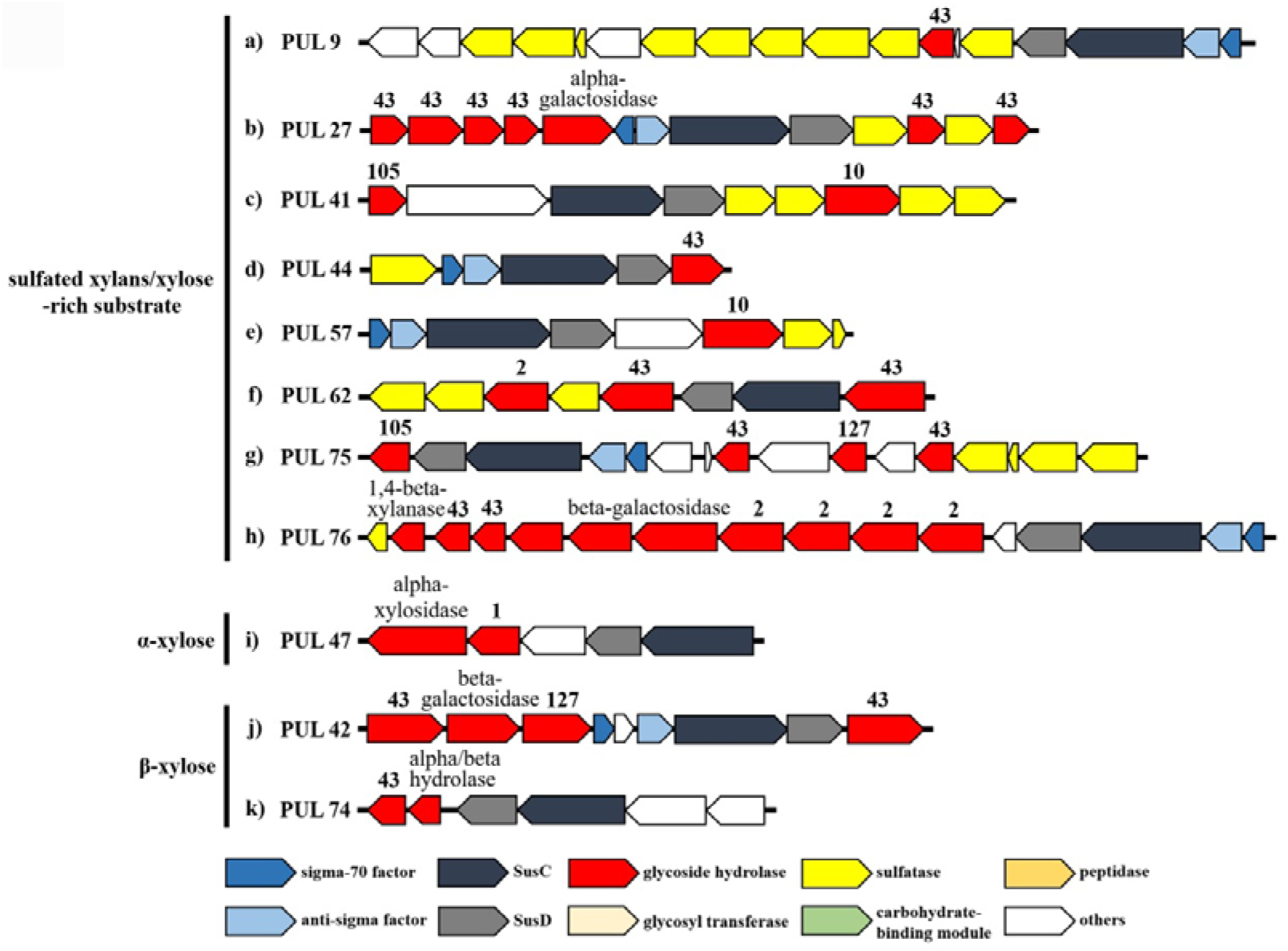
PULs predicted to target xylan (mainly sulfated xylan, α-xylose and β-xylose), α-mannan and β-mannan in the genome of *M. comscasis* WC007^T^. Color coding of genes indicate gene types and corresponding numbers indicate CAZymes family associations.

**Figure 3 – figure supplement 6.**
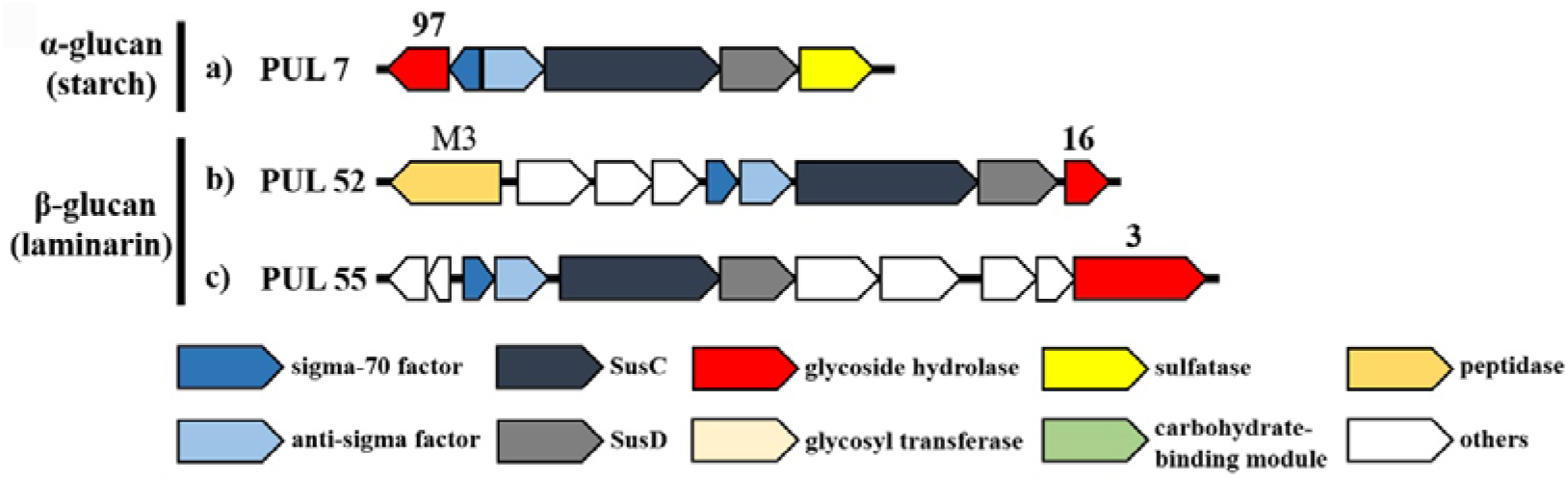
PULs predicted to target glucan (mainly starch and laminarin) in the genome of *M. comscasis* WC007^T^. Color coding of genes indicates gene types and corresponding numbers indicate CAZymes family associations.

**Figure 4 – figure supplement 1.**
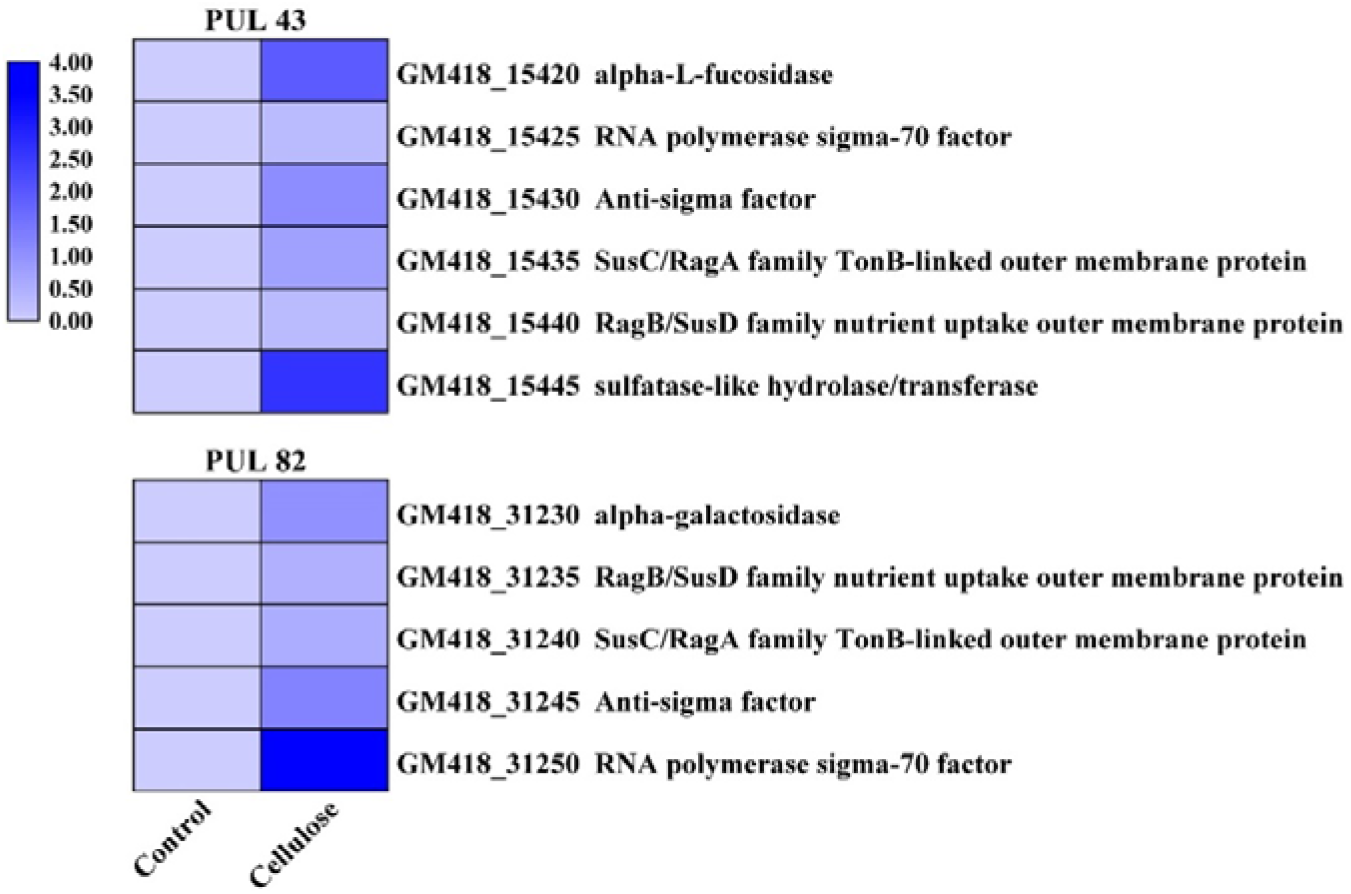
Transcriptomics based heat map showing the PULs with all genes up-regulated when cultured *M. comscasis* WC007^T^ in the medium supplemented with 1 g/L cellulose.

**Figure 4 – figure supplement 2.**
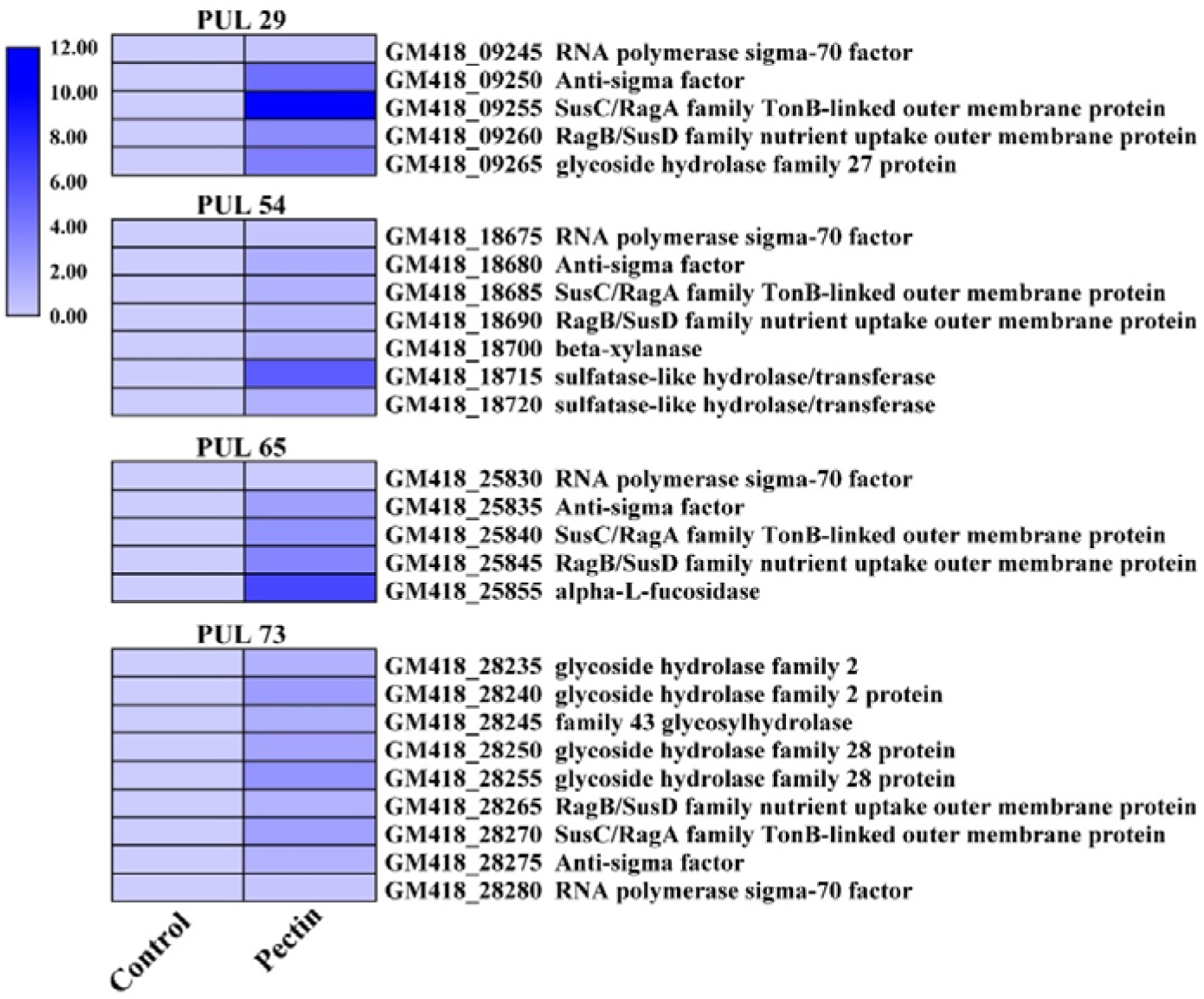
Transcriptomics based heat map showing the PULs with all genes up-regulated when cultured *M. comscasis* WC007^T^ in the medium supplemented with 1 g/L pectin.

**Figure 4 – figure supplement 3.**
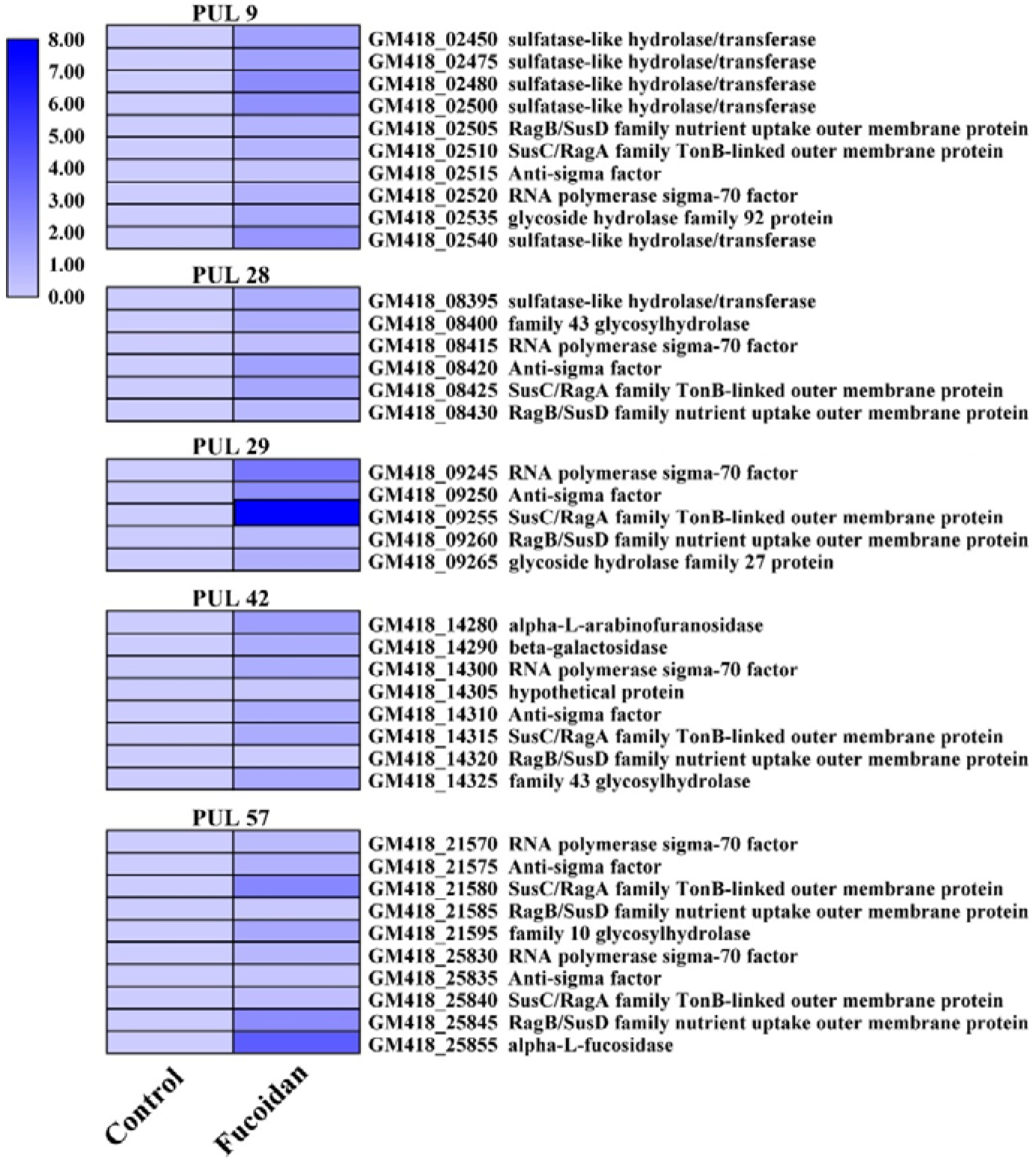
Transcriptomics based heat map showing the PULs with all genes up-regulated when cultured *M. comscasis* WC007^T^ in the medium supplemented with 1 g/L fucoidan.

**Figure 4 – figure supplement 4.**
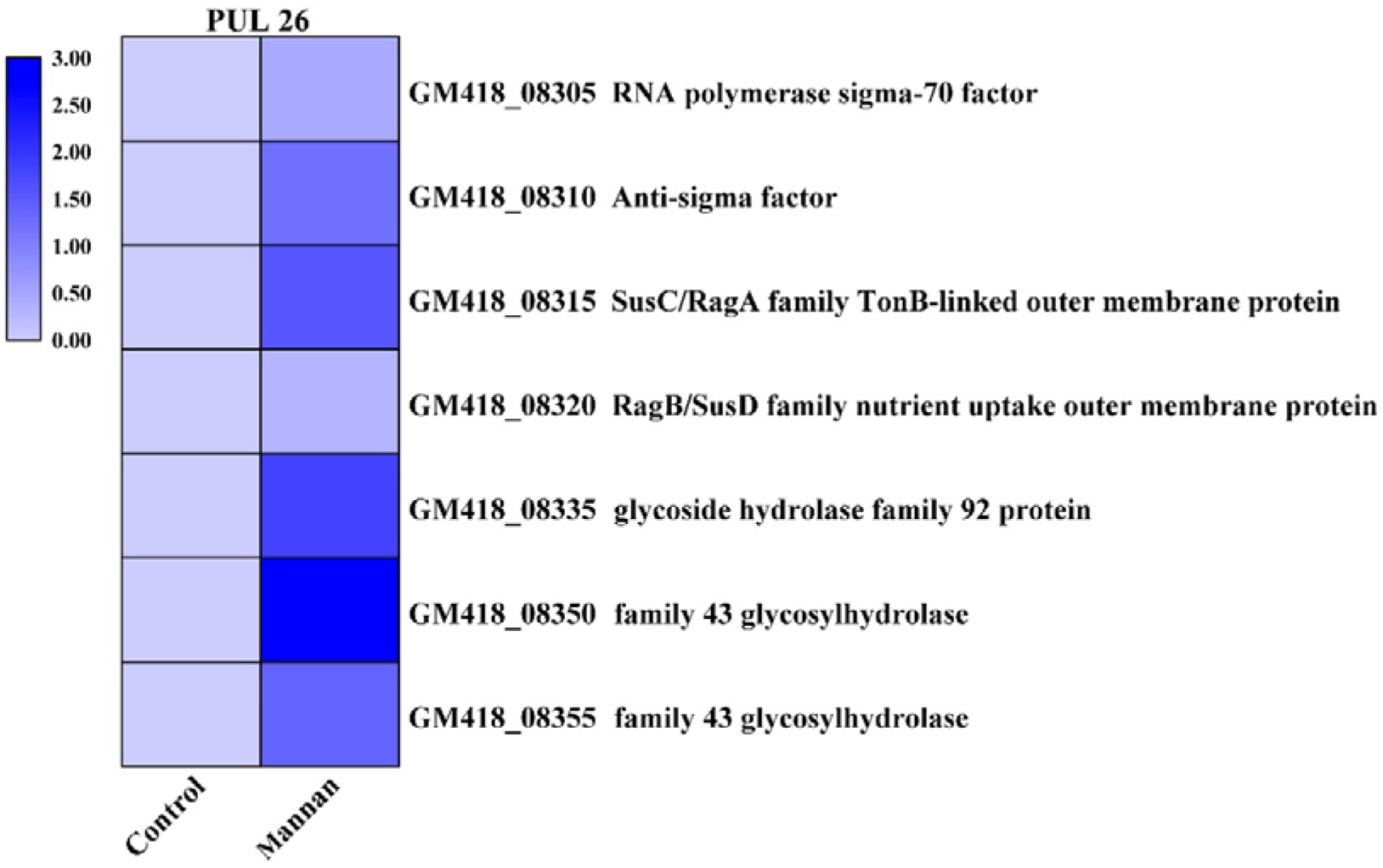
Transcriptomics based heat map showing the PULs with all genes up-regulated when cultured *M. comscasis* WC007^T^ in the medium supplemented with 1 g/L mannan.

**Figure 4 – figure supplement 5.**
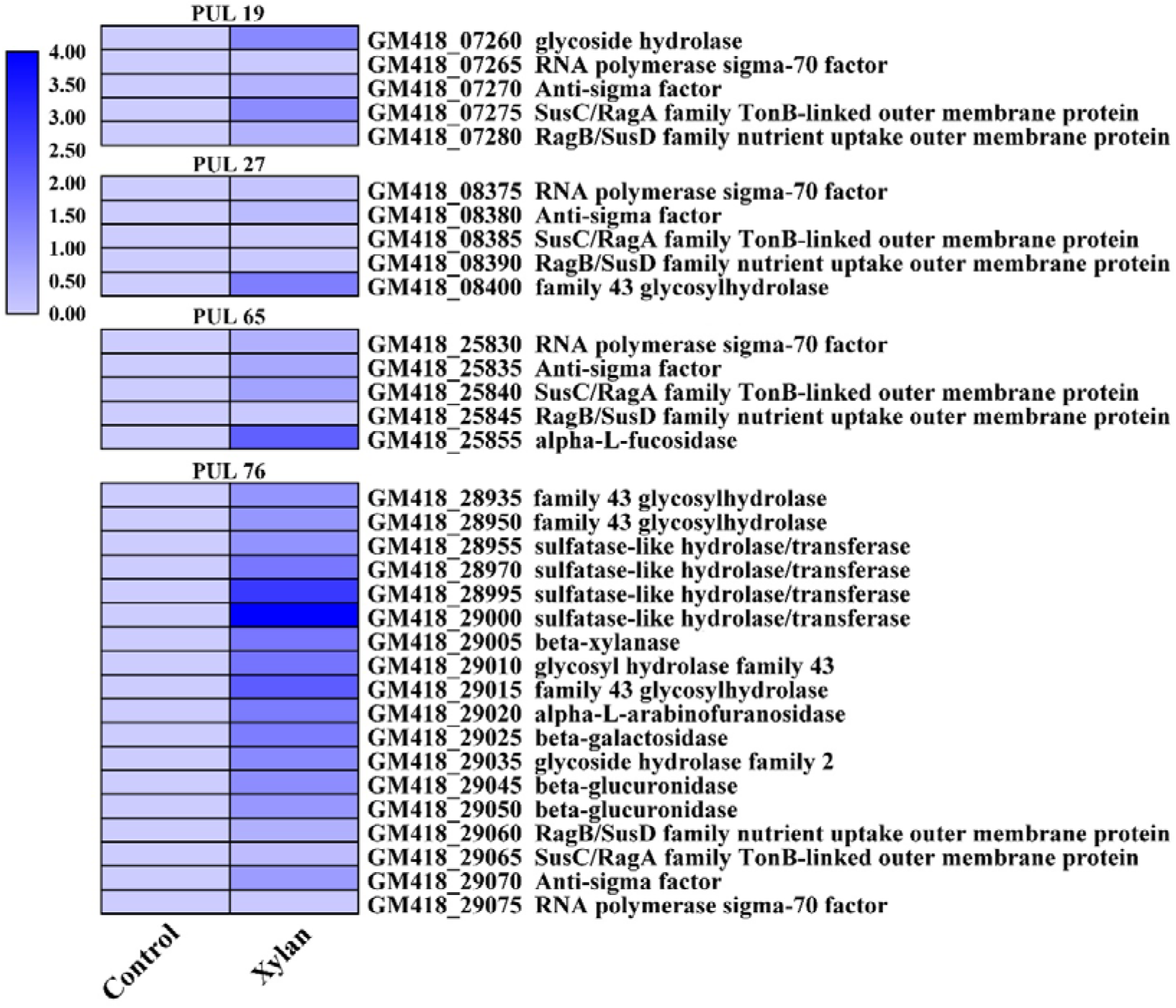
Transcriptomics based heat map showing the PULs with all genes up-regulated when cultured *M. comscasis* WC007^T^ in the medium supplemented with 1 g/L xylan.

**Figure 4 – figure supplement 6.**
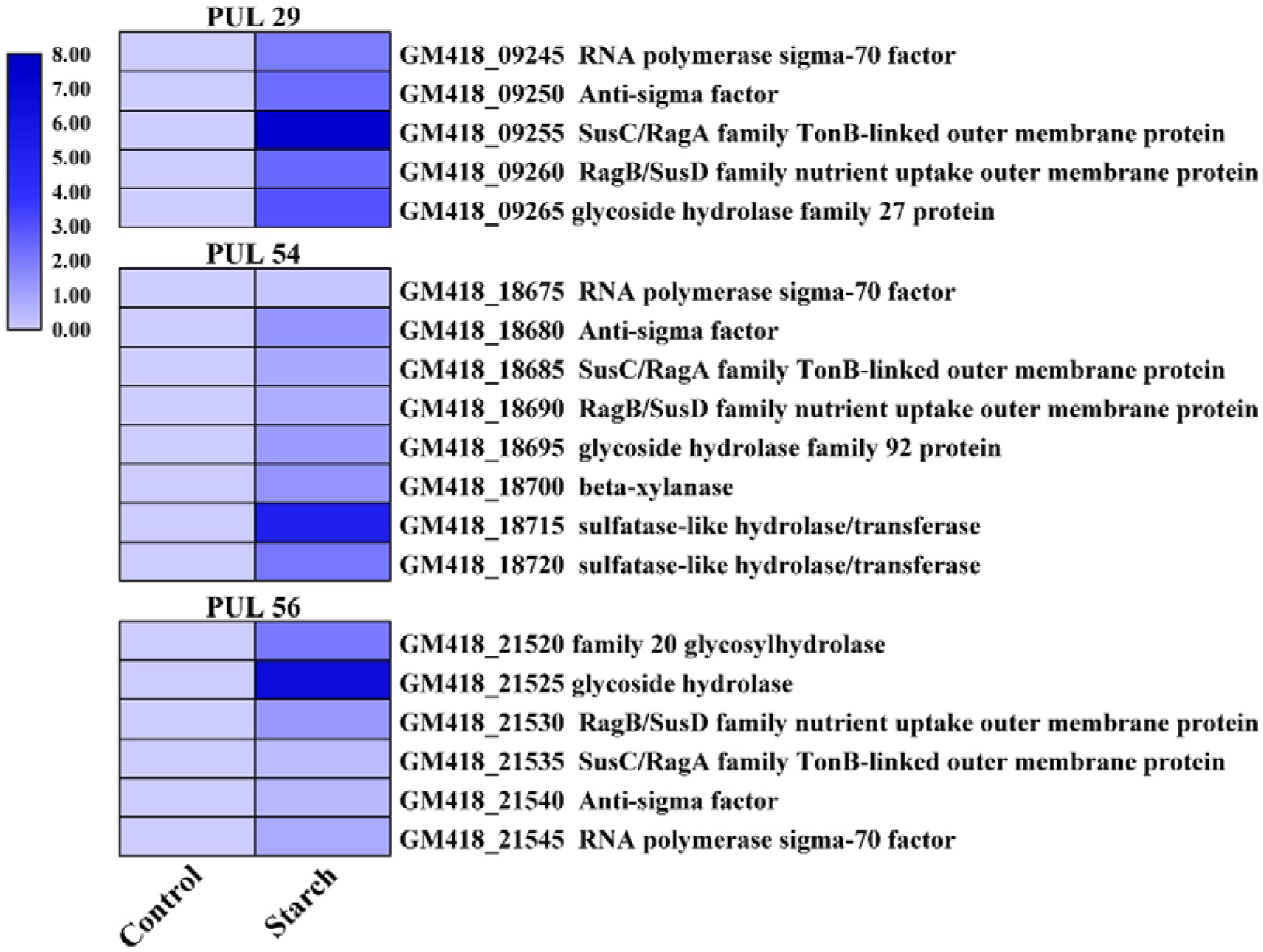
Transcriptomics based heat map showing the PULs with all genes up-regulated when cultured *M. comscasis* WC007^T^ in the medium supplemented with 1 g/L starch.

**Figure 6 – figure supplement 1.**
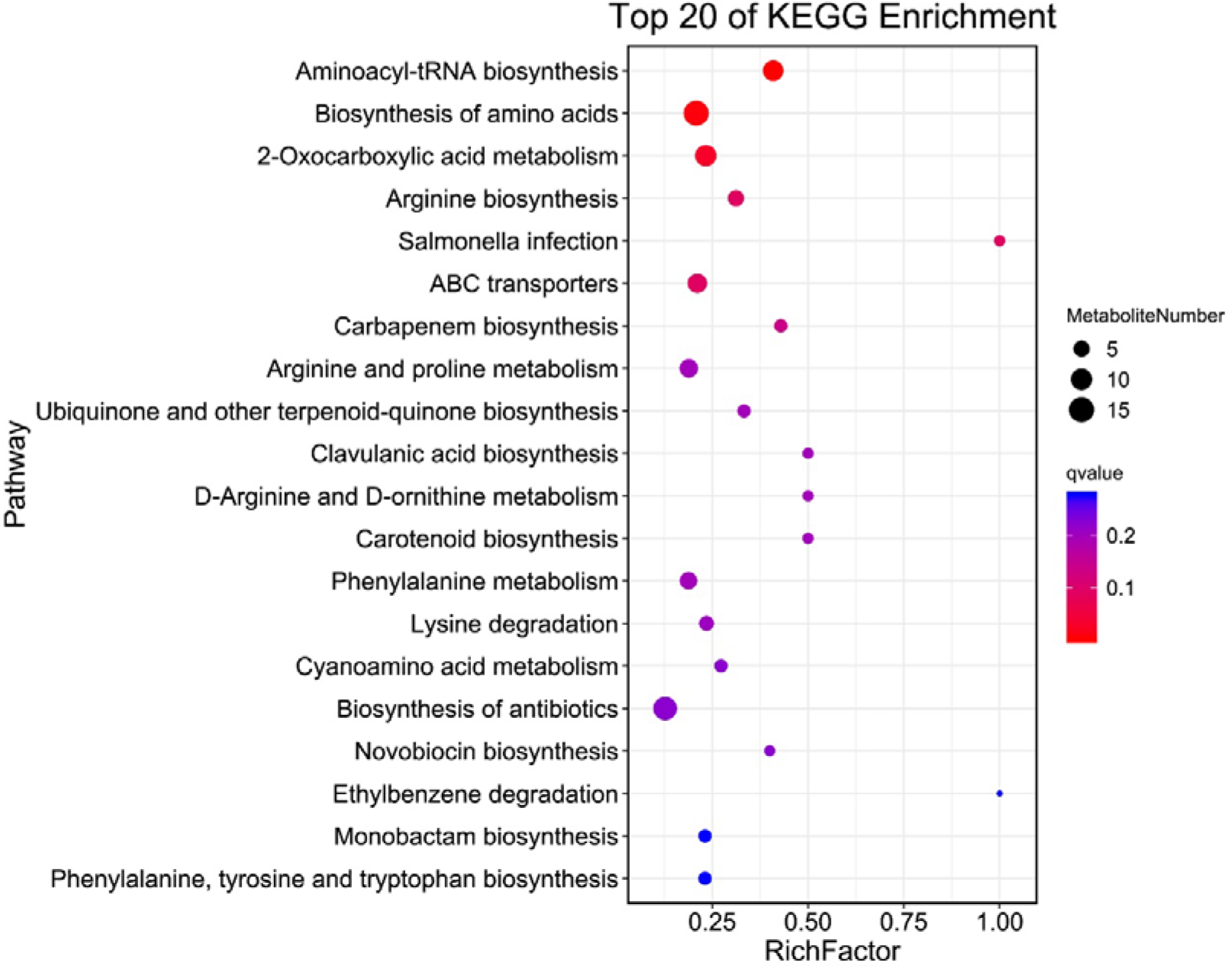
KEGG pathway enrichment of differential metabolites when cultured *M. comscasis* WC007^T^ in the medium supplemented with 1 g/L cellulose. Only Top 20 pathway enrichments were shown.

**Figure 7 – figure supplement 1.**
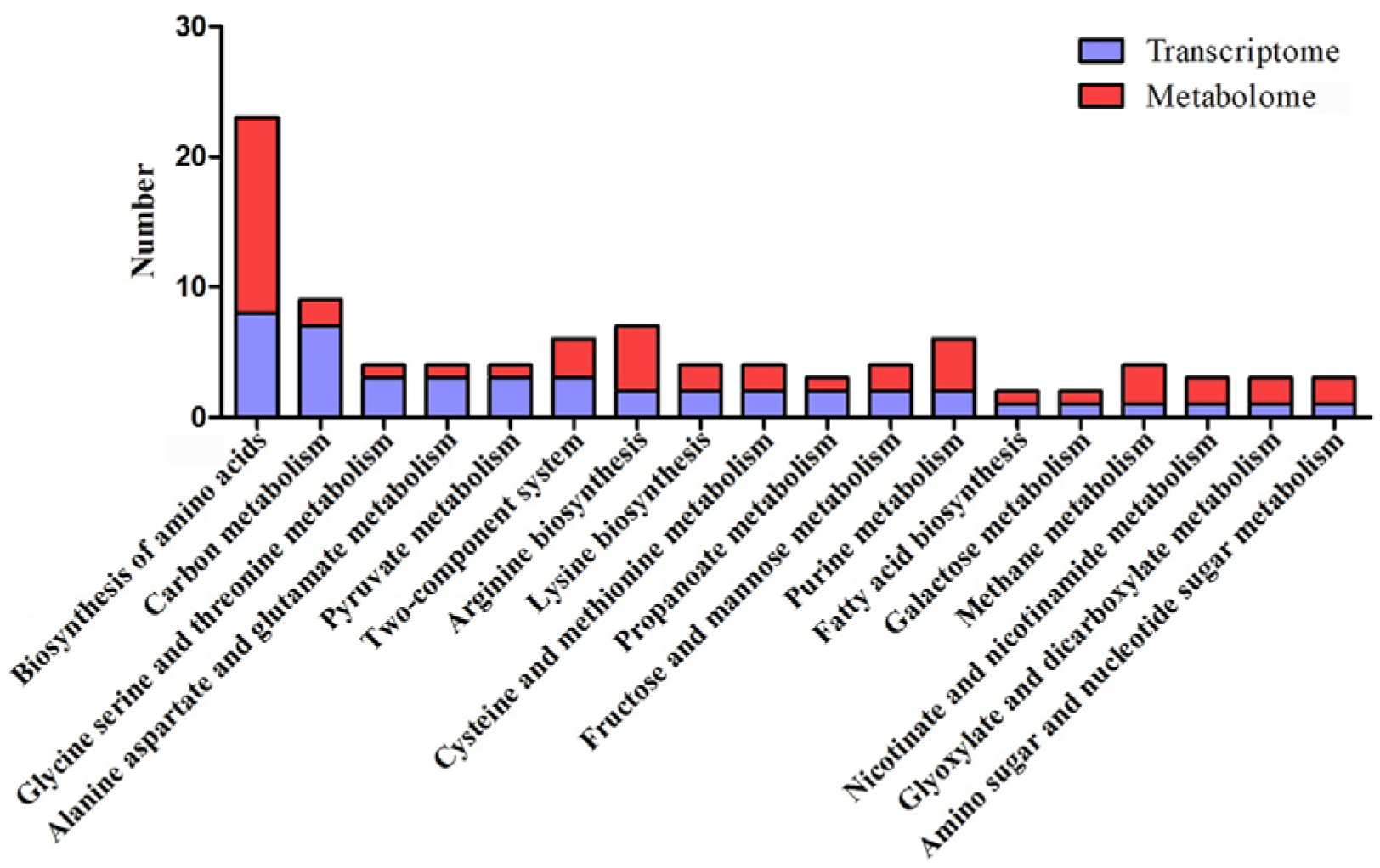
Combination analysis of enriched pathways associated genes and metabolites respectively identified in the transcriptomic and metabonomic analyses of *M. comscasis* WC007^T^ treated with 1 g/L cellulose. Blue column represents the number of differentially expressed genes identified in transcriptomic analysis. Red column represents the number of differentially abundant metabolites identified in metabonomic analysis.

