## Supplemental Files 1-8 for "Bacteroidetes contribute to the carbon and nutrient cycling of deep sea through breaking down diverse glycans"

**Materials and Methods**

**Transcriptional profiling of *M.comscasis* WC007^T^ cultured with different polysaccharides**

**(1) Library preparation for strand-specific transcriptome sequencing**

A total amount of 3 μg RNA per sample was used as input material for the RNA sample preparations. Sequencing libraries were generated using NEBNext^®^ Ultra™ Directional RNA Library Prep Kit for Illumina^®^ (NEB, USA) following manufacturer’s recommendations and index codes were added to attribute sequences to each sample. rRNA is removed using a specialized kit that leaves the mRNA. Fragmentation was carried out using divalent cations under elevated temperature in NEBNext First Strand Synthesis Reaction Buffer (5×). First strand cDNA was synthesized using random hexamer primer and M-MuLV Reverse Transcriptase (RNaseH^-^). Second strand cDNA synthesis was subsequently performed using DNA Polymerase I and RNase H. In the reaction buffer, dNTPs with dTTP were replaced by dUTP. Remaining overhangs were converted into blunt ends via exonuclease/polymerase activities. After adenylation of 3’ ends of DNA fragments, NEBNext Adaptor with hairpin loop structure was ligated to prepare for hybridization. In order to select cDNA fragments of preferentially 150~200 bp in length, the library fragments were purified with AMPure XP system (Beckman Coulter, Beverly, USA). Then 3 μL USER Enzyme (NEB，USA) was used with size-selected, adaptor-ligated cDNA at 37 °C for 15 min followed by 5 min at 95 °C before PCR. Then PCR was performed with Phusion High-Fidelity DNA polymerase, Universal PCR primers and Index (X) Primer. At last, products were purified (AMPure XP system) and library quality was assessed on the Agilent Bioanalyzer 2100 system.

**(2) Clustering and sequencing**

The clustering of the index-coded samples was performed on a cBot Cluster Generation System using TruSeq PE Cluster Kit v3-cBot-HS (Illumia) according to the manufacturer’s instructions. After cluster generation, the library preparations were sequenced on an Illumina Hiseq platform and paired-end reads were generated.

**(3) Data analysis**

Raw data (raw reads) of fastq format were firstly processed through in-house perl scripts. In this step, clean data (clean reads) were obtained by removing reads containing adapter, reads containing ploy-N and low quality reads from raw data. At the same time, Q20, Q30 and GC content the clean data were calculated. All the downstream analyses were based on the clean data with high quality. Reference genome and gene model annotation files were downloaded from genome website directly. Both building index of reference genome and aligning clean reads to reference genome were used Bowtie2-2.2.3 (**LangmeadSalzberg, 2012**). HTSeq v0.6.1 was used to count the reads numbers mapped to each gene. And then FPKM of each gene was calculated based on the length of the gene and reads count mapped to this gene. FPKM, expected number of Fragments Per Kilobase of transcript sequence per Millions base pairs sequenced, considers the effect of sequencing depth and gene length for the reads count at the same time, and is currently the most commonly used method for estimating gene expression levels (**Trapnell et al., 2009**).

**(4) Differential expression analysis**

Differential expression analysis of two conditions/groups (two biological replicates per condition) was performed using the DESeq R package (1.18.0) (**AndersHuber, 2010**). DESeq provide statistical routines for determining differential expression in digital gene expression data using a model based on the negative binomial distribution. The resulting *P*-values were adjusted using the Benjamini and Hochberg’s approach for controlling the false discovery rate. Genes with an adjusted *P*-value < 0.05 found by DESeq were assigned as differentially expressed. (For DEGSeq without biological replicates) Prior to differential gene expression analysis, for each sequenced library, the read counts were adjusted by edgeR program package through one scaling normalized factor. Differential expression analysis of two conditions was performed using the DEGSeq R package (1.20.0) (**Wang et al., 2010**). The *P* values were adjusted using the Benjamini & Hochberg method. Corrected *P*-value of 0.005 and log_2_ (Fold change) of 1 were set as the threshold for significantly differential expression.

**(5) GO and KEGG enrichment analysis of differentially expressed genes**

Gene Ontology (GO) enrichment analysis of differentially expressed genes was implemented by the GOseq R package, in which gene length bias was corrected (**Young et al., 2010**). GO terms with corrected *P* value less than 0.05 were considered significantly enriched by differential expressed genes. KEGG is a database resource for understanding high-level functions and utilities of the biological system, such as the cell, the organism and the ecosystem, from molecular-level information, especially large-scale molecular datasets generated by genome sequencing and other high-throughput experimental technologies (<http://www.genome.jp/kegg/>) (**Kanehisa et al., 2008**). We used KOBAS software to test the statistical enrichment of differential expression genes in KEGG pathways.

**Results**

**Prediction of PULs in the genome of *M. comscasis* WC007^T^**

**(1) PULs of cellulose.** Four PULs predicted to target cellulose are found in the genome of WC007^T^ (Figure 3 – figure supplement 1), and two representative PULs are displayed in Figure 3A. Cellulose is a homopolymer of β-1,4-glucose and the major structural polysaccharide of the plant cell wall. These PULs are predicted to encode cellulase-like glycosylhydrolase, as well as GH5, GH30 and GH88. Cellulase is a kind of complex enzyme, mainly composed of exo-β-glucanase, endo-β-glucanase and β-glucosidase, as well as highly active xylanase. GH5, GH30 and GH88 family enzymes are respectively classified as endo-β-1,4-glucanase/cellulase, xylanase, and beta-glucuronyl hydrolase, which are all closely associated with cellulose degradation.

**(2) PULs of pectin.** Pectins are D-galacturonic acid-rich plant cell wall polysaccharides, including homogalacturonan (HG), xylogalacturonan (XGA), rhamnogalacturonan I (RG-I), rhamnogalacturonan II (RG-II) and apiogalacturonan (AGA) (**Atmodjo et al., 2013**). Eight PULs predicted to target rhamnogalacturonan and two PULs predicted to target xylogalacturonan are identified in the genome of WC007^T^ (Figure 3 – figure supplement 2), and two representative PULs are displayed in Figure 3A. These PULs are predicted to encode GH28, GH78, GH2, GH43, GH92 and GH105 family enzymes, as well as β-galactosidase enzymes and sulfatase. GH28 family enzymes are classified as α-L-rhamnosidases and often complemented by GH2, GH43 and GH105 family enzymes; GH78 family enzymes are also classified as α-L-rhamnosidases and are often complemented by GH2 and GH92 family enzymes. Thus, WC007^T^ was suggested to degrade rhamnogalacturonan and xylogalacturonan (**Bjursell et al., 2006**).

**(3) PULs of fucose-containing sulfated polysaccharides (FCSP) and L-fucoidan**. Nine PULs predicted to target FCSP are identified in the genome of WC007^T^ (Figure 3 – figure supplement 3), and two representative PULs are displayed in Figure 3A, suggesting that WC007^T^ might have a potential ability to degrade FCSP. A prominent substrate of this group is fucoidan, a highly diverse polysaccharide that is prominent in brown algae (**Ale et al., 2011**). Fucoidan is designated as a group of FCSP that contained substantial percentages of L-fucose and sulfate ester groups owing to its linear backbone of α-1,3-linked α-L-fucopyranosyl or alternating α-1,3 and 1,4-linked α-L-fucopyranosyl residues. In addition to L-fucose, different types of FCSPs also contain minor amounts of xylose, mannose, glucose, galactose and/or glucuronic acid (**Jiao et al., 2011**). In accordance with the structural complexity of FCSP, their corresponding PULs display equally complex gene repertoires (Figure 3A and Figure 3 – figure supplement 3), and these PULs contain a large amount of α-L-fucosidases, such as GH29 and GH95 family enzymes.

**(4) PULs of mannose-rich polysaccharides**. Eight PULs predicted to target mannose-rich polysaccharides are identified in the genome of WC007^T^ (Figure 3 – figure supplement 4), and two representative PULs are displayed in Figure 3A. CAZymes within these PULs are mainly classified as GH92 family enzymes and also some other family enzymes including GH2, GH31, GH76, GH95, GH125, and often associated with highly abundant sulfatase. GH92 family enzymes comprise solely α-mannosidases that cleave different linkage types (**Zhu et al., 2010**) and are predicted to degrade α-mannose-rich N-glycosylated glycoproteins, which ubiquitously exist in algae and other eukaryotes. Families GH2 and GH31 have a broader substrate spectrum, but all include β-mannosidase. GH76 (endo-α-1,6-mannanase) and GH125 (exo-α-1,6-mannanase) coexist in the same PUL. Thus, WC007^T^ is suggested to degrade sulfated α-glucuronomannan.

**(5) PULs of β-xylose-containing polysaccharides**. Eleven PULs predicted to target sulfated xylans, or xylose-rich substrates are identified in the genome of WC007^T^ (Figure 3 – figure supplement 5), and two representative PULs are displayed in Figure 3A. These PULs predicted to target xylans encode GH43, GH10, GH1, GH2, GH105, GH127 family enzymes, as well as α-galactosidase and highly abundant sulfatases. GH43 enzymes have a broader degradation capability towards mixed xylose-containing substrates such as α-L-arabinofuranosides and arabinoxylan (**Kappelmann et al., 2019**). GH10 family enzymes possess endo-β-1,4-xylanase activity, which could cleave large β-1,4-xylan backbones into oligosaccharides. GH127 and GH105 family enzymes are respectively classified as beta-L-arabinofuranosidase and unsaturated rhamnogalacturonyl hydrolase. Thus, WC007^T^ is suggested to degrade sulfated xylans and arabinoxylan.

**(6) PULs of glucans**. Three PULs predicted to target α-glucans (such as starch, glycogen starch) and β-glucans (laminarin) are identified in the genome of WC007^T^ (Figure 3 – figure supplement 6). Two respective PULs featured a SusCD-like gene triplet together with sigma -70/anti-sigma regulatory factors and at least one predicted α-glycosidase (Figure 3A). CAZymes within these PULs belong to GH97, GH3 and GH16 family enzymes. GH97 family enzymes is classified as α-glucosidase, which is known to hydrolyze various α-1,2-, α-1,3-, α-1,4-, and α-1,6-linked glycosidic bonds (**SmithSalyers, 1991**). GH3 and GH16 family enzymes, annotated as β-glucosidase, are predicted to target laminarin.

**Supplementary File 1. The BioProject and BioSample accession numbers of metagenome-assembled genomes (MAGs).**

| **SUBID** | **BioProject** | **BioSample** | **Organism** |
| --- | --- | --- | --- |
| SUB8282618 | PRJNA667764 | SAMN16383629 | Bacteroidetes bacterium C1.bin.15 |
| SUB8282685 | PRJNA667764 | SAMN16383636 | Bacteroidetes bacterium C1.bin.25 |
| SUB8282696 | PRJNA667764 | SAMN16383637 | Bacteroidetes bacterium C2.bin.15 |
| SUB8282703 | PRJNA667764 | SAMN16383723 | Bacteroidetes bacterium C2.bin.32 |
| SUB8282711 | PRJNA667764 | SAMN16383724 | Bacteroidetes bacterium C2.bin.40 |
| SUB8282714 | PRJNA667764 | SAMN16383725 | Bacteroidetes bacterium C4.bin.5 |
| SUB8282716 | PRJNA667764 | SAMN16383726 | Bacteroidetes bacterium C4.bin.18 |
| SUB8282719 | PRJNA667764 | SAMN16383730 | Bacteroidetes bacterium C4.bin.23 |
| SUB8282722 | PRJNA667788 | SAMN16383731 | Chloroflexi bacterium C1.bin.34 |
| SUB8282727 | PRJNA667788 | SAMN16383740 | Chloroflexi bacterium C1.bin.35 |
| SUB8282730 | PRJNA667788 | SAMN16383741 | Chloroflexi bacterium C2.bin.4 |
| SUB8282733 | PRJNA667788 | SAMN16383742 | Chloroflexi bacterium C2.bin.6 |
| SUB8282735 | PRJNA667788 | SAMN16383744 | Chloroflexi bacterium C2.bin.8 |
| SUB8283627 | PRJNA667788 | SAMN16384053 | Chloroflexi bacterium C2.bin.9 |
| SUB8283710 | PRJNA667788 | SAMN16384055 | Chloroflexi bacterium C2.bin.12 |
| SUB8283916 | PRJNA667788 | SAMN16384056 | Chloroflexi bacterium C2.bin.17 |
| SUB8283919 | PRJNA667788 | SAMN16384070 | Chloroflexi bacterium C2.bin.19 |
| SUB8283923 | PRJNA667788 | SAMN16384071 | Chloroflexi bacterium C2.bin.33 |
| SUB8283928 | PRJNA667788 | SAMN16384074 | Chloroflexi bacterium C2.bin.34 |
| SUB8283931 | PRJNA667788 | SAMN16384075 | Chloroflexi bacterium C2.bin.38 |
| SUB8283933 | PRJNA667788 | SAMN16384076 | Chloroflexi bacterium C2.bin.45 |
| SUB8283938 | PRJNA667788 | SAMN16384077 | Chloroflexi bacterium C2.bin.48 |
| SUB8283942 | PRJNA667811 | SAMN16384078 | Proteobacteria bacterium C1.bin.1 |
| SUB8284001 | PRJNA667811 | SAMN16384079 | Proteobacteria bacterium C1.bin.10 |
| SUB8284004 | PRJNA667811 | SAMN16384081 | Proteobacteria bacterium C1.bin.13 |
| SUB8284008 | PRJNA667811 | SAMN16384082 | Proteobacteria bacterium C1.bin.14 |
| SUB8284065 | PRJNA667811 | SAMN16384084 | Proteobacteria bacterium C1.bin.17 |
| SUB8284074 | PRJNA667811 | SAMN16385446 | Proteobacteria bacterium C1.bin.18 |
| SUB8284864 | PRJNA667811 | SAMN16386271 | Proteobacteria bacterium C1.bin.20 |
| SUB8284868 | PRJNA667811 | SAMN16386291 | Proteobacteria bacterium C1.bin.21 |
| SUB8284882 | PRJNA667811 | SAMN16386292 | Proteobacteria bacterium C1.bin.22 |
| SUB8284884 | PRJNA667811 | SAMN16386294 | Proteobacteria bacterium C1.bin.24 |
| SUB8284889 | PRJNA667811 | SAMN16386296 | Proteobacteria bacterium C1.bin.26 |
| SUB8284895 | PRJNA667811 | SAMN16386302 | Proteobacteria bacterium C1.bin.30 |
| SUB8284907 | PRJNA667811 | SAMN16386616 | Proteobacteria bacterium C1.bin.31 |
| SUB8284928 | PRJNA667811 | SAMN16386617 | Proteobacteria bacterium C1.bin.32 |
| SUB8284934 | PRJNA667811 | SAMN16386618 | Proteobacteria bacterium C1.bin.33 |
| SUB8284936 | PRJNA667811 | SAMN16386620 | Proteobacteria bacterium C1.bin.36 |
| SUB8284939 | PRJNA667811 | SAMN16386621 | Proteobacteria bacterium C1.bin.38 |
| SUB8284943 | PRJNA667811 | SAMN16386622 | Proteobacteria bacterium C1.bin.42 |
| SUB8284946 | PRJNA667811 | SAMN16386630 | Proteobacteria bacterium C2.bin.20 |
| SUB8284957 | PRJNA667811 | SAMN16386635 | Proteobacteria bacterium C2.bin.44 |
| SUB8284969 | PRJNA667811 | SAMN16386640 | Proteobacteria bacterium C4.bin.4 |
| SUB8284972 | PRJNA667811 | SAMN16386641 | Proteobacteria bacterium C4.bin.6 |
| SUB8284973 | PRJNA667811 | SAMN16386660 | Proteobacteria bacterium C4.bin.7 |
| SUB8284994 | PRJNA667811 | SAMN16387347 | Proteobacteria bacterium C4.bin.17 |
| SUB8285004 | PRJNA667811 | SAMN16387380 | Proteobacteria bacterium C4.bin.19 |
| SUB8285029 | PRJNA667811 | SAMN16387385 | Proteobacteria bacterium C4.bin.20 |
| SUB8285036 | PRJNA667811 | SAMN16387386 | Proteobacteria bacterium C4.bin.21 |
| SUB8285037 | PRJNA667811 | SAMN16387387 | Proteobacteria bacterium C4.bin.24 |
| SUB8285039 | PRJNA667811 | SAMN16387413 | Proteobacteria bacterium C4.bin.25 |

**Supplementary File 2. Assembly statistics and quality metrics of reconstructed genome bins of Proteobacteria, Bacteroidetes and Chloroflexi used in this study.**

| Bin name | Phylum | Completeness (%) | Contamination (%) | GC (%) | N50 (bp) | Genome size (Mbp) | Number of CAZymes genes |
| --- | --- | --- | --- | --- | --- | --- | --- |
| C1.bin.15 | Bacteroidetes | 59.65 | 5.172 | 31.8 | 8249 | 2.61 | 78.06 |
| C1.bin.25 | Bacteroidetes | 62.27 | 9.408 | 46.6 | 4213 | 2.67 | 114.01 |
| C2.bin.15 | Bacteroidetes | 82.06 | 3.763 | 32.8 | 5232 | 2.31 | 76.28 |
| C2.bin.32 | Bacteroidetes | 52.41 | 2.15 | 38.1 | 5636 | 1.73 | 87.92 |
| C2.bin.40 | Bacteroidetes | 89.51 | 2.867 | 36.8 | 9187 | 3.90 | 108.10 |
| C4.bin.5 | Bacteroidetes | 96.77 | 0.268 | 29.3 | 48827 | 3.75 | 77.55 |
| C4.bin.18 | Bacteroidetes | 59.08 | 0.238 | 31.4 | 29918 | 1.20 | 78.58 |
| C4.bin.23 | Bacteroidetes | 78.96 | 3.641 | 44.5 | 4832 | 2.60 | 87.34 |
| C1.bin.34 | Chloroflexi | 76.21 | 2.828 | 61.2 | 3548 | 2.59 | 87.32 |
| C1.bin.35 | Chloroflexi | 58.64 | 1.818 | 45.5 | 8245 | 1.93 | 82.22 |
| C2.bin.4 | Chloroflexi | 82.83 | 0 | 48.6 | 39431 | 0.94 | 48.86 |
| C2.bin.6 | Chloroflexi | 70.92 | 0 | 49.5 | 7628 | 0.83 | 64.06 |
| C2.bin.8 | Chloroflexi | 74.02 | 0.99 | 52.5 | 4817 | 0.76 | 70.00 |
| C2.bin.9 | Chloroflexi | 80.36 | 1.98 | 54.8 | 6764 | 1.05 | 65.62 |
| C2.bin.12 | Chloroflexi | 54.49 | 2.727 | 52.3 | 3759 | 1.64 | 72.42 |
| C2.bin.17 | Chloroflexi | 65.4 | 4.158 | 54.2 | 4882 | 0.62 | 67.61 |
| C2.bin.19 | Chloroflexi | 77.41 | 2.58 | 52.3 | 5048 | 1.98 | 63.16 |
| C2.bin.33 | Chloroflexi | 63.82 | 1.386 | 60.9 | 3494 | 1.09 | 69.44 |
| C2.bin.34 | Chloroflexi | 62.68 | 2.727 | 47.9 | 4652 | 2.21 | 91.88 |
| C4.bin.38 | Chloroflexi | 72.49 | 4.022 | 61.9 | 3598 | 2.73 | 92.75 |
| C4.bin.45 | Chloroflexi | 87.29 | 1.485 | 53.7 | 9527 | 1.65 | 72.83 |
| C2.bin.48 | Chloroflexi | 61.22 | 8.25 | 45.2 | 5264 | 0.97 | 62.63 |
| C1.bin.1 | Proteobacteria | 66.02 | 1.269 | 51.6 | 4135 | 1.88 | 75.61 |
| C1.bin.10 | Proteobacteria | 53.72 | 1.76 | 49.4 | 12409 | 1.55 | 79.13 |
| C1.bin.13 | Proteobacteria | 73.27 | 4.491 | 37 | 9298 | 2.28 | 76.60 |
| C1.bin.14 | Proteobacteria | 63.01 | 1.612 | 46.7 | 3367 | 1.97 | 74.47 |
| C1.bin.17 | Proteobacteria | 91.93 | 3.064 | 44.6 | 10510 | 2.72 | 81.02 |
| C1.bin.18 | Proteobacteria | 68.48 | 5.026 | 48.3 | 3535 | 1.99 | 66.46 |
| C1.bin.20 | Proteobacteria | 58.18 | 0.681 | 48.2 | 3398 | 1.10 | 81.75 |
| C1.bin.21 | Proteobacteria | 50.58 | 1.511 | 44.3 | 3747 | 1.54 | 63.50 |
| C1.bin.22 | Proteobacteria | 92.74 | 3.602 | 53.6 | 10690 | 2.13 | 98.26 |
| C1.bin.24 | Proteobacteria | 51.31 | 9.631 | 41.1 | 2312 | 1.76 | 55.70 |
| C1.bin.26 | Proteobacteria | 59.14 | 5.412 | 47.9 | 3308 | 3.22 | 60.00 |
| C1.bin.30 | Proteobacteria | 61.89 | 3.248 | 50.8 | 4098 | 2.26 | 72.21 |
| C1.bin.31 | Proteobacteria | 51.48 | 1.724 | 40.3 | 3199 | 0.71 | 93.57 |
| C1.bin.32 | Proteobacteria | 53.83 | 4.782 | 62.5 | 3206 | 2.94 | 63.31 |
| C1.bin.33 | Proteobacteria | 80.54 | 2.898 | 53.9 | 11265 | 3.37 | 74.76 |
| C1.bin.36 | Proteobacteria | 80.64 | 0.691 | 46.8 | 16683 | 1.90 | 91.57 |
| C1.bin.38 | Proteobacteria | 87.27 | 3.043 | 61.8 | 8879 | 3.84 | 75.77 |
| C1.bin.42 | Proteobacteria | 86.58 | 2.173 | 49 | 29878 | 2.90 | 85.77 |
| C2.bin.20 | Proteobacteria | 77.05 | 4.659 | 46.3 | 5260 | 1.90 | 91.61 |
| C2.bin.44 | Proteobacteria | 70 | 6.049 | 45.9 | 3766 | 4.65 | 78.04 |
| C4.bin. 4 | Proteobacteria | 66.03 | 2.983 | 33.2 | 2442 | 2.19 | 68.51 |
| C4.bin.6 | Proteobacteria | 60 | 3.369 | 49.2 | 3251 | 1.41 | 87.00 |
| C4.bin.7 | Proteobacteria | 77.94 | 2.728 | 49.8 | 5769 | 2.10 | 71.49 |
| C4.bin.17 | Proteobacteria | 61.2 | 1.724 | 45.5 | 6656 | 2.05 | 65.85 |
| C4.bin.19 | Proteobacteria | 67.43 | 0.565 | 64.4 | 3026 | 1.75 | 89.18 |
| C4.bin.20 | Proteobacteria | 51.05 | 5.243 | 43 | 3608 | 1.43 | 58.60 |
| C4.bin.21 | Proteobacteria | 83.47 | 2.365 | 49.8 | 5522 | 3.02 | 79.54 |
| C4.bin.24 | Proteobacteria | 50.58 | 4.257 | 40 | 1059 | 2.20 | 36.82 |
| C4.bin.25 | Proteobacteria | 64.43 | 2.258 | 46.1 | 6909 | 1.74 | 80.01 |

**Supplementary File 3. Analysis of the number of genes encoding CAZymes, SusC, SusD and sulfatase present in the 8 MAGs of deep-sea Bacteroidetes.**

| Bin name | SusC | SusD | CAZymes | sulfatases |
| --- | --- | --- | --- | --- |
| C1.bin.15 | 25 | 2 | 53 | 0 |
| C1.bin.25 | 7 | 2 | 104 | 9 |
| C2.bin.15 | 4 | 1 | 31 | 2 |
| C2.bin.32 | 5 | 2 | 35 | 3 |
| C2.bin.40 | 12 | 2 | 117 | 3 |
| C4.bin.5 | 13 | 2 | 48 | 0 |
| C4.bin.18 | 6 | 2 | 21 | 1 |
| C4.bin.23 | 8 | 0 | 59 | 1 |

**Supplementary File 4.** **Genomic characteristics of *M. comscasis* WC007^T^.**

| Characteristics | WC007^T^ | XSD2^T^ | | MCCC 1A00734^T^ | SY21^T^ | FH5^T^ | G22^T^ |
| --- | --- | --- | --- | --- | --- | --- | --- |
| Gene Bank ID | CP046401 | QWGR00000000 | JRHC00000000 | | QWET00000000 | CP007451 | FQZE00000000 |
| Genome size (bp) | 7,811,310 | 6,145,806 | | 3,426,862 | 5,826,160 | 5,073,141 | 4,779,684 |
| No.scaffolds/contigs | 1 | 354 | | 13 | 70 | 89 | 81 |
| GC-content (%) | 38.4 | 44.1 | | 40.9 | 41.7 | 41.3 | 41.8 |
| ANIb (%) | 100 | 70.11 | | 70.06 | 70.28 | 70.51 | 70.81 |
| ANIm (%)  AAI (%) | 100  100 | 84.94  71.0 | | 83.75 | 83.71  71.7 | 84.35  71.3 | 83.30 |
| Tetra | 1 | 0.92022 | | 0.94869 | 0.9496 | 0.9397 | 0.93089 |
| isDDH (%) | 100 | 20.40 | | 19.00 | 18.40 | 20.20 | 18.10 |

**Supplementary File 5.** **Differential physiological characteristics of the novel strain WC007^T^ and its closest related type strain *Maribellus luteus* XSD2^T^. Strains: 1, WC007^T^ (all data from this study); 2, *Maribellus luteus* XSD2^T^ (all data from this study except DNA G+C content and polar lipids). +, Positive result or growth; -, negative result or no growth.**

| **Characteristic** | **1** | **2** |
| --- | --- | --- |
| Cell length (µm)  Temperature range  for growth (°C)  Optimum  pH range for growth  Optimum  NaCl range for growth (%)  Optimum  Oxidase activity  Hydrolysis of:  Cellulose  Pectin  Xylan  Utilization as a sole carbon source:  Acetate  Maltose  Fructose  Ethanol  Formate  Lactate  Sorbitol  D-mannose  Major menaquinone(s)  Polar lipids  Major fatty acids (>10 %)  DNA G+C content (mol%)  Isolation source | 2.0-6.0  28-37  30  6.0-8.0  7.0  0-5  1  +  +  +  +  +  +  +  -  -  +  +  +  MK-7  PE, PL, AL, L  iso-C_15:0_, C_16:0_, summed feature 3, summed feature 8  38.38  deep-sea sediments | 2.0-9.0  20–40  28  6.0-8.5  7.0  1-5  2  -  -  -  -  -  -  -  +  +  -  -  -  MK-7  PE, AL, 3L  C16:0, C18:1*ω*9*c*, summed feature 8, C18:0  44.1  surface seawater |

**Supplementary File 6. Percentages of fatty acids useful for distinguishing** **WC007^T^ from its closest relative *Maribellus luteus* XSD2^T^. Strains: 1, WC007^T^ (all data from this study); 2, *Maribellus luteus* XSD2^T^ (all data from this study)**.

| **Fatty acid** | **Percentage (w/v) of total fatty acids**  **1 2** | |
| --- | --- | --- |
| Saturated:  C_18:0_  Branched: | 2.39 | 17.33 |
| iso-C_15:0_  C_18:1_*ω*9*c* | 14.07  7.62 | 1.45  16.57 |

**Supplementary File 7. Comparison of SusC/SusD pairs, CAZymes and sulfatases from different sources of Bacteroides.**

| **Sources** | **Strain name** | **NCBI GenBank accession number of genomes** | **SusC** | **SusD** | **GH** | **CE** | **PL** | **CBM** | **GT** | **sulfatases** | **peptidases** |
| --- | --- | --- | --- | --- | --- | --- | --- | --- | --- | --- | --- |
| **Gut** | Tidjanibacter massiliensis Marseille-P3084 | GCA_900104605.1 | 9 | 2 | 12 | 7 | 0 | 10 | 27 | 24 | 128 |
|  | Prevotellamassilia timonensis Marseille-P2831 | GCA_900106785.1 | 9 | 6 | 57 | 14 | 0 | 29 | 49 | 59 | 148 |
|  | Millionella massiliensis Marseille-P3215 | GCA_900104655.1 | 9 | 4 | 134 | 31 | 5 | 44 | 33 | 105 | 136 |
|  | Lascolabacillus massiliensis SIT8 | GCA_001282625.1 | 44 | 33 | 112 | 35 | 4 | 44 | 40 | 93 | 190 |
|  | Gabonibacter massiliensis GM7 | GCA_001487125.1 | 60 | 43 | 12 | 21 | 2 | 10 | 42 | 33 | 243 |
|  | Culturomica massiliensis Marseille-P2698 | GCA_900091655.1 | 52 | 30 | 66 | 20 | 4 | 42 | 45 | 78 | 262 |
|  | Bacteroides fragilis YCH46 | GCA_000009925.1 | 91 | 67 | 162 | 29 | 6 | 60 | 91 | 155 | 289 |
|  | Bacteroides bouchesdurhonensis Marseille-P2653 | GCA_900155865.1 | 50 | 35 | 169 | 32 | 5 | 61 | 81 | 129 | 247 |
| **Lake** | Tangfeifania diversioriginum G22 | GCA_900141875.1 | 54 | 30 | 195 | 49 | 7 | 52 | 54 | 163 | 265 |
|  | Mongoliibacter ruber YIM 4-4 | GCA_003003005.1 | 48 | 21 | 112 | 53 | 5 | 38 | 70 | 116 | 364 |
|  | Flavobacterium aciduliphilum JJ013 | GCA_003268855.1 | 17 | 3 | 28 | 23 | 2 | 21 | 49 | 48 | 176 |
|  | Belliella buryatensis 5C | GCA_900188245.1 | 34 | 15 | 67 | 27 | 7 | 24 | 53 | 78 | 276 |
|  | Aquirufa antheringensis 30S-ANTBAC | GCA_004307425.1 | 21 | 8 | 50 | 31 | 1 | 23 | 35 | 73 | 185 |
|  | Alkalitalea saponilacus SCBZ-SP2 | GCA_002201795.1 | 50 | 20 | 174 | 40 | 13 | 85 | 58 | 127 | 247 |
| **Shallow ocean** | Mariniphaga anaerophila DSM26910 | GCA_900129025.1 | 95 | 73 | 213 | 54 | 22 | 60 | 59 | 228 | 310 |
|  | Marinifilum flexuosum DSM 29150 | GCA_003610555.1 | 68 | 30 | 87 | 31 | 10 | 43 | 38 | 124 | 327 |
|  | Maribellus luteus XSD2 | GCA_003576475.1 | 113 | 83 | 277 | 53 | 12 | 94 | 46 | 258 | 402 |
|  | Luteibaculum oceani CC-AMWY-103B | GCA_007995015.1 | 12 | 2 | 13 | 14 | 0 | 49 | 55 | 61 | 193 |
|  | Lishizhenia tianjinensis H6 | GCA_900116425.1 | 7 | 1 | 30 | 22 | 0 | 29 | 53 | 69 | 212 |
|  | Labilibaculum filiforme 59.10-1M | GCA_002843315.1 | 54 | 26 | 184 | 32 | 8 | 52 | 31 | 193 | 285 |
|  | Geofilum rhodophaeum HF401 | GCA_002210225.1 | 56 | 29 | 140 | 32 | 18 | 65 | 66 | 113 | 227 |
|  | Fabibacter misakiensis SK-8 | GCA_001747105.1 | 34 | 1 | 94 | 43 | 4 | 33 | 56 | 73 | 332 |
|  | Draconibacterium orientale FH5 | GCA_000626635.1 | 70 | 39 | 219 | 59 | 5 | 73 | 66 | 187 | 292 |
|  | Catalinimonas alkaloidigena CNU-914 | GCA_900100765.1 | 99 | 60 | 217 | 71 | 18 | 140 | 83 | 214 | 462 |
|  | Brumimicrobium aurantiacum N62 | GCA_003402975.1 | 9 | 1 | 12 | 19 | 0 | 50 | 63 | 73 | 221 |
|  | Aureicoccus marinus SG-18 | GCA_002954325.1 | 27 | 5 | 79 | 23 | 3 | 24 | 47 | 44 | 233 |
|  | Ancylomarina subtilis FA102 | GCA_004217115.1 | 48 | 20 | 37 | 22 | 4 | 17 | 26 | 62 | 274 |
|  | Algoriphagus halophilus JC2051 | GCA_900129785.1 | 45 | 20 | 139 | 77 | 8 | 41 | 78 | 181 | 392 |
|  | Algoriphagus formosus XAY3209 | GCA_002807035.1 | 39 | 11 | 99 | 49 | 5 | 31 | 73 | 111 | 356 |
|  | Algoriphagus faecimaris LYX05 | GCA_900101705.1 | 36 | 14 | 98 | 50 | 6 | 44 | 74 | 109 | 356 |
|  | Algoriphagus boseongensis BS-R1 | GCA_004362805.1 | 27 | 7 | 106 | 56 | 6 | 38 | 89 | 121 | 351 |
| **Deep-sea** | Maribellus comscasis WC007T | GCA_009762775.1 | 120 | 82 | 374 | 88 | 21 | 77 | 74 | 422 | 475 |
|  | Sunxiuqinia dokdonensis DH11 | GCA_001270965.1 | 72 | 42 | 192 | 46 | 12 | 66 | 51 | 183 | 260 |
|  | Myroides profundi D25 | GCA_000833025.1 | 28 | 16 | 8 | 29 | 1 | 14 | 27 | 57 | 231 |
|  | Flammeovirga pacifica WPAGA1 | GCA_000807855.2 | 72 | 49 | 247 | 39 | 13 | 168 | 56 | 297 | 390 |
| **Others** | Pseudarcicella hirudinis E92 | GCA_900115665.1 | 96 | 63 | 182 | 70 | 11 | 72 | 72 | 179 | 399 |
|  | Flavobacterium urumqiense Sr25 | GCA_900108015.1 | 27 | 9 | 41 | 27 | 1 | 17 | 65 | 46 | 231 |
|  | Flavobacterium cheniae NJ-26 | GCA_004363695.1 | 15 | 3 | 28 | 20 | 2 | 25 | 58 | 45 | 184 |
|  | Flavobacterium anhuiense D3 | GCA_003590565.1 | 69 | 42 | 159 | 56 | 13 | 72 | 67 | 160 | 335 |
|  | Dyadobacter sediminis Z12 | GCA_005860765.1 | 98 | 64 | 137 | 71 | 7 | 109 | 93 | 197 | 408 |
|  | Cryomorpha ignava QSSC1-22 | GCA_010686655.1 | 20 | 5 | 52 | 33 | 4 | 61 | 80 | 109 | 286 |
|  | Chryseobacterium camelliae THG C4-1 | GCA_002770595.1 | 18 | 12 | 74 | 52 | 0 | 37 | 64 | 108 | 265 |
|  | Cecembia calidifontis RQ-33 | GCA_004216715.1 | 45 | 21 | 104 | 54 | 4 | 40 | 49 | 112 | 350 |
|  | Algoriphagus machipongonensis PR1 | GCA_900156785.1 | 41 | 26 | 90 | 36 | 7 | 29 | 52 | 94 | 286 |

**Supplementary File 8. Detail information of 82 PULs present in the genome of WC007^T^.**

| **Sus pair number** | **PUL composition** | **Identity** |
| --- | --- | --- |
| 1 | GM418_00650 glycoside hydrolase family 5  GM418_00655 acetylxylan esterase CE3  GM418_00660 glycoside hydrolase family 5  GM418_00665 carbohydrate binding protein with CBM9 domain  GM418_00670 sigma-70 family RNA polymerase sigma factor  GM418_00675 anti-sigma factor  GM418_00680 TonB-linked outer membrane protein, SusC/RagA family  GM418_00685 SusD/RagB Starch-binding associating with outer membrane  GM418_00690 putative endoglucanase  GM418_00695 amylo-alpha-1,6-glucosidase  GM418_00700 alginate lyase family protein  GM418_00705 glycoside hydrolase family 88 | 62%  76%  76%  66%  22%  43%  33%  42%  38%  62%  47%  17% |
| 2 | GM418_00725 sugar metabolism transcriptional regulator  GM418_00730 sigma-70 family RNA polymerase sigma factor  GM418_00735 anti-sigma factor  GM418_00740 TonB-linked outer membrane protein, SusC/RagA family  GM418_00745 SusD/RagB Starch-binding associating with outer membrane  GM418_00750 secreted glycosyl hydrolase  GM418_00755 phosphoglycerate dehydrogenase  GM418_00760 adenylosuccinate synthetase  GM418_00765 carbohydrate binding protein with CBM9 domain | 34%  27%  37%  82%  83%  61%  63%  39%  54% |
| 3 | GM418_01175 1,4-beta-xylanase  GM418_01180 xylanase  GM418_01185 arylsulfatase  GM418_01190 arylsulfatase  GM418_01195 MBL fold metallo-hydrolase  GM418_01200 hypothetical protein  GM418_01205 methyltransferase domain-containing protein  GM418_01210 glyoxalase  GM418_01215 hemolysin III family protein  GM418_01220 rhamnulokinase  GM418_01225 DUF4412 domain-containing protein  GM418_01230 catalase/peroxidase HPI  GM418_01235 bacterioferritin  GM418_01240 hypothetical protein  GM418_01245 oxidoreductase  GM418_01250 arylsulfatase  GM418_01255 phosphotransferase  GM418_01260 SusD/RagB Starch-binding associating with outer membrane  GM418_01265 TonB-linked outer membrane protein, SusC/RagA family  GM418_01270 glyoxalase  GM418_01275 alpha-1,3-galactosidase B  GM418_01280 metallophosphoesterase  GM418_01285 transglycosylase associated family protein  GM418_01290 MFS transporter  GM418_01295 Crp/FNR family transcriptional regulator  GM418_01300 hypothetical protein  GM418_01305 alpha/beta hydrolase  GM418_01310 hypothetical protein  GM418_01315 glycosylase | 75%  58%  70%  46%  62%  80%  49%  68%  41%  60%  66%  80%  31%  49%  81%  70%  42%  60%  61%  80%  66%  58%  47%  77%  19%  50%  60%  43%  72% |
| 4 | GM418_01600 adenylate cyclase  GM418_01605 hydrolase  GM418_01610 RNA polymerase sigma-70 factor  GM418_01615 anti-sigma factor  GM418_01620 TonB-linked outer membrane protein, SusC/RagA family  GM418_01625 SusD/RagB Starch-binding associating with outer membrane  GM418_01630 MFS transporter  GM418_01635 peptidyl-prolyl cis-trans isomerase | 52%  80%  20%  49%  74%  70%  77%  24% |
| 5 | GM418_01695 glycosyl hydrolase  GM418_01700 DEAD/DEAH box helicase  GM418_01705 hypothetical protein  GM418_01710 isopentenyl-diphosphate Delta-isomerase  GM418_01715 phosphoenolpyruvate carboxylase  GM418_01720 gfo/Idh/MocA family oxidoreductase  GM418_01725 RNA polymerase sigma-70 factor  GM418_01730 anti-sigma factor  GM418_01735 TonB-linked outer membrane protein, SusC/RagA family  GM418_01700 SusD/RagB Starch-binding associating with outer membrane | 62%  68%  42%  54%  26%  79%  24%  55%  59%  63% |
| 6 | GM418_01745 arylsulfatase  GM418_01750 RNA polymerase sigma-70 factor  GM418_01755 anti-sigma factor  GM418_01760 TonB-linked outer membrane protein, SusC/RagA family  GM418_01765 SusD/RagB Starch-binding associating with outer membrane  GM418_01770 xanthan lyase  GM418_01775 xanthan lyase  GM418_01780 xanthan lyase  GM418_01785 carbon starvation protein A  GM418_01790 FAD-dependent oxidoreductase  GM418_01795 6-phosphogluconolactonase  GM418_01800 glucose-6-phosphate dehydrogenase  GM418_01805 4'-phosphopantetheinyl transferase superfamily protein | 73%  25%  28%  73%  81%  72%  65%  75%  23%  75%  73%  48%  51% |
| 7 | GM418_02045 glycoside hydrolase family 97 protein  GM418_02050 RNA polymerase sigma-70 factor  GM418_02055 anti-sigma factor  GM418_02060 TonB-linked outer membrane protein, SusC/RagA family  GM418_02065 SusD/RagB Starch-binding associating with outer membrane  GM418_02070 sulfatase-like hydrolase/transferase | 48%  20%  44%  51%  23%  74% |
| 8 | GM418_02355 plasma alpha-L-fucosidase isoform X1  GM418_02360 alpha-L-fucosidase  GM418_02365 alpha-L-fucosidase  GM418_02370 alpha-L-fucosidase precursor  GM418_02375 sulfatase  GM418_02380 sulfatase-like hydrolase/transferase  GM418_02385 plasma alpha-L-fucosidase  GM418_02390 SusD/RagB Starch-binding associating with outer membrane  GM418_02395 TonB-linked outer membrane protein, SusC/RagA family  GM418_02400 anti-sigma factor  GM418_02405 RNA polymerase sigma-70 factor | 29%  62%  78%  78%  28%  59%  27%  37%  31%  68%  25% |
| 9 | GM418_02435 L-fucose:H^+^ symporter permease  GM418_02440 hypothetical protein  GM418_02445 sulfatase  GM418_02450 arylsulfatase  GM418_02455 sulfatase-like hydrolase/transferase  GM418_02460 ADP-ribosylglycohydrolase family protein  GM418_02465 heparan N-sulfatase  GM418_02470 sulfatase  GM418_02475 sulfatase  GM418_02480 arylsulfatase  GM418_02485 sulfatase-like hydrolase/transferase  GM418_02490 family 43 glycosylhydrolase  GM418_02495 hypothetical protein  GM418_02500 arylsulfatase  GM418_02505 SusD/RagB Starch-binding associating with outer membrane  GM418_02510 TonB-linked outer membrane protein, SusC/RagA family  GM418_02515 anti-sigma factor  GM418_02520 RNA polymerase sigma-70 factor | 78%  70%  66%  64%  88%  71%  74%  76%  64%  82%  59%  48%  50%  70%  32%  39%  78%  87% |
| 10 | GM418_02535 glycoside hydrolase family 92 protein  GM418_02540 sulfatase-like hydrolase/transferase  GM418_02545 SusD/RagB Starch-binding associating with outer membrane  GM418_02550 TonB-linked outer membrane protein, SusC/RagA family  GM418_02555 anti-sigma factor  GM418_02560 RNA polymerase sigma-70 factor  GM418_02565 hypothetical protein  GM418_02570 alpha-L-fucosidase  GM418_02575 diaminopimelate decarboxylase  GM418_02580 saccharopine dehydrogenase | 68%  61%  32%  37%  45%  22%  54%  72%  18%  91% |
| 11 | GM418_02635 alginate lyase family protein  GM418_02640 hypothetical protein  GM418_02645 heparinase  GM418_02650 polysaccharide lyase 8 family protein  GM418_02655 alpha/beta hydrolase  GM418_02660 lipoprotein  GM418_02665 hypothetical protein  GM418_02670 heparinase  GM418_02675 SusD/RagB Starch-binding associating with outer membrane  GM418_02680 TonB-linked outer membrane protein, SusC/RagA family  GM418_02685 Glycosyl Hydrolase Family 88  GM418_02690 acetyl-CoA hydrolase/transferase family protein  GM418_02695 acyl-CoA thioesterase  GM418_02700 acetate--CoA ligase  GM418_02705 MFS transporter | 68%  80%  72%  26%  37%  82%  65%  40%  70%  65%  59%  93%  36%  86%  77% |
| 12 | GM418_02725 GntR family transcriptional regulator  GM418_02730 MFS transporter  GM418_02735 L-arabinose isomerase  GM418_02740 TonB-linked outer membrane protein, SusC/RagA family  GM418_02745 SusD/RagB Starch-binding associating with outer membrane  GM418_02750 family 78 glycoside hydrolase catalytic domain  GM418_02755 Glycoside hydrolase family 2 sugar binding  GM418_02760 sulfatase | 32%  52%  18%  76%  59%  61%  68%  57% |
| 13 | GM418_03610 family 78 glycoside hydrolase catalytic domain  GM418_03615 glycoside hydrolase family 92 protein  GM418_03620 exo-alpha-sialidase  GM418_03625 Glycosyl transferase family 2  GM418_03630 alpha/beta hydrolase  GM418_03635 hypothetical protein  GM418_03640 glycoside hydrolase family 2  GM418_03645 polysaccharide deacetylase family protein  GM418_03650 Glycosyl hydrolases family 2, sugar binding domain  GM418_03655 family 78 glycoside hydrolase catalytic domain  GM418_03660 exo-alpha-sialidase  GM418_03665 glycerophosphodiester phosphodiesterase family protein  GM418_03670 lysoplasmalogenase  GM418_03675 hypothetical protein  GM418_03680 family 78 glycoside hydrolase catalytic domain  GM418_03685 SusD/RagB Starch-binding associating with outer membrane  GM418_03690 TonB-linked outer membrane protein, SusC/RagA family  GM418_03695 anti-sigma factor  GM418_03700 RNA polymerase sigma-70 factor | 71%  65%  67%  20%  71%  68%  70%  60%  28%  69%  19%  68%  64%  52%  71%  90%  83%  62%  23% |
| 14 | GM418_03710 glycosyl hydrolase  GM418_03715 TonB-linked outer membrane protein, SusC/RagA family  GM418_03720 SusD/RagB Starch-binding associating with outer membrane  GM418_03725 glycosyl hydrolase  GM418_03730 alpha-L-fucosidase | 39%  56%  47%  50%  81% |
| 15 | GM418_03735 glycoside hydrolase family 92 protein  GM418_03740 glycoside hydrolase family 92 protein  GM418_03745 RNA polymerase sigma-70 factor  GM418_03750 SusD/RagB Starch-binding associating with outer membrane  GM418_03755 TonB-linked outer membrane protein, SusC/RagA family  GM418_03760 anti-sigma factor | 60%  51%  26%  72%  69%  26% |
| 16 | GM418_06470 TetR family transcriptional regulator  GM418_06475 YeeE/YedE family protein  GM418_06480 magnesium transporter CorA family protein  GM418_06485 PHP domain-containing protein  GM418_06490 sulfatase-like hydrolase/transferase  GM418_06495 sulfatase-like hydrolase/transferase  GM418_06500 SusD/RagB Starch-binding associating with outer membrane  GM418_06505 TonB-linked outer membrane protein, SusC/RagA family  GM418_06510 anti-sigma factor  GM418_06515 RNA polymerase sigma-70 factor  GM418_06520 serine hydrolase  GM418_06525 GNAT family N-acetyltransferase | 75%  19%  77%  73%  81%  60%  54%  52%  43%  19%  65%  56% |
| 17 | GM418_06565 Lytic transglycosylase  GM418_06570 TonB-linked outer membrane protein, SusC/RagA family  GM418_06575 SusD/RagB Starch-binding associating with outer membrane  GM418_06580 glycerophosphodiester phosphodiesterase  GM418_06585 histidinol-phosphatase  GM418_06590 DNA-3-methyladenine glycosylase 2 family protein  GM418_06595 hypothetical protein  GM418_06600 hypothetical protein  GM418_06605 TonB-dependent receptor  GM418_06610 M56 family metallopeptidase  GM418_06615 BlaI/MecI/CopY family transcriptional regulator  GM418_06620 glycosyl hydrolase  GM418_06625 family 78 glycoside hydrolase catalytic domain | 66%  85%  80%  37%  65%  44%  41%  56%  56%  36%  26%  79%  62% |
| 18 | GM418_07185 membrane lipoprotein  GM418_07190 Gfo/Idh/MocA family oxidoreductase  GM418_07195 RNA polymerase sigma-70 factor  GM418_07200 anti-sigma factor  GM418_07205 TonB-linked outer membrane protein, SusC/RagA family  GM418_07210 SusD/RagB Starch-binding associating with outer membrane  GM418_07215 plasma alpha-L-fucosidase  GM418_07220 family 10 glycosylhydrolase  GM418_07225 glycoside hydrolase family 10 protein  GM418_07230 alpha-L-fucosidase  GM418_07235 sugar phosphate isomerase/epimerase  GM418_07240 plasma alpha-L-fucosidase | 67%  84%  67%  28%  65%  72%  34%  18%  18%  87%  81%  21% |
| 19 | GM418_07250 methyltransferase  GM418_07255 L-rhamnose-proton symporter  GM418_07260 glycoside hydrolase, family 29  GM418_07265 RNA polymerase sigma-70 factor  GM418_07270 anti-sigma factor  GM418_07275 TonB-linked outer membrane protein, SusC/RagA family  GM418_07280 SusD/RagB Starch-binding associating with outer membrane  GM418_07285 sulfatase | 76%  29%  57%  22%  47%  55%  62%  73% |
| 20 | GM418_07295 glycosyl hydrolase  GM418_07300 IS1380 family transposase  GM418_07305 glycerophosphodiester phosphodiesterase  GM418_07310 TonB-linked outer membrane protein, SusC/RagA family  GM418_07315 SusD/RagB Starch-binding associating with outer membrane | 46%  17%  46%  78%  84% |
| 21 | GM418_07325 RNA polymerase sigma-70 factor  GM418_07330 anti-sigma factor  GM418_07335 TonB-linked outer membrane protein, SusC/RagA family  GM418_07340 SusD/RagB Starch-binding associating with outer membrane  GM418_07345 N-acetylgalactosamine 6-sulfate sulfatase  GM418_07350 alpha-L-fucosidase precursor  GM418_07355 sulfatase-like hydrolase/transferase  GM418_07360 sulfatase-like hydrolase/transferase | 21%  32%  66%  74%  68%  88%  74%  68% |
| 22 | GM418_07365 alpha-L-fucosidase  GM418_07370 RNA polymerase sigma-70 factor  GM418_07375 anti-sigma factor  GM418_07380 TonB-linked outer membrane protein, SusC/RagA family  GM418_07385 SusD/RagB Starch-binding associating with outer membrane  GM418_07390 sulfatase  GM418_07395 sulfatase-like hydrolase/transferase  GM418_07400 sulfatase | 72%  24%  40%  40%  59%  73%  79%  67% |
| 23 | GM418_07415 family 78 glycoside hydrolase catalytic domain  GM418_07420 family 78 glycoside hydrolase catalytic domain  GM418_07425 family 78 glycoside hydrolase catalytic domain  GM418_07430 L-arabinose isomerase  GM418_07435 RNA polymerase sigma-70 factor  GM418_07440 IS4 family transposase  GM418_07445 anti-sigma factor  GM418_07450 TonB-linked outer membrane protein, SusC/RagA family  GM418_07455 SusD/RagB Starch-binding associating with outer membrane  GM418_07460 MBL fold metallo-hydrolase  GM418_07465 sulfatase  GM418_07470 MFS transporter  GM418_07475 sulfatase | 68%  73%  69%  17%  25%  60%  56%  54%  70%  55%  74%  76%  71% |
| 24 | GM418_07505 family 20 glycosylhydrolase  GM418_07510 glycosyl hydrolase  GM418_07515 RNA polymerase sigma-70 factor  GM418_07520 anti-sigma factor  GM418_07525 TonB-linked outer membrane protein, SusC/RagA family  GM418_07530 SusD/RagB Starch-binding associating with outer membrane  GM418_07535 sulfatase  GM418_07540 sulfatase  GM418_07545 sulfatase  GM418_07550 sulfatase  GM418_07555 ankyrin repeat domain-containing protein  GM418_07560 MFS transporter  GM418_07565 methyltransferase  GM418_07570 family 10 glycosylhydrolase  GM418_07575 plasma alpha-L-fucosidase | 72%  38%  24%  41%  73%  76%  81%  70%  80%  79%  39%  54%  80%  25%  41% |
| 25 | GM418_08200 Gfo/Idh/MocA family oxidoreductase  GM418_08205 TonB-linked outer membrane protein, SusC/RagA family  GM418_08210 SusD/RagB Starch-binding associating with outer membrane  GM418_08215 DUF2264 domain-containing protein  GM418_08220 glycosyl hydrolase, family 88  GM418_08225 oxidoreductase  GM418_08230 M28 family metallohydrolase  GM418_08235 M48 family metalloprotease  GM418_08240 glycoside hydrolase family 5 protein | 85%  72%  58%  85%  28%  83%  50%  68%  79% |
| 26 | GM418_08290 methyltransferase  GM418_08295 peptidase M16  GM418_08300 RNA polymerase sigma-70 factor  GM418_08305 anti-sigma factor  GM418_08310 TonB-linked outer membrane protein, SusC/RagA family  GM418_08315 SusD/RagB Starch-binding associating with outer membrane  GM418_08320 SusE outer membrane protein  GM418_08325 glycosyl hydrolase family 76  GM418_08330 glycoside hydrolase family 92 protein  GM418_08335 glycoside hydrolase family 125 protein | 54%  33%  21%  29%  74%  79%  58%  77%  79%  82% |
| 27 | GM418_08350 family 43 glycosylhydrolase  GM418_08355 family 43 glycosylhydrolase  GM418_08360 family 43 glycosylhydrolase  GM418_08365 family 43 glycosylhydrolase  GM418_08370 alpha-galactosidase  GM418_08375 RNA polymerase sigma-70 factor  GM418_08380 anti-sigma factor  GM418_08385 TonB-linked outer membrane protein, SusC/RagA family  GM418_08390 SusD/RagB Starch-binding associating with outer membrane  GM418_08395 arylsulfatase  GM418_08400 family 43 glycosylhydrolase  GM418_08405 sulfatase-like hydrolase/transferase  GM418_08410 family 43 glycosylhydrolase | 75%  69%  59%  79%  30%  26%  62%  42%  38%  58%  30%  48%  48% |
| 28 | GM418_08415 RNA polymerase sigma-70 factor  GM418_08420 anti-sigma factor  GM418_08425 TonB-linked outer membrane protein, SusC/RagA family  GM418_08430 SusD/RagB Starch-binding associating with outer membrane  GM418_08435 amylo-alpha-1,6-glucosidase  GM418_08440 glycoside hydrolase family 127 protein  GM418_08445 Beta-L-arabinofuranosidase, GH127  GM418_08450 beta-galactosidase  GM418_08455 glycerate dehydrogenase | 17%  32%  71%  81%  51%  69%  55%  78%  53% |
| 29 | GM418_09245 RNA polymerase sigma-70 factor  GM418_09250 anti-sigma factor  GM418_09255 TonB-linked outer membrane protein, SusC/RagA family  GM418_09260 SusD/RagB Starch-binding associating with outer membrane  GM418_09265 glycoside hydrolase family 27 protein  GM418_09270 arylsulfatase  GM418_09275 sulfatase  GM418_09280 glycerophosphodiester phosphodiesterase  GM418_09285 sulfatase  GM418_09290 carbohydrate binding family 9 domain-containing protein  GM418_09295 L-lactate dehydrogenase | 25%  58%  62%  64%  69%  78%  76%  23%  79%  74%  32% |
| 30 | GM418_09905 nucleoside hydrolase  GM418_09910 nucleoside hydrolase  GM418_09915 sulfatase  GM418_09920 SusD/RagB Starch-binding associating with outer membrane  GM418_09925 TonB-linked outer membrane protein, SusC/RagA family  GM418_09930 anti-sigma factor  GM418_09935 RNA polymerase sigma-70 factor  GM418_09940 HipA domain-containing protein  GM418_09945 type II toxin-antitoxin system HipA family toxin  GM418_09950 helix-turn-helix transcriptional regulator  GM418_09955 hypothetical protein  GM418_09960 glycoside hydrolase family 28 protein  GM418_09965 rhamnogalacturonan acetylesterase  GM418_09970 family 43 glycosylhydrolase  GM418_09975 glycoside hydrolase family 105 protein  GM418_09980 prolyl oligopeptidase family serine peptidase | 62%  23%  58%  49%  46%  56%  24%  83%  30%  30%  39%  70%  70%  63%  33%  67% |
| 31 | GM418_10245 cellulase family glycosylhydrolase  GM418_10250 glycoside hydrolase family 30 protein  GM418_10255 beta-galactosidase domain protein  GM418_10260 beta-galactosidase  GM418_10265 threonine synthase  GM418_10270 RNA polymerase sigma-70 factor  GM418_10275 anti-sigma factor  GM418_10280 TonB-linked outer membrane protein, SusC/RagA family  GM418_10285 SusD/RagB Starch-binding associating with outer membrane | 46%  67%  53%  35%  20%  26%  53%  59%  71% |
| 32 | GM418_10515 TonB-linked outer membrane protein, SusC/RagA family  GM418_10520 SusD/RagB Starch-binding associating with outer membrane  GM418_10525 Surface glycan-binding protein B xyloglucan binding domain  GM418_10530 Polygalacturonase  GM418_10535 beta-mannosidase | 54%  70%  29%  28%  59% |
| 33 | GM418_10540 RNA polymerase sigma-70 factor  GM418_10545 anti-sigma factor  GM418_10550 TonB-linked outer membrane protein, SusC/RagA family  GM418_10555 SusD/RagB Starch-binding associating with outer membrane  GM418_10560 sulfatase-like hydrolase/transferase  GM418_10565 sulfatase-like hydrolase/transferase  GM418_10570 sulfatase-like hydrolase/transferase  GM418_10575 arylsulfatase  GM418_10580 sulfatase-like hydrolase/transferase | 21%  49%  71%  81%  65%  73%  49%  62%  51% |
| 34 | GM418_10705 indole-3-glycerol phosphate synthase TrpC  GM418_10710 phosphoribosylanthranilate isomerase  GM418_10715 tryptophan synthase subunit beta  GM418_10720 tryptophan synthase subunit alpha  GM418_10725 hypothetical protein  GM418_10730 TonB-linked outer membrane protein, SusC/RagA family  GM418_10735 SusD/RagB Starch-binding associating with outer membrane | 41%  33%  92%  39%  91%  81%  80% |
| 35 | GM418_10745 TonB-linked outer membrane protein, SusC/RagA family  GM418_10750 SusD/RagB Starch-binding associating with outer membrane  GM418_10755 peptidylprolyl isomerase  GM418_10760 aminopeptidase  GM418_10765 alpha-galactosidase  GM418_10766 Gfo/Idh/MocA family oxidoreductase | 60%  48%  60%  62%  45%  83% |
| 36 | GM418_11575 SusD/RagB Starch-binding associating with outer membrane  GM418_11580 TonB-linked outer membrane protein, SusC/RagA family  GM418_11585 4-hydroxy-tetrahydrodipicolinate synthase  GM418_11590 GntR family transcriptional regulator  GM418_11595 assimilatory sulfite reductase (NADPH) flavoprotein subunit  GM418_11600 PepSY domain-containing protein  GM418_11605 TonB-dependent receptor  GM418_11610 beta-mannosidase | 49%  57%  25%  22%  39%  62%  65%  66% |
| 37 | GM418_11690 response regulator  GM418_11695 TonB-linked outer membrane protein, SusC/RagA family  GM418_11700 SusD/RagB Starch-binding associating with outer membrane  GM418_11705 alpha-N-arabinofuranosidase  GM418_11710 sulfatase  GM418_11715 beta-lactamase  GM418_11720 sulfatase-like hydrolase/transferase  GM418_11725 glycoside hydrolase family 2 protein  GM418_11730 arylsulfatase  GM418_11735 arylsulfatase | 54%  38%  47%  19%  53%  39%  73%  80%  67%  74% |
| 38 | GM418_13200 polysulfide reductase  GM418_13205 PKD domain-containing protein  GM418_13210 DUF4832 domain-containing protein  GM418_13215 SusD/RagB Starch-binding associating with outer membrane  GM418_13220 TonB-linked outer membrane protein, SusC/RagA family  GM418_13225 anti-sigma factor  GM418_13230 RNA polymerase sigma-70 factor  GM418_13235 methylmalonyl-CoA mutase  GM418_13240 four helix bundle protein  GM418_13245 methylmalonyl-CoA mutase  GM418_13250 glycosyl hydrolase  GM418_13255 hypothetical protein  GM418_13260 thioredoxin-dependent thiol peroxidase  GM418_13265 mannose-1-phosphate guanylyltransferase | 84%  50%  40%  66%  66%  52%  18%  26%  44%  83%  33%  48%  22%  77% |
| 39 | GM418_13460 metallophosphatase  GM418_13465 glycerophosphodiester phosphodiesterase family protein  GM418_13470 SusD/RagB Starch-binding associating with outer membrane  GM418_13475 TonB-linked outer membrane protein, SusC/RagA family  GM418_13480 anti-sigma factor  GM418_13485 RNA polymerase sigma-70 factor  GM418_13490 DNA alkylation repair protein  GM418_13495 ABC transporter permease  GM418_13500 aminopeptidase P family protein  GM418_13505 FtsX-like permease family protein | 70%  59%  55%  54%  26%  24%  20%  19%  21%  69% |
| 40 | GM418_14150 SGNH/GDSL hydrolase family protein  GM418_14155 RNA polymerase sigma-70 factor  GM418_14160 anti-sigma factor  GM418_14165 TonB-linked outer membrane protein, SusC/RagA family  GM418_14170 SusD/RagB Starch-binding associating with outer membrane  GM418_14175 PKD domain-containing protein | 55%  23%  53%  23%  48%  39% |
| 41 | GM418_14190 glycoside hydrolase family 105 protein  GM418_14195 histidine kinase  GM418_14200 TonB-linked outer membrane protein, SusC/RagA family  GM418_14205 SusD/RagB Starch-binding associating with outer membrane  GM418_14210 arylsulfatase  GM418_14215 sulfatase  GM418_14220 family 10 glycosylhydrolase  GM418_14225 sulfatase  GM418_14230 sulfatase | 22%  38%  70%  71%  64%  64%  32%  69%  47% |
| 42 | GM418_14280 alpha-L-arabinofuranosidase  GM418_14285 family 43 glycosylhydrolase  GM418_14290 beta-galactosidase  GM418_14295 glycoside hydrolase family 127 protein  GM418_14300 RNA polymerase sigma-70 factor  GM418_14305 hypothetical protein  GM418_14310 anti-sigma factor  GM418_14315 TonB-linked outer membrane protein, SusC/RagA family  GM418_14320 SusD/RagB Starch-binding associating with outer membrane  GM418_14325 family 43 glycosylhydrolase | 64%  79%  24%  74%  21%  46%  54%  80%  78%  67% |
| 43 | GM418_15400 MFS transporter  GM418_15405 amidase  GM418_15410 DUF4922 domain-containing protein  GM418_15415 glycosyltransferase family 2 protein  GM418_15420 alpha-L-fucosidase  GM418_15425 RNA polymerase sigma-70 factor  GM418_15430 anti-sigma factor  GM418_15435 TonB-linked outer membrane protein, SusC/RagA family  GM418_15440 SusD/RagB Starch-binding associating with outer membrane  GM418_15445 sulfatase  GM418_15450 arylsulfatase | 65%  67%  59%  74%  73%  19%  19%  71%  70%  70%  79% |
| 44 | GM418_15460 sulfatase-like hydrolase/transferase  GM418_15465 RNA polymerase sigma-70 factor  GM418_15470 anti-sigma factor  GM418_15475 TonB-linked outer membrane protein, SusC/RagA family  GM418_15480 SusD/RagB Starch-binding associating with outer membrane  GM418_15485 sulfatase  GM418_15490 family 10 glycosylhydrolase | 65%  23%  44%  45%  54%  59%  72% |
| 45 | GM418_15570 Gfo/Idh/MocA family oxidoreductase  GM418_15575 SusD/RagB Starch-binding associating with outer membrane  GM418_15580 TonB-linked outer membrane protein, SusC/RagA family  GM418_15585 Polyprenol monophosphomannose synthase  GM418_15590 sigma 54-interacting transcriptional regulator | 88%  63%  56%  66%  76% |
| 46 | GM418_15595 GNAT family N-acetyltransferase  GM418_15600 RNA polymerase sigma-70 factor  GM418_15605 anti-sigma factor  GM418_15610 TonB-linked outer membrane protein, SusC/RagA family  GM418_15615 SusD/RagB Starch-binding associating with outer membrane  GM418_15620 Gfo/Idh/MocA family oxidoreductase  GM418_15625 PIG-L family deacetylase  GM418_15630 DUF1080 domain-containing protein  GM418_15635 sugar phosphate isomerase/epimerase  GM418_15640 Gfo/Idh/MocA family oxidoreductase  GM418_15645 sugar phosphate isomerase/epimerase | 70%  23%  69%  77%  78%  87%  76%  83%  61%  94%  87% |
| 47 | GM418_15795 alpha-xylosidase  GM418_15800 aryl-phospho-beta-D-glucosidase BglC (GH1 family)  GM418_15805 hypothetical protein  GM418_15810 SusD/RagB Starch-binding associating with outer membrane  GM418_15815 TonB-linked outer membrane protein, SusC/RagA family | 61%  57%  57%  70%  77% |
| 48 | GM418_16150 glycosyl hydrolase  GM418_16155 gliding motility lipoprotein GldB  GM418_16160 NAD+ synthase  GM418_16165 TonB-linked outer membrane protein, SusC/RagA family  GM418_16170 SusD/RagB Starch-binding associating with outer membrane  GM418_16175 cAMP-activated global transcriptional regulator CRP  GM418_16180 alanine racemase | 67%  63%  82%  70%  66%  17%  46% |
| 49 | GM418_16200 phosphoglycolate phosphatase  GM418_16205 TonB-linked outer membrane protein, SusC/RagA family  GM418_16210 SusD/RagB Starch-binding associating with outer membrane  GM418_16215 hypothetical protein  GM418_16220 BlaI/MecI/CopY family transcriptional regulator | 20%  70%  65%  59%  24% |
| 50 | GM418_16465 alpha/beta hydrolase  GM418_16470 alkaline phosphatase  GM418_16475 arylsulfatase  GM418_16480 alkaline phosphatase family protein  GM418_16485 sulfatase-like hydrolase/transferase  GM418_16490 SusD/RagB Starch-binding associating with outer membrane  GM418_16495 TonB-linked outer membrane protein, SusC/RagA family  GM418_16500 anti-sigma factor  GM418_16505 RNA polymerase sigma-70 factor  GM418_16510 Glycosyl hydrolases family 31  GM418_16515 cellulase family glycosylhydrolase  GM418_16520 alpha-mannosidase | 62%  75%  76%  67%  68%  59%  48%  38%  23%  70%  25%  28% |
| 51 | GM418_17555 GNAT family N-acetyltransferase  GM418_17560 glycosyl transferase family 2  GM418_17565 recombination protein RecR  GM418_17570 Gfo/Idh/MocA family oxidoreductase  GM418_17575 NADP oxaloacetate-decarboxylating malate dehydrogenase  GM418_17580 RNA polymerase sigma-70 factor  GM418_17585 anti-sigma factor  GM418_17590 TonB-linked outer membrane protein, SusC/RagA family  GM418_17595 SusD/RagB Starch-binding associating with outer membrane  GM418_17600 sulfatase  GM418_17605 sulfatase  GM418_17610 polyketide cyclase | 21%  69%  45%  85%  47%  23%  35%  64%  71%  51%  80%  17% |
| 52 | GM418_18550 M3 family peptidase  GM418_18555 mandelate racemase/muconate lactonizing enzyme family protein  GM418_18560 bile acid:sodium symporter family protein  GM418_18565 dimethylmenaquinone methyltransferase  GM418_18570 RNA polymerase sigma-70 factor  GM418_18575 anti-sigma factor  GM418_18580 TonB-linked outer membrane protein, SusC/RagA family  GM418_18585 SusD/RagB Starch-binding associating with outer membrane  GM418_18590 family 16 glycosylhydrolase | 79%  62%  66%  62%  21%  50%  61%  70%  54% |
| 53 | GM418_18610 AraC family transcriptional regulator  GM418_18615 TonB-linked outer membrane protein, SusC/RagA family  GM418_18620 SusD/RagB Starch-binding associating with outer membrane  GM418_18625 hypothetical protein  GM418_18630 cytochrome C biosynthesis protein | 66%  77%  73%  44%  52% |
| 54 | GM418_18675 RNA polymerase sigma-70 factor  GM418_18680 anti-sigma factor  GM418_18685 TonB-linked outer membrane protein, SusC/RagA family  GM418_18690 SusD/RagB Starch-binding associating with outer membrane  GM418_18695 glycoside hydrolase family 92 protein  GM418_18700 endo-1,4-beta-xylanase  GM418_18705 family 78 glycoside hydrolase catalytic domain  GM418_18710 glycoside hydrolase family 16 protein  GM418_18715 sulfatase  GM418_18720 sulfatase  GM418_18725 hypothetical protein  GM418_18730 family 43 glycosylhydrolase | 72%  27%  56%  52%  66%  54%  71%  31%  47%  68%  74%  65% |
| 55 | GM418_19200 GNAT family N-acetyltransferase  GM418_19205 DMT family transporter  GM418_19210 RNA polymerase sigma-70 factor  GM418_19215 anti-sigma factor  GM418_19220 TonB-linked outer membrane protein, SusC/RagA family  GM418_19225 SusD/RagB Starch-binding associating with outer membrane  GM418_19230 argininosuccinate lyase  GM418_19235 2-methylcitrate dehydratase  GM418_19240 sensor histidine kinase  GM418_19245 response regulator  GM418_19250 glycoside hydrolase family 3 protein | 47%  27%  83%  65%  59%  60  26%  15%  58%  84%  83% |
| 56 | GM418_21520 family 20 glycosylhydrolase  GM418_21525 glycosyl hydrolases family 31  GM418_21530 SusD/RagB Starch-binding associating with outer membrane  GM418_21535 TonB-linked outer membrane protein, SusC/RagA family  GM418_21540 anti-sigma factor  GM418_21545 RNA polymerase sigma-70 factor | 59%  53%  56%  38%  33%  92% |
| 57 | GM418_21570 RNA polymerase sigma-70 factor  GM418_21575 anti-sigma factor  GM418_21580 TonB-linked outer membrane protein, SusC/RagA family  GM418_21585 SusD/RagB Starch-binding associating with outer membrane  GM418_21590 hypothetical protein  GM418_21595 family 10 glycosylhydrolase  GM418_21600 arylsulfatase  GM418_21605 sulfatase | 21%  80%  50%  58%  72%  34%  82%  50% |
| 58 | GM418_21865 arylsulfatase  GM418_21870 sulfatase  GM418_21875 SusD/RagB Starch-binding associating with outer membrane  GM418_21880 TonB-linked outer membrane protein, SusC/RagA family  GM418_21885 anti-sigma factor  GM418_21890 RNA polymerase sigma-70 factor  GM418_21895 DNA mismatch repair protein MutS  GM418_21900 TIGR01777 family protein | 69%  65%  63%  57%  53%  22%  45%  36% |
| 59 | GM418_22540 alpha-L-fucosidase  GM418_22545 copper homeostasis protein CutC  GM418_22550 tRNA (adenosine(37)-N6)-dimethylallyltransferase MiaA  GM418_22555 threonine/serine exporter family protein  GM418_22560 threonine/serine exporter  GM418_22565 acyl-ACP thioesterase  GM418_22570 type I methionyl aminopeptidase  GM418_22575 hypothetical protein  GM418_22580 RNA polymerase sigma-70 factor  GM418_22585 anti-sigma factor  GM418_22590 TonB-linked outer membrane protein, SusC/RagA family  GM418_22595 SusD/RagB Starch-binding associating with outer membrane  GM418_22600 sulfatase | 82%  32%  31%  24%  85%  23%  51%  44%  22%  34%  50%  61%  76% |
| 60 | GM418_24200 beta-galactosidase  GM418_24205 PAS domain S-box protein  GM418_24210 family 78 glycoside hydrolase catalytic domain  GM418_24215 acetylxylan esterase CE3  GM418_24220 SusD/RagB Starch-binding associating with outer membrane  GM418_24225 TonB-linked outer membrane protein, SusC/RagA family  GM418_24230 anti-sigma factor  GM418_24235 RNA polymerase sigma-70 factor | 35%  28%  85%  79%  28%  43%  32%  20% |
| 61 | GM418_24280 acyltransferase  GM418_24285 agmatine deiminase  GM418_24290 RNA polymerase sigma-70 factor  GM418_24295 anti-sigma factor  GM418_24300 TonB-linked outer membrane protein, SusC/RagA family  GM418_24305 SusD/RagB Starch-binding associating with outer membrane  GM418_24310 TetR/AcrR family transcriptional regulator | 67%  30%  21%  52%  79%  80%  17% |
| 62 | GM418_24875 arylsulfatase  GM418_24880 arylsulfatase  GM418_24885 glycoside hydrolase family 2  GM418_24890 sulfatase  GM418_24895 glycosyl hydrolase family 43  GM418_24900 SusD/RagB Starch-binding associating with outer membrane  GM418_24905 TonB-linked outer membrane protein, SusC/RagA family  GM418_24910 family 43 glycosylhydrolase | 83%  81%  71%  75%  36%  87%  87%  82% |
| 63 | GM418_24970 TonB-linked outer membrane protein, SusC/RagA family  GM418_24975 SusD/RagB Starch-binding associating with outer membrane  GM418_24980 glycoside hydrolase family 2  GM418_24985 hypothetical protein  GM418_24990 sulfatase  GM418_24995 arylsulfatase  GM418_25000 aminopeptidase  GM418_25005 glycoside hydrolase family 31 protein  GM418_25010 glycoside hydrolase family 95 protein  GM418_25015 sulfatase-like hydrolase/transferase  GM418_25020 S9 family peptidase  GM418_25025 alanine dehydrogenase  GM418_25030 L-serine ammonia-lyase  GM418_25035 glycoside hydrolase family 92 protein  GM418_25040 glycoside hydrolase family 2  GM418_25045 sugar MFS transporter  GM418_25050 hypothetical protein  GM418_25055 glycoside hydrolase family 92 protein  GM418_25060 hypothetical protein  GM418_25065 phosphoesterase  GM418_25070 glycoside hydrolase family 92 protein | 67%  61%  23%  95%  68%  67%  69%  82%  73%  71%  77%  35%  40%  76%  73%  59%  63%  71%  50%  91%  72% |
| 64 | GM418_25775 sulfatase  GM418_25780 sulfatase  GM418_25785 Gfo/Idh/MocA family oxidoreductase  GM418_25790 methane monooxygenase PmoA-like  GM418_25795 sulfatase  GM418_25800 SusD/RagB Starch-binding associating with outer membrane  GM418_25805 TonB-linked outer membrane protein, SusC/RagA family  GM418_25810 anti-sigma factor  GM418_25815 RNA polymerase sigma-70 factor  GM418_25820 glycosyl hydrolase family 88 | 56%  70%  75%  49%  64%  31%  35%  72%  22%  60% |
| 65 | GM418_25830 RNA polymerase sigma-70 factor  GM418_25835 anti-sigma factor  GM418_25840 TonB-linked outer membrane protein, SusC/RagA family  GM418_25845 SusD/RagB Starch-binding associating with outer membrane  GM418_25850 FAD-dependent oxidoreductase  GM418_25855 alpha-L-fucosidase  GM418_25860 6-bladed beta-propeller  GM418_25865 3-oxoacyl-[acyl-carrier-protein] reductase  GM418_25870 hypothetical protein  GM418_25875 3-oxoacyl-[acyl-carrier-protein] reductase  GM418_25880 deoxyribose-phosphate aldolase  GM418_25885 alcohol dehydrogenase  GM418_25890 3-oxoacyl-[acyl-carrier-protein] reductase  GM418_25895 xylulokinase  GM418_25900 arylsulfatase | 18%  56%  37%  55%  22%  48%  18%  30%  30%  36%  50%  37%  41%  30%  61% |
| 66 | GM418_26085 2-keto-3-deoxy-L-rhamnonate aldolase  GM418_26090 family 78 glycoside hydrolase catalytic domain  GM418_26095 DUF1349 domain-containing protein  GM418_26100 regulator  GM418_26105 metallophosphoesterase  GM418_26110 RNA polymerase sigma-70 factor  GM418_26115 anti-sigma factor  GM418_26120 hypothetical protein  GM418_26125 TonB-linked outer membrane protein, SusC/RagA family  GM418_26130 SusD/RagB Starch-binding associating with outer membrane | 28%  35%  62%  68%  71%  24%  23%  35%  76%  88% |
| 67 | GM418_26935 cellulase family glycosylhydrolase  GM418_26940 peptidase M16  GM418_26945 hypothetical protein  GM418_26950 thioredoxin  GM418_26955 RNA polymerase sigma-70 factor  GM418_26960 anti-sigma factor  GM418_26965 TonB-linked outer membrane protein, SusC/RagA family  GM418_26970 SusD/RagB Starch-binding associating with outer membrane | 49%  59%  73%  43%  56%  51%  57%  67% |
| 68 | GM418_26995 methyltransferase domain-containing protein  GM418_27000 TonB-dependent receptor  GM418_27005 RNA polymerase sigma-70 factor  GM418_27010 anti-sigma factor  GM418_27015 TonB-linked outer membrane protein, SusC/RagA family  GM418_27020 SusD/RagB Starch-binding associating with outer membrane  GM418_27025 T9SS C-terminal target domain-containing protein  GM418_27030 Pectate lyase L  GM418_27035 plasma alpha-L-fucosidase  GM418_27040 xylan esterase  GM418_27045 L1_beta_lactamase | 62%  83%  24%  38%  64%  70%  32%  30%  25%  49%  25% |
| 69 | GM418_27125 RNA polymerase sigma-70 factor  GM418_27130 anti-sigma factor  GM418_27135 TonB-linked outer membrane protein, SusC/RagA family  GM418_27140 SusD/RagB Starch-binding associating with outer membrane  GM418_27145 glycosyl hydrolase  GM418_27150 Gfo/Idh/MocA family oxidoreductase  GM418_27155 alpha-L-fucosidase  GM418_27160 ADP-ribosylglycohydrolase family protein  GM418_27165 right-handed parallel beta-helix repeat-containing protein  GM418_27170 carbohydrate-binding family 6 protein  GM418_27175 arylsulfatase  GM418_27180 family 43 glycosylhydrolase  GM418_27185 MFS transporter | 22%  50%  38%  50%  40%  71%  67%  60%  46%  29%  62%  39%  16% |
| 70 | GM418_27605 metallophosphoesterase family protein  GM418_27610 hypothetical protein  GM418_27615 DUF2961 domain-containing protein  GM418_27620 glycoside hydrolase family 28 protein  GM418_27625 hypothetical protein  GM418_27630 hypothetical protein  GM418_27635 DUF3748 domain-containing protein  GM418_27640 metallophosphoesterase  GM418_27645 SusD/RagB Starch-binding associating with outer membrane  GM418_27650 TonB-linked outer membrane protein, SusC/RagA family | 68%  70%  67%  67%  71%  59%  64%  60%  82%  88% |
| 71 | GM418_27670 TonB-linked outer membrane protein, SusC/RagA family  GM418_27675 SusD/RagB Starch-binding associating with outer membrane  GM418_27680 PhoPQ-activated pathogenicity-like protein PqaA type  GM418_27685 hypothetical protein  GM418_27690 response regulator transcription factor  GM418_27695 histidine kinase | 75%  75%  82%  80%  62%  58% |
| 72 | GM418_27965 M20/M25/M40 family metallo-hydrolase  GM418_27970 3-oxoacyl-[acyl-carrier-protein] reductase  GM418_27975 hypothetical protein  GM418_27980 SusD/RagB Starch-binding associating with outer membrane  GM418_27985 TonB-linked outer membrane protein, SusC/RagA family  GM418_27990 PIG-L family deacetylase  GM418_27995 hypothetical protein  GM418_28000 DUF2961 domain-containing protein  GM418_28005 DNA polymerase III subunit beta  GM418_28010 arabinogalactan endo-1,4-beta-galactosidase  GM418_28015 Gfo/Idh/MocA family oxidoreductase  GM418_28020 Sulfatase | 36%  16%  48%  72%  68%  73%  44%  61%  24%  36%  81%  72% |
| 73 | GM418_28235 glycoside hydrolase family 2  GM418_28240 glycoside hydrolase family 2 protein  GM418_28245 family 43 glycosylhydrolase  GM418_28250 glycoside hydrolase family 28 protein  GM418_28255 glycoside hydrolase family 28 protein  GM418_28260 family 43 glycosylhydrolase  GM418_28265 SusD/RagB Starch-binding associating with outer membrane  GM418_28270 TonB-linked outer membrane protein, SusC/RagA family  GM418_28275 anti-sigma factor  GM418_28280 RNA polymerase sigma-70 factor | 81%  67%  65%  73%  63%  66%  78%  77%  53%  19% |
| 74 | GM418_28810 family 43 glycosylhydrolase  GM418_28815 alpha/beta hydrolase  GM418_28820 SusD/RagB Starch-binding associating with outer membrane  GM418_28825 TonB-linked outer membrane protein, SusC/RagA family  GM418_28830 FAD-dependent oxidoreductase  GM418_28835 sodium transporter | 70%  72%  71%  43%  73%  73% |
| 75 | GM418_28885 glycoside hydrolase family 105 protein  GM418_28890 SusD/RagB Starch-binding associating with outer membrane  GM418_28895 TonB-linked outer membrane protein, SusC/RagA family  GM418_28900 anti-sigma factor  GM418_28905 RNA polymerase sigma-70 factor  GM418_28910 ATP-binding protein  GM418_28915 IS4 family transposase  GM418_28920 family 43 glycosylhydrolase  GM418_28925 lanthionine synthetase  GM418_28930 glycoside hydrolase family 127 protein  GM418_28935 galactose mutarotase  GM418_28940 family 43 glycosylhydrolase  GM418_28945 sulfatase  GM418_28950 sulfatase-like hydrolase/transferase  GM418_28955 arylsulfatase  GM418_28960 arylsulfatase | 18%  55%  56%  81%  18%  83%  47%  80%  17%  61%  48%  73%  60%  45%  77%  76% |
| 76 | GM418_29000 arylsulfatase  GM418_29005 1,4-beta-xylanase  GM418_29010 glycoside hydrolase family 43 protein  GM418_29015 family 43 glycosylhydrolase  GM418_29020 alpha-L-arabinofuranosidase  GM418_29025 beta-galactosidase  GM418_29030 beta-galactosidase  GM418_29035 glycoside hydrolase family 2  GM418_29040 glycoside hydrolase family 2  GM418_29045 glycosyl hydrolases family 2  GM418_29050 glycoside hydrolase family 2  GM418_29055 DUF3823 domain-containing protein  GM418_29060 SusD/RagB Starch-binding associating with outer membrane  GM418_29065 TonB-linked outer membrane protein, SusC/RagA family  GM418_29070 anti-sigma factor  GM418_29075 RNA polymerase sigma-70 factor | 60%  76%  84%  74%  80%  20%  21%  67%  60%  21%  56%  50%  57%  55%  44%  77% |
| 77 | GM418_29920 acetylhydrolase  GM418_29925 glycoside hydrolase family 127 protein  GM418_29930 sulfatase  GM418_29935 hypothetical protein  GM418_29940 bifunctional 2-methylcitrate dehydratase/aconitate hydratase  GM418_29945 argininosuccinate lyase  GM418_29950 SusD/RagB Starch-binding associating with outer membrane  GM418_29955 TonB-linked outer membrane protein, SusC/RagA family  GM418_29960 anti-sigma factor  GM418_29965 RNA polymerase sigma-70 factor | 41%  18%  51%  57%  19%  24%  50%  55%  52%  19% |
| 78 | GM418_30275 MFS transporter  GM418_30280 glycoside hydrolase family 92 protein  GM418_30285 SusD/RagB Starch-binding associating with outer membrane  GM418_30290 TonB-linked outer membrane protein, SusC/RagA family  GM418_30295 anti-sigma factor  GM418_30300 RNA polymerase sigma-70 factor | 66%  36%  71%  57%  33%  28% |
| 79 | GM418_30445 hydrolase  GM418_30450 sulfatase-like hydrolase/transferase  GM418_30455 exo-alpha-sialidase  GM418_30460 SusD/RagB Starch-binding associating with outer membrane  GM418_30465 TonB-linked outer membrane protein, SusC/RagA family  GM418_30470 LacI family transcriptional regulator  GM418_30475 TRAP transporter small permease  GM418_30480 2,3-diketo-L-gulonate transporter large permease YiaN  GM418_30485 uroporphyrinogen decarboxylase  GM418_30490 DctP family TRAP transporter solute-binding subunit  GM418_30495 2-keto-3-deoxy-L-rhamnonate aldolase  GM418_30500 YhcH/YjgK/YiaL family protein  GM418_30505 RraA family protein  GM418_30510 gluconate dehydratase  GM418_30515 D-glycerate dehydrogenase  GM418_30520 galactonate dehydratase  GM418_30525 sugar kinase | 46%  52%  38%  78%  71%  37%  33%  36%  17%  31%  26%  41%  79%  85%  40%  53%  73% |
| 80 | GM418_30990 S41 family peptidase  GM418_30995 HAMP domain-containing histidine kinase  GM418_31000 phenol degradation protein meta  GM418_31005 sulfatase  GM418_31010 glycoside hydrolase family 127 protein  GM418_31015 Sulfatase  GM418_31020 Glycosyl transferase family 2  GM418_31025 SusD/RagB Starch-binding associating with outer membrane  GM418_31030 TonB-linked outer membrane protein, SusC/RagA family  GM418_31035 anti-sigma factor  GM418_31040 RNA polymerase sigma-70 factor | 75%  47%  14%  74%  19%  68%  22%  64%  61%  61%  68% |
| 81 | GM418_31180 transketolase  GM418_31185 alginate lyase  GM418_31190 glycoside hydrolase family 28 protein  GM418_31195 sulfatase  GM418_31200 acylhydrolase  GM418_31205 SusD/RagB Starch-binding associating with outer membrane  GM418_31210 TonB-linked outer membrane protein, SusC/RagA family  GM418_31215 anti-sigma factor  GM418_31220 RNA polymerase sigma-70 factor | 80%  20%  17%  42%  18%  28%  38%  46%  23% |
| 82 | GM418_31235 SusD/RagB Starch-binding associating with outer membrane  GM418_31240 TonB-linked outer membrane protein, SusC/RagA family  GM418_31245 anti-sigma factor  GM418_31250 RNA polymerase sigma-70 factor  GM418_31255 methane monooxygenase PmoA-like  GM418_31260 uroporphyrinogen decarboxylase  GM418_31265 MFS transporter  GM418_31270 dehydrogenase  GM418_31275 substrate-binding domain-containing protein  GM418_31280 D-mannonate oxidoreductase  GM418_31285 Gfo/Idh/MocA family oxidoreductase  GM418_31290 sugar kinase  GM418_31295 D-mannonate epimerase  GM418_31300 Gfo/Idh/MocA family oxidoreductase | 67%  42%  60%  21%  42%  52%  46%  68%  73%  82%  78%  75%  84%  77% |
